# Spatial isoform sequencing at sub-micrometer single-cell resolution reveals novel patterns of spatial isoform variability in brain cell types

**DOI:** 10.1101/2025.06.25.661563

**Authors:** Lieke Michielsen, Andrey D. Prjibelski, Careen Foord, Yelizaveta Spiegelman, Taewoo Kim, Wen Hu, Julien Jarroux, Justine Hsu, Rebecca Pfeil, Xinyi Zhang, Li Gan, Alexandru I. Tomescu, Iman Hajirasouliha, Hagen U. Tilgner

## Abstract

Spatial long-read technologies are becoming increasingly common but lack nanometer and therefore often single-cell resolution. This leaves the question unanswered of whether spatially variable isoforms represent spatial variability within one cell type or differences in cell-type composition between different regions. Here, we developed Spl-ISO-Seq2 (220nm spot size and 500nm resolution), and the accompanying software packages Spl-IsoQuant-2 and Spl-IsoFind, enabling long-read sequencing using up to 500 million barcodes compared to 80,000 previously. Applying this to the adult mouse brain, we compared spatial variability by examining (a) differential isoform abundance between known brain regions and (b) spatial isoform patterns that do not align with predefined regions. While the former revealed more spatial isoform differences, both approaches identified overlapping hits, e.g., *Rps24* in oligodendrocytes. For *Snap25,* previously known to exhibit spatial isoform variation, we now show that this variability occurs in excitatory neurons. The second approach also uncovered patterns not captured by predefined-region comparisons, e.g., *Tnnc1* in excitatory neurons. Furthermore, we show that a surprising number of spatial isoform signals is not driven by cell-type composition alone. Finally, we applied our software to newly generated Visium HD 3’ long-read data to demonstrate its applicability and strong reproducibility across protocols and biological replicates. Taken together, our experimental and analytical methods enrich spatial transcriptomics with a so-far elusive isoform view of spatial variation for individual cell types.

## Introduction

A fundamental question in spatial isoform biology is whether a cell’s spatial position influences its isoform expression. This question can be confounded by different cell types dominating distinct areas in the brain. Therefore, single-cell-resolution spatial isoform data is essential for disentangling spatially distributed isoforms from cell-type-derived ones.

Single-cell and single-nucleus long-read sequencing have enabled profiling isoforms in distinct cell types^1–5^. Short-read sequencing-based approaches have also described cell-type-specific splice junctions^6–8^, revealing exons that are used in specific cell types or altered in evolution and/or disease^5,9,10^. However, a major shortcoming of these methods is that they lose the spatial location of cells, preventing the study of how isoforms may be spatially regulated at the cell-type-specific level.

Simultaneously, spatial gene-expression profiling has moved the field forward by localizing cell types and gene expression across tissue sections^11–18^. Based on 10X Genomics’ Visium technology, we and our colleagues simultaneously engineered spatial isoform sequencing, revealing many isoform switches that correlated tightly with brain structures, and other isoforms (e.g., of *Snap25*) that showed isoform changes within a region^4,19^. However, Visium’s 55μm spot size exceeds the average cell diameter in the mouse brain, and as such, results in “pseudo-bulk” measurements that likely represent multiple cells and thus potentially multiple cell types.

By adapting Slide-SeqV2^12^, we further developed spatial ISOform sequencing (Spl-ISO-Seq) with 10μm resolution and corresponding software Spl-IsoQuant^20^. Spl-ISO-Seq uses exome-sequencing probes^5^ and long-molecule selection^20^ to enrich for spliced and (near-)complete cDNAs. Although 10μm resolution is sufficient to identify large cells in the human brain, like excitatory neurons, it importantly lacks resolution to identify smaller cells. This is especially problematic when considering other common model organisms, like mice.

Since then, new spatial transcriptomic technologies supporting a higher resolution have been released, including the Visium HD 3’ spatial assay^21^ and Stereo-seq^22^. Stereo-seq offers a higher resolution compared to Visium HD (500nm vs 2μm), making it well-suited for single-cell long-read spatial transcriptomics. Although segmentation from ssDNA staining, which labels only nuclei, and ∼10µm z-sampling can still lead to doublets^23^, the substantial gain in resolution is highly promising for spatially studying individual cells.

Therefore, based on the Stereo-seq approach, coupled with platform-specific artifact removal and our previously developed exome enrichment^5^ and long-molecule selection^20^, here we devise Spl-ISO-Seq2 with 20-fold improved spatial resolution (500nm instead of 10μm) for over 100 million barcodes. We show that this spatial isoform technology can work with PacBio (PB) as well as with Oxford Nanopore Technologies (ONT) long reads. To complement this wet lab approach, we devised Spl-IsoQuant-2, which adds highly-specific barcode calling to our IsoQuant algorithm^24^ that we further substantiate through dry-lab simulation experiments. We show that unique molecules sequenced on both PB and ONT reveal high accuracy of transcript assignments. Furthermore, Spl-IsoQuant-2 can be applied to long-read data generated using all single-cell and spatial protocols. It has been tested on long reads derived from 10x 3’ single-cell, 10x Visium, Visium HD 3’, and Curio Biosciences’ Slide-seqV2. Moreover, Spl-IsoQuant-2 supports user-specified molecule structure and can process long-read data generated using any custom protocol.

To detect spatially variable isoforms, we developed Spatial Isoform Finder (Spl-IsoFind). For similar questions regarding gene expression, many analytical frameworks have been developed, e.g. for identifying spatially variable genes^25–32^. However, methods to detect spatially variable isoforms have lagged behind. Spl-IsoFind uses Moran’s I^33^, a global spatial autocorrelation score, which has been successfully used for gene-expression studies^25,34–36^. We identify cell-type-specific spatially variable isoforms, such as *Snap25* for excitatory neurons and *Rps24* for oligodendrocytes, as well as isoforms with spatial patterns that do not align with predefined regions. Furthermore, using Spl-IsoFind’s cell-type-constrained permutations, we show that most spatial isoform signals are not solely due to differences in cell-type composition alone. Finally, we demonstrate Spl-IsoQuant-2 and Spl-IsoFind’s versatility by applying it to Visium HD 3’ long-read mouse brain samples. Despite slight differences in slide positioning and brain region captured, our results show strong reproducibility across protocols and biological replicates.

Overall, Spl-ISO-Seq2, Spl-IsoQuant-2, and Spl-IsoFind together provide a long-read-compatible spatial transcriptomics framework at single-cell resolution. Applied to the adult mouse brain, this approach reveals cell-type-specific isoform changes both between predefined regions and in previously unrecognized spatial patterns. Altogether, this high-resolution technology can be applied to any tissue type to investigate single-cell spatial isoform patterns.

## Results

### Gene-expression patterns in coronal brain slices

To define spatially variable isoforms for a given cell type, we used two consecutive coronal slices of an adult mouse brain, covering cortex, hippocampus, thalamus, and midbrain (**Fig S1a-b**). We used the Stereo-seq spatial approach to barcode cDNAs at 500nm resolution^22^ and performed short-read sequencing as per Stereo-seq guidelines. The same cDNA was used to perform exome enrichment and long-molecule selection, as we have previously done^20^, followed by PB Kinnex and ONT long-read sequencing. Short-read sequencing, in conjunction with an ssDNA-stain, was used to segment individual cells, assign cell types, and anchor barcodes to specific spatial locations (Methods). Long reads, both of the PB and ONT platforms, were used to define novel isoforms and to quantify annotated and novel isoforms (**Fig 1a**). In this study, we focused our analysis on the complete right hemisphere. Short-read gene expression clearly indicated different brain regions, including distinct cortical layers (**Fig 1b, S1c**). We used Robust Cell Type Decomposition (RCTD), a state-of-the-art deconvolution method, to assign cell types to the segmented cells^37–39^. Although Stereo-seq has high resolution, overlapping cells can still cause doublets^23^ (**Fig S2**). We defined our major cell types using RCTD’s singlet spots only (**Fig 1c, S1d, S3**). While excluding doublets reduces the statistical power, especially for smaller cell types such as astrocytes and oligodendrocytes, it provides a more accurate basis for detecting cell-type-specific spatially variable isoforms. Furthermore, staining and short-read data were sufficient to align the two coronal brain slices (**Fig 1d**). Of note, the cortex, and above all, some hippocampal areas stood out in terms of UMI density, owing to their higher cell density (**Fig 1e, S1e**).

**Figure 1.**
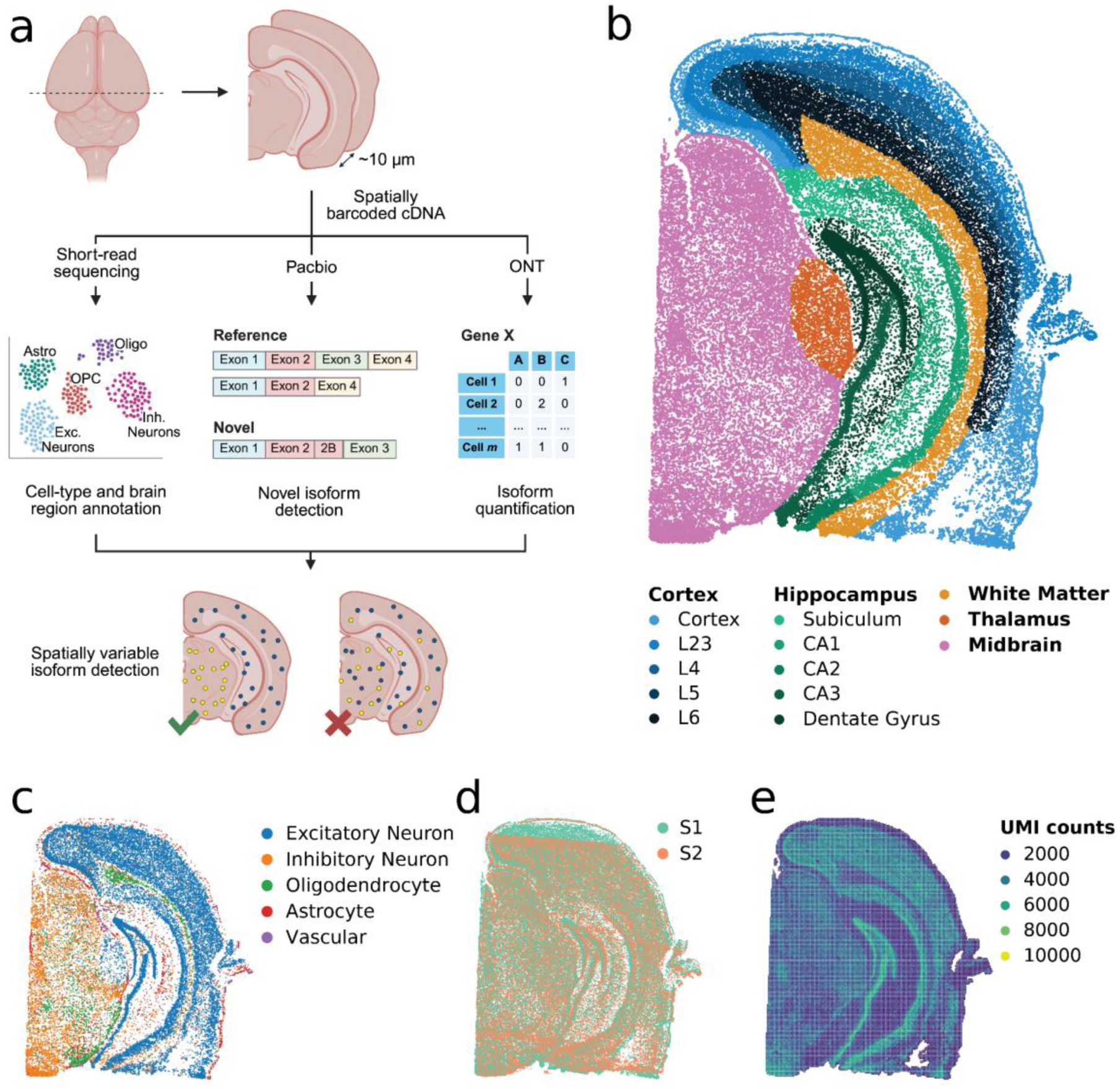
Long-read spatial transcriptomics of two mouse brain slides. a) Schematic overview of the study design. Spatially barcoded cDNA was generated for two coronal sections (10μm apart) of a P56 mouse brain using the Stereo-seq platform^22^. The pool of cDNA was sequenced using 1) short-read technologies, 2) PB, and 3) ONT. All sequencing data were combined to detect (cell-type-specific) spatially variable (novel) isoforms. b) (Sub)region annotation of Sample 1. c) Cell-type annotations of Sample 1. Only cells annotated as singlets and cell types with >100 cells are plotted. d) Alignment of Sample 1 (S1) and Sample 2 (S2). e) UMI counts per bin (50x50spots) for Sample 1.

### A long-read approach for spatial isoform expression with high precision

We engineered a long-read approach for spatial isoform expression using the spatially barcoded, exome-enriched cDNAs after long-molecule selection (Methods). The Stereo-seq protocol, due to the usage of similar primers at 3’ and 5’ ends, can allow for the concatenation of multiple barcoded cDNAs. To counter this effect, we used a PCR strategy to append an additional sequence to the barcoded end of the read (Methods). Although this strategy decreased the rate of molecule concatenation, it did not fully eliminate it. This concatenation affects the cDNA itself, and is unrelated to the combination of reads involved in PB library preparation, which can easily be split with existing software (Methods). To account for cDNA read concatenations, Spl-IsoQuant-2 first implements a read ‘splitting’ strategy to split concatenated molecules followed by a barcode recognition algorithm (Methods). Consequently, extracted cDNAs are assigned to individual genes and isoforms, and PCR duplicates are removed (**Fig 2a**, Methods). To account for possible previously unannotated isoforms we first ran IsoQuant^24^ using all sequenced data to generate an annotation containing 28,891 known and 26,445 novel isoforms, which were then used as input to the Spl-IsoQuant-2 pipeline. To generate this annotation, Gencode version 36 mouse gene annotation^40^ was used (but other annotations, including RefSeq^41^ or Ensembl^42^, can be used as well). As a core pipeline integrated into Spl-IsoQuant-2, IsoQuant has demonstrated high precision in novel transcript discovery by previous reports^24,43,44^.

**Figure 2.**
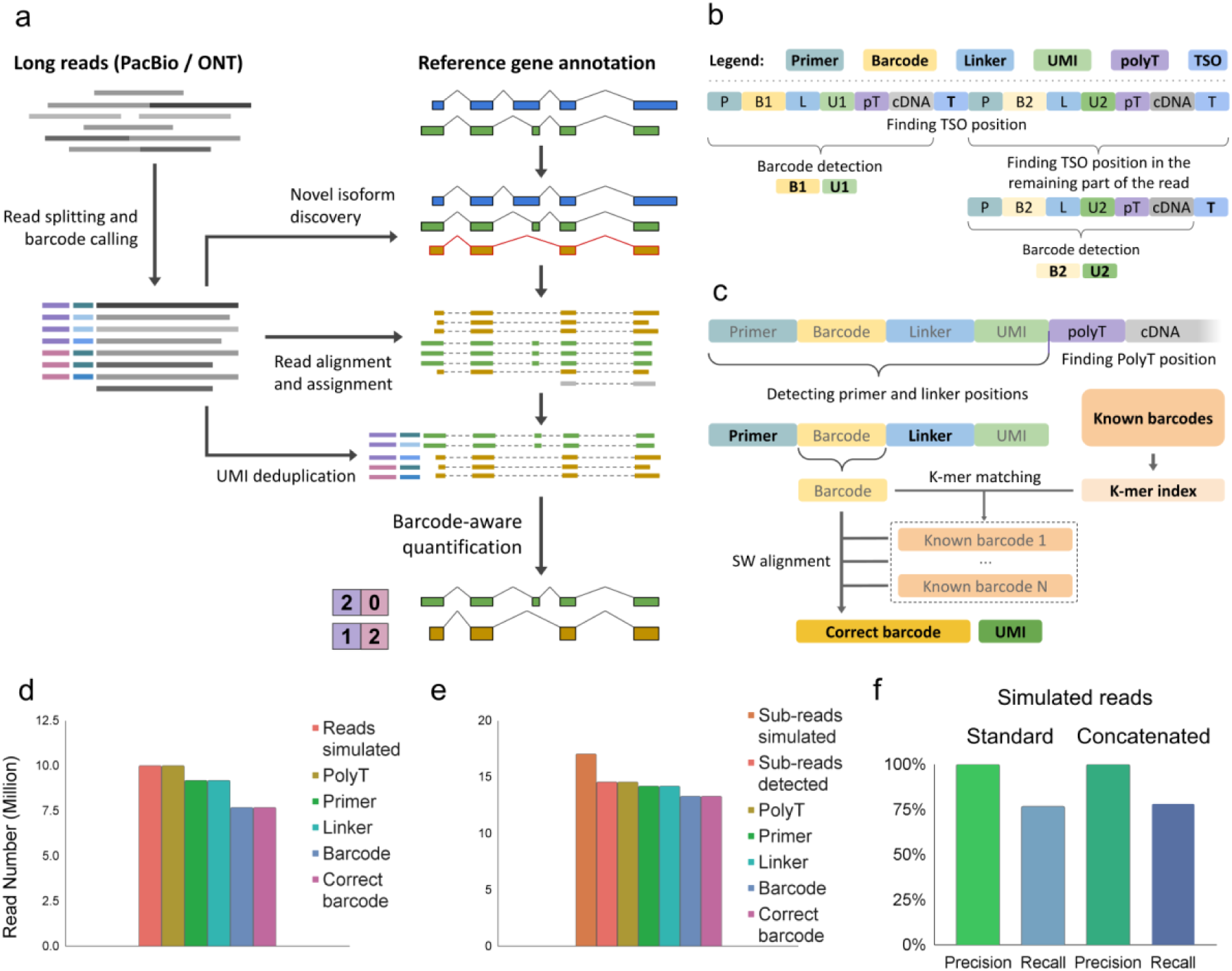
Processing long reads obtained with Spl-ISO-Seq2. a) Spl-IsoQuant-2 pipeline overview. Purple and pink bars next to the reads indicate different barcodes. Blue bars between the reads and the barcodes indicate UMIs. Final diagram shows isoform count per barcode (purple/pink). b) Splitting reads into individual cDNAs by locating the template-switch oligos (TSO). Elements are denoted as follows: P (primer), B (barcode), L (linker), U (UMI), pT (polyT stretch), T (TSO). c) Barcode detection using k-mer indexing and Smith-Waterman local alignment. d) Primer, linker and barcode detection statistics for simulated reads without cDNA concatenation. e) Primer, linker and barcode detection statistics for simulated reads with cDNA concatenation. A simulated read can consist of multiple concatenated subreads resulting in >16 million subreads in total. f) Precision and recall of the designed barcode calling algorithm for simulated data without (top) and with cDNA concatenation (bottom). g) Agreement between real PB and ONT reads.

Detection of multiple concatenated isoforms is performed by locating the template-switch oligo (TSO) as well as the linker and barcode sequences far from read extremities (**Fig 2b**). To enable downstream analysis of long-read Stereo-seq data, Spl-IsoQuant-2 corrects read barcodes by finding the closest candidate from the whitelist using a k-mer index and local alignment (**Fig 2c**) (see Methods for details). Our implementation indexes about half a billion Stereo-seq 25bp barcodes in less than 10 minutes using 16 threads. Barcode correction allows processing roughly 60 million ONT reads per hour without deconcatenation and 10 million reads per hour with deconcatenation in 16 threads, making it applicable even for large datasets. Although various software for barcode detection in ONT reads exists at the moment^45–47^, none of these tools are capable of “parsing” Stereo-seq’s molecule structure. Even when using Flexiplex, arguably the most versatile tool, we were able to process only 1 million ONT reads in 5 days (Methods).

To assess recall and precision, we first simulated reads with a 3.8% error rate without simulating read concatenation (Methods). A polyT sequence, representing the polyA tail, could be found in virtually all reads, however primer and linker were detected in only 91.78% of the reads, owing to simulated sequencing reads. Spl-IsoQuant-2 detected a barcode in 76.66% of reads and correctly called the known simulated barcode in 76.62% of reads (**Fig 2d**), thus yielding a precision of 99.95% and a recall of 76.65% (**Fig 2f, top panel**). Similar simulation of sequencing errors along with cDNA concatenation yielded a precision of 99.94% and a recall of 78.11% (**Fig 2e,f**). In summary, Spl-IsoQuant-2 yields close to perfect precision, while delivering high recall for barcode recognition.

Besides Stereo-seq, Spl-IsoQuant-2 also supports various protocols, such as 10x v2/v3 3’ single-cell, 10x Visium 3’, 10x Visium HD and Curio Biosciences’ Slide-seqV2 spatial protocol. Furthermore, it implements a molecule description format that allows a user to set their own molecule structure and perform barcode calling for any custom protocol (Methods).

We then enriched barcoded cDNA using exome-sequencing probes^4,5,9,20^ and long-molecule selection^20^. We sequenced 203.7 and 58.8 million reads on the ONT platform for the two coronal slices (from now on called Sample 1 AllExome (AE) and Sample 2 (AE) respectively) as well as 19.9 million and 17.6 million PB reads. For Sample 1 (AE), we used two ONT flow cells instead of one, explaining the difference in read depth. We also performed one ONT long-read run on cDNAs enriched for 3378 genes related to various brain diseases and synaptic function (from now on called Sample 1 (3.3K)), rather than for the whole exome (**Table S1-8**) (Methods). Gene expression measured using the short-read and long-read sequencing technologies is strongly correlated (**Fig S4**). We compared the number of sequenced reads required to obtain a given number of informative reads (spliced reads with an assigned isoform that overlap a segmented cell in the correct hemisphere) to a similar long-read spatial transcriptomics dataset but generated on the Visium platform^19^ (Methods). Both platforms show a comparable sensitivity as a similar number of sequenced reads (∼200M) yields a comparable number of informative reads (∼8M) (**Table S9**). Importantly, Spl-ISO-Seq2 additionally provides single-cell resolution.

Of note, due to the amplification and splitting of cDNA libraries for distinct sequencing protocols, cDNA copies of the same original molecule can be sequenced on both platforms. We therefore tested reproducibility and the influence of higher error rates in ONT sequencing on transcript assignments by taking advantage of such pairs of reads sequenced on both platforms. Indeed, 3.5 and 2.3 million unique molecules for Sample 1 (AE) and Sample 2 (AE), respectively, were sequenced on both ONT and PB (Methods). In both replicates, 99.4% of such read pairs were assigned to the same isoform (**Fig 2g**). This observation supports that errors in ONT reads do not lead to widespread mis-assignments of reads to the wrong isoform.

### Detecting spatially variable isoforms using predefined brain regions

We compared differential relative isoform expression (not confounded by gene expression) between brain region pairs on ONT combined Sample 1+2 (AE) using our *n* x 2 isoform tests^4,9^. For each gene, we built an *n* × 2 table of isoform counts and applied a chi-squared test with Benjamini-Hochberg (BH) correction (Methods). The midbrain showed the most differentially used isoforms, especially when compared to hippocampus and cortex (**Fig 3a, Table S10-14**). Interestingly, while the percentage of significant genes was similar between the midbrain and thalamus, the total number of significant genes in the thalamus was much lower, which is explained by the lower read depth. A similar pattern appears in two previously published coronal brain sections^19^ (see Methods) (**Fig S5, Table S15**). However, in these data, we also detected many differences involving the white matter, which reflects imperfections in harmonizing regional annotations between studies.

**Figure 3.**
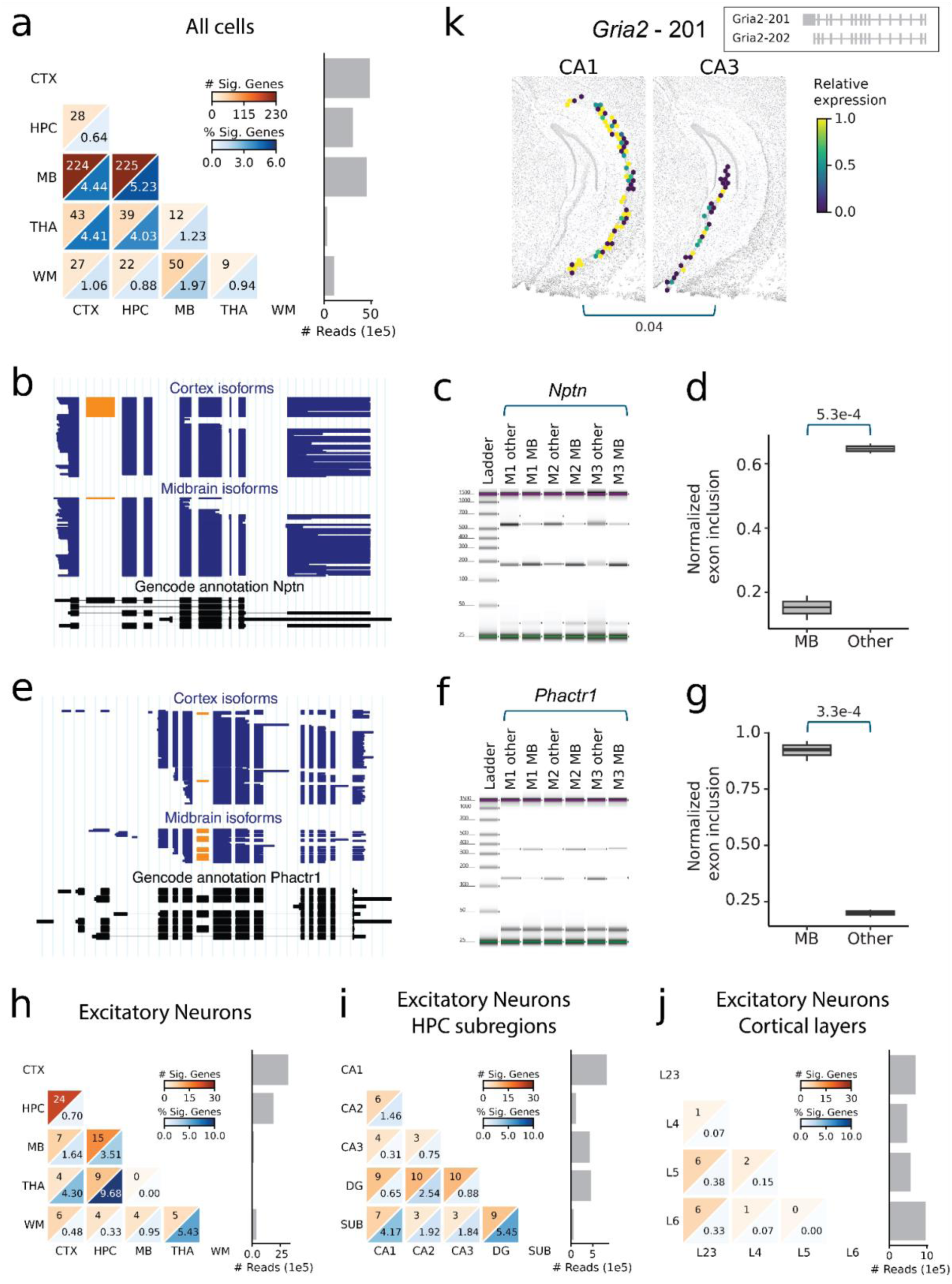
Detecting genes exhibiting differential isoform expression across predefined regions. a) Number and percentage of genes changing isoform expression across predefined brain regions using all cells in combined Sample1+2 (AE). Barplot on the right shows the number of long reads used during the analysis per brain region. b,e) Single-cell long-reads for b) *Nptn* and e) *Phactr1* measured in the cortex and midbrain. Each line in the top two tracks represents a single cDNA molecule. The orange exon drives the difference between isoforms in the two brain regions. The bottom black track shows the Gencode annotation b) (chr9: 58489523-58565238) and e) (chr13: 42834099-43292002). c,f) Tapestation gel of PCR amplified products using exon-of-interest-spanning c) *Nptn* and f) *Phactr1* specific primers. d,g) Normalized quantification of tapestation gel for d) *Nptn* and g) *Phactr1* (Methods). Welch’s two-sample t-test was used for statistical testing. h-j) Number and percentage of genes changing isoform expression in excitatory neurons across (sub)regions. k) Relative expression of *Gria2*-201 in excitatory neurons in different hippocampal subregions of Sample 1 (AE). Every hexagon shows the mean relative expression of the underlying cells. Relative expression indicates the fraction of long reads for that gene (*Gria2*) that are assigned to our isoform of interest (201). A hexagon is only plotted when there are underlying cells expressing *Gria2* (Methods). As such, a yellow spot means that *Gria2*-201 is expressed, while a darkblue spot means that *Gria2*-201 is not expressed at that location, but another *Gria2* isoform is expressed. The ssDNA staining is shown in the background. Numbers indicate BY-corrected p-values. The schematic diagram in the corner shows the gene structure of *Gria2*-201 and *Gria2-*202, the two most common isoforms in the data. CTX = cortex, HPC = hippocampus, MB = midbrain, THA = thalamus, WM = white matter, CA = Cornu Ammonis, DG = Dentate Gyrus, SUB = Subiculum, Mx = mouse sample, N.S. = not significant

Two example genes differentially spliced between the midbrain and cortex are Neuroplastin (*Nptn*) and Phosphatase and Actin Regulator 1 (*Phactr1*) (BH-corrected p-value = 0.017 and 0.0067, respectively). For both genes, one exon drives the difference between the cortex and midbrain so we used PCR to validate these exons’ spatial inclusion (**Fig 3b-g, S6**) (Methods). For *Nptn*, normalized exon inclusion was decreased in the midbrain compared to other regions (p<5.4e-4, Welch’s two-sided two-sample t-test). Of note, *Nptn* encodes a synaptic glycoprotein involved in synaptogenesis, long-term potentiation and memory formation^48^. These two *Nptn* isoforms have been documented by the field as Np55 and Np65, where Np65 contains an additional immunoglobulin domain^49^. For *Phactr1*, which helps to regulate actin cytoskeletal organization^50,51^, normalized exon inclusion increased in the midbrain (p<3.4e-4, Welch’s two-sided two-sample t-test).

We initially expected differences in cell-type composition to drive most significant results. However, restricting the analysis to excitatory neurons alone resulted in a similar percentage of significant isoforms for each comparison (**Fig 3h**). For each comparison, except for cortex vs. midbrain and thalamus vs. white matter, the confidence intervals for all cells and excitatory neurons overlap (**Fig S7**), indicating that cell-type composition alone is unlikely to explain results from all cells.

Within the hippocampus, especially the CA3 subregion compared to the dentate gyrus (DG) showed differential isoform abundance (**Fig 3i**). Likewise, in the cortex, layer 2-3 showed important changes in isoform regulation especially when compared to layer 5 and layer 6 (**Fig 3j**). For some genes with layer-specific splicing changes, such as *Nnat* and *Arpp21*, the relative isoform abundance shows a gradient across the layers, while for others, such as *Dtnbp1* and *Tpm1*, one isoform seems to be more specific to one layer (**Fig S8**). Since excitatory neuron subtypes in the cortex are strongly layer-enriched, laminar isoform patterns may reflect subtype-specific isoform regulation. However, these subtypes are enriched rather than strictly confined to individual layers, and individual layers might be enriched for multiple subtypes as well. Interestingly, *Gria2* shows isoform expression changes between excitatory neurons in the hippocampus and cortex. Isoform 201 is more frequently used in the hippocampus compared to the cortex. However, examining these broader brain regions does not capture all nuances: within the hippocampus, differences are also evident between the CA3 and CA1 subregions (**Fig 3k, S9**). Similarly, in the cortex, notable isoform changes are observed between layers 2-3 and layer 5 (**Fig S10**). *Gria2* encodes the GluA2 protein, one of 4 subunits which constructs the glutamatergic α-amino-3-hydroxy-5-methyl-4-isoxazolepropionic (AMPA) receptor. The two spatially variable *Gria2* isoforms identified here represent the *flip* and *flop* variants, a known alternative-splicing driven variance which alters the channel’s rate of desensitization^52–54^. In the other major cell types (inhibitory neurons, oligodendrocytes, and astrocytes), we identified fewer significant genes compared to excitatory neurons, which is mainly explained by a lower read depth (**Fig S11**).

While the previous analysis focused on testing differentially used isoforms, we also tested for alternatively spliced cassette exons (Methods). During these tests, the number of significant results decreases (**Fig S12**), even though a higher number of reads were used during the tests. This decrease is likely due to the isoform tests indirectly capturing a broader range of variations, such as alternative 3’ and 5’ splice sites, transcription start sites, and polyadenylation sites, which are not considered in the exon-specific tests.

### Spl-IsoFind identifies spatially variable isoforms beyond predefined brain regions

Previously, we compared isoform usage between predefined regions, assuming that the underlying alternative splicing mechanisms respect these borders. In theory, isoform usage might differ within a region or even across regions (**Fig 4a**). Therefore, we developed Spatial Isoform Finder (Spl-IsoFind) to calculate spatial autocorrelation using Moran’s I^33^ (Methods). Moran’s I indicates whether an isoform exhibits a spatial pattern by comparing the values of neighboring cells. This method has been used to detect spatially variable genes^25,34–36^, but not yet spatially variable isoforms (SVI). Since we compare relative isoform usage, the latter is slightly more challenging since many cells have missing values as the gene was not measured in the long-read data.

**Figure 4.**
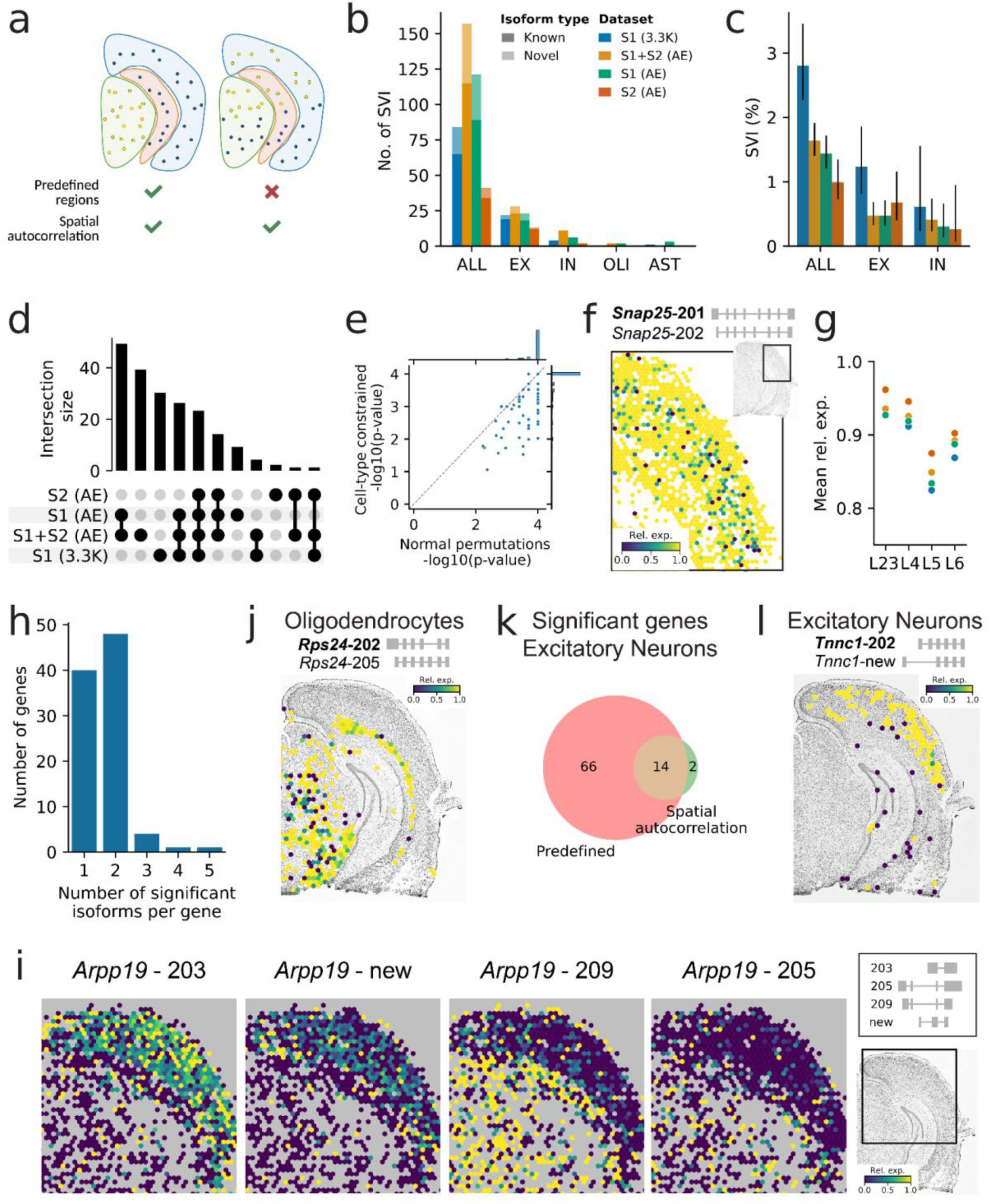
Detecting spatially variable isoforms using spatial autocorrelation. a) Schematic example of a pattern (left) which can be detected using either predefined regions and spatial autocorrelation and a pattern (right) which can only be detected using spatial autocorrelation. b) Number and c) percentage of significant spatially variable isoforms (FDR < 0.05) per dataset and celltype. Error bars indicate 95% confidence interval. d) The overlap of significant isoforms across the datasets. Note: not all isoforms were tested in all datasets due to differences in read depth. e) Comparing the uncorrected p-values obtained using normal and cell-type-constrained permutations for the isoforms significant in Sample 1+2 (AE) using all cells. f) Spatial distribution of *Snap25*-201 across the cortical layers in the Sample 1 (AE). Every hexagon shows the mean relative expression of the underlying cells (Methods). The schematic gene diagram shows the structure of the two *Snap25* isoforms in our data. g) Mean relative expression of *Snap25*-201 in different cortical layers. Colors indicate the different datasets. Lower relative expression of *Snap25*-201 in Layer 5 implies higher relative expression of *Snap25*-202. h) Number of significant isoforms per gene in Sample 1+2 (AE). i) Spatial distribution of the four significant isoforms of *Arpp19* in Sample 1 (AE). Every hexagon shows the mean relative expression of the underlying cells (Methods). The schematic gene diagram shows the structure of the isoforms. j) Spatial distribution of *Rps24*-202 in oligodendrocytes of Sample 1 (AE). Every hexagon shows the mean relative expression of the underlying cells (Methods). The gene diagram on top shows the structure of the two most common *Rps24* isoforms in the data. k) Overlap between the significant genes found using both methods for the excitatory neurons in the Sample 1 (AE) + Sample 2 (AE) data. l) Spatial distribution of *Tnnc1*-202 in excitatory neurons of Sample 1 (AE) which was only discovered using spatial autocorrelation. Every hexagon shows the mean relative expression of the underlying cells (Methods). The gene diagram on top shows the structure of the two most common *Tnnc1* isoforms in the data. SVI = spatially variable isoforms, ALL = all cells, EX = excitatory neurons, IN = inhibitory neurons, OLI = oligodendrocytes, AST = astrocytes, AE = AllExome dataset, 3.3K = 3.3K targeted dataset.

First, we assessed Spl-IsoFind’s precision and recall using simulated data (Methods). We converted Moran’s I score to a p-value using permutation tests and adjusted for multiple testing using Benjamini-Yekutieli correction. In simulated random data without a spatial pattern, the false discovery rate is low (0.17%) (**Fig S13**). Only the smallest simulated dataset (50 cells) has some false positives (5 out of 165 simulations). The recall, however, greatly depends on the underlying simulated pattern. We detect distinct patterns, such as a border between two areas, during all simulations, regardless of the number of cells. For complex patterns (e.g., a small area of interest which is surrounded by cells with random values), the result depends on the number of cells in our simulation.

Next, we ran Spl-IsoFind on all ONT datasets in a cell-type-agnostic and cell-type-specific fashion (**Fig 4bc, Table S16-20**). In the combined Sample1+2 (AE) dataset, we detected the most SVIs (157 SVIs corresponding to 94 genes). Gene ontology (GO) enrichment analysis revealed that these genes are enriched in synaptic vesicle trafficking, synaptic signaling, and membrane excitability (**Fig S14**). The cell-type-agnostic mode resulted in more SVIs compared to the cell-type-specific mode for two reasons: 1) we use more cells during testing, and 2) isoforms can be cell-type-specific and the distribution of cell types can change across regions. As shown with the simulations, having more cells and thus more reads results in more power to detect a pattern. Sample 2 (AE) has the lowest number of reads per isoform (**Table S7**) and as such the lowest number of detected SVIs. The higher percentage of detected isoforms in Sample 1 (3.3K) is most likely due to our enrichment for neuronal genes. In general, the Moran’s I score is very correlated to Geary’s C^55^, another autocorrelation metric more sensitive to local variations, which discovered almost the exact same set of SVIs for combined Sample1+2 (AE) (**Fig S15**).

Most SVIs (59.6%) are detected in two or more datasets (**Fig 4d**). However, 152 isoforms (386 pairwise combinations) are tested in two (or more) datasets, but significant in only one. In 348 cases (90.2%) the isoform is significant in the dataset with the higher read depth (**Fig S16**). Both combined Sample1+2 (AE) and Sample 1 (3.3K) have many unique SVIs. The former highlights the benefit of a higher read depth in general, while the latter demonstrates that targeted sequencing leads to a higher read depth for our genes of interest, leading to many new discoveries. Besides strong agreement between the Spl-ISO-Seq2 samples, SVIs detected in the combined Sample1+2 (AE) data also greatly overlap with SVIs we discovered in the two previously published coronal brain sections^19^ (**Fig S17**).

To better assess the effect of read depth, we performed a downsampling analysis (Methods) (**Fig S18**). While deduplicated and informative reads do not seem fully saturated, the number of SVIs detected in all cells starts to saturate at ∼100M sequenced reads for Sample1 (AE). In contrast, cell-type-specific SVIs do not show clear saturation, suggesting that additional sequencing depth would increase detection power. Combining consecutive slides, as in the combined Sample1+2 (AE) dataset, increases the number of detected SVIs further, even for the cell-type-agnostic SVIs.

In previous work, we proposed three models to explain alternative splicing across brain regions: 1) multiple cell types change splicing, 2) one cell type changes splicing, 3) no cell type directly changes splicing, but the cell type composition changes^4^. Cell-type-agnostic SVIs can fall into either the first or third category. To distinguish between these, we calculated p-values using cell-type-constrained permutations (Methods). Unlike our previous approach, where the values of all cells were shuffled randomly to convert Moran’s I to a p-value, we now shuffle values only within the same cell type (**Fig S19**). For Sample 1+2 (AE), the uncorrected p-values of most isoforms remain very similar (**Fig 4e**). After applying multiple testing corrections, 146 out of the 157 isoforms remain significant (FDR < 0.05, BY corrected), suggesting that at least one cell type exhibits spatial variability. Thus, most isoform-level spatial signals cannot be explained by shifts in cell-type composition alone.

An example of significant SVIs after applying cell-type-constrained permutations are *Snap25*-201 and *Snap25*-202. These isoforms exhibit a spatial pattern across cortical layers and in the midbrain (**Fig 4f-g, S20**). In general, *Snap25*-201 is the dominant isoform, except for parts of the midbrain and the deeper cortical layers which also show expression of *Snap25*-202. Previous studies have identified similar spatial patterns in the brain, but their resolution was insufficient to pinpoint individual cell types^4,19^. With our cellular resolution, we can analyze these patterns at the cell type level. Notably, we observe significance in excitatory neurons (where the pattern is present in the cortex) and inhibitory neurons (where the pattern appears in the midbrain) (**Fig S20**). These inhibitory neurons may be dopaminergic, though potentially mislabeled, and the region in the midbrain exhibiting the inclusion of *Snap25*-202 is likely the substantia nigra or VTA.

Besides region- and cell-type-specific splicing, *Snap25* exhibits sex- and development-specific splicing differences^56^. While *Snap25*-202 is the main isoform during early development, *Snap25*-201 is during adulthood. In the mouse hippocampus, this switch occurs later in females compared to males. As sex-specific splicing occurs in more genes^57,58^, our discovered SVIs in the male mouse brain might not translate to females. Using human single-nuclei isoform RNA sequencing (SnISOr-Seq) data consisting of three female and three male hippocampal samples^9^, we identified 229 exons with sex-specific inclusion differences, the majority of which were detected in excitatory neurons (154 exons) (**Fig S21a**). One gene, *PPP3CA*, showed both sex-specific splicing differences in human and has two SVIs in our mouse data (**Fig S21b-c**). The same exon, which is conserved between human and mouse, distinguished the SVIs and was differentially included between males and females. *PPP3CA* regulates synaptic and transcriptional signaling through presynaptic and postsynaptic dephosphorylation events and NFAT-mediated transcription, and plays a role in brain circuits involved in reward, memory, and aging^59,60^. The sex-specific exon, which has higher inclusion in females, does not alter known protein domains but has been reported to modulate protein activity in differentiated myotubes^61^. This exon shows spatial variation within the cortex and the hippocampus, with the biggest observed difference between the dentate gyrus (DG) and CA regions (**Fig S21e)**. In the single-nucleus data, exon inclusion is higher in excitatory neurons in both brain regions in females (**Fig S21d)**. *SNAP25* did not show any sex-specific exon inclusion. All human donors were 28-40 years old, and as such, the developmental switch has happened in both females and males already.

*Snap25* serves as a classic example of two isoforms switching between different conditions. However, for some genes, we observe more complex patterns, with either only one or more than two significant isoforms (**Fig 4h**). For instance, *Arpp19* exhibits four distinct SVIs. Two isoforms (203 and a newly discovered isoform) are specific to the cortex and mainly expressed in neurons. *Arpp19*-209 is highly expressed across all brain regions except the cortex, while *Arpp19*-205 shows the lowest overall expression but is predominantly used in the hippocampus, midbrain, and thalamus (**Fig 4i, S22-23**). *Arpp19*-205 and *Arpp19*-209 both encode ARPP19, whereas *Arpp19*-203 and *Arpp19*-new encode ARPP16. ARPP16 and ARPP19 differentially regulate protein phosphatase 2A (PP2A) activity. ARPP19 has established roles in cell-cycle regulation and phosphatase inhibition, whereas ARPP16 appears more enriched in neural tissues and has been implicated in brain-specific PP2A modulation^62^. In addition, the two ARPP19 isoforms (*Arpp19*-205 and *Arpp19*-209) differ in their 3′ UTR length, suggesting potential differences in post-transcriptional regulation, such as mRNA stability or translational control, which is a common regulatory mechanism in mammalian brains^63^.

We identified two significant isoforms of *Rps24* across all cell types except astrocytes. For example, in oligodendrocytes, we observed a distinct spatial difference between the midbrain and white matter (**Fig 4j** **S24**). Isoform *Rps24*-202 was predominantly expressed in the white matter, while the midbrain exhibited a more balanced expression of *Rps24*-202 and *Rps24*-205.

We then compared our SVIs to the genes we identified using the predefined regions. This comparison is challenging, as Spl-IsoFind directly pinpoints isoforms, whereas using the predefined regions, we identified genes with splicing changes, but it was not always clear which specific isoforms were affected. In general, using the predefined regions, we identify more significant genes (**Fig. 4k, S25**), as fewer reads are typically required to detect significance. However, by applying Spl-IsoFind, we also uncover new SVIs, such as *Tnnc1*-202 and a newly discovered *Tnnc1* isoform (**Fig. 4l, S26**). *Tnnc1*-202, the canonical transcript, encodes a 161 amino acid (aa) protein containing four EF-hand motifs, three of which are functional Ca²⁺-binding domains^64^. The newly identified isoform is predicted to encode a truncated 117 aa protein lacking the N-terminal region which includes the first EF-hand motif. The remaining EF-hand domains are preserved, suggesting that the protein may retain partial Ca²⁺-binding capacity. While the precise role of *Tnnc1* in the brain remains unclear, this truncation may alter calcium sensitivity or regulatory interactions.

During all experiments, we initially ran Spl-IsoFind using 10 nearest neighbors (*k*=10) as this emphasizes the direct neighborhood of a cell. We evaluated the effect of this parameter by using a range of neighborhood sizes on the combined Sample1+2 (AE) data. The optimal number of *k* differs per cell type: for all cells combined and excitatory neurons, most SVIs were detected using *k*=50, while for inhibitory neurons and oligodendrocytes, *k*=100 seems most optimal (**Fig S27a**). Most importantly, for most cell types, any number between *k*=10 and *k*=200 yielded broadly consistent results, with substantial overlap between the significant hits (**Fig S27b**). Specifically, 78 of 157 SVIs (49.7%) detected using *k*=10 were detected using all *k* between 10 and 200. Among the remaining 79 SVIs, 65 could not be tested using all *k* because of insufficient read depth. Although the selected neighborhood size (*k*=10) may be sub-optimal, it was maintained without retrospective adjustment to avoid overfitting.

### Spatially variable isoforms strongly replicate across protocols and biological samples

To showcase that our software is generalizable to other long-read spatial transcriptomics datasets, we applied Spl-IsoQuant-2 and Spl-IsoFind to three Visium HD 3’ long-read mouse brain datasets: V1, V2, and V3. V1 is sequenced using both ONT and PB after enriching the barcoded cDNA using exome enrichment and long-molecule selection. V2 and V3 are two publicly available datasets sequenced using ONT and PB, respectively. For V2 and V3, we analyzed 8μm bins instead of individual cells, as a high-resolution image required for cell segmentation was unavailable.

While Visium HD has 2μm resolution, the very similar barcodes of neighboring spots make it challenging to assign reads to the exact location, especially for ONT data, considering the error rate. Using simulated data with an ONT error profile, we compared the barcode detection performance of Spl-IsoQuant-2 and Space Ranger, the 10x Genomics’ software developed to analyze short-read Visium data. Spl-IsoQuant-2 has a higher precision while maintaining a comparable recall to Space Ranger (**Fig 5a**). For Spl-IsoQuant-2, using a minimal alignment score of 13 for each of the two barcode sequences (see Methods) is the best trade-off between precision and recall. Although grouping the individual spots in 8 or 16μm bins greatly improved precision and recall, we preferred to maintain a 2μm resolution for V1 and group spots afterwards based on our cell segmentation. For V2 and V3, we used the 8μm bins since this increased precision.

**Figure 5.**
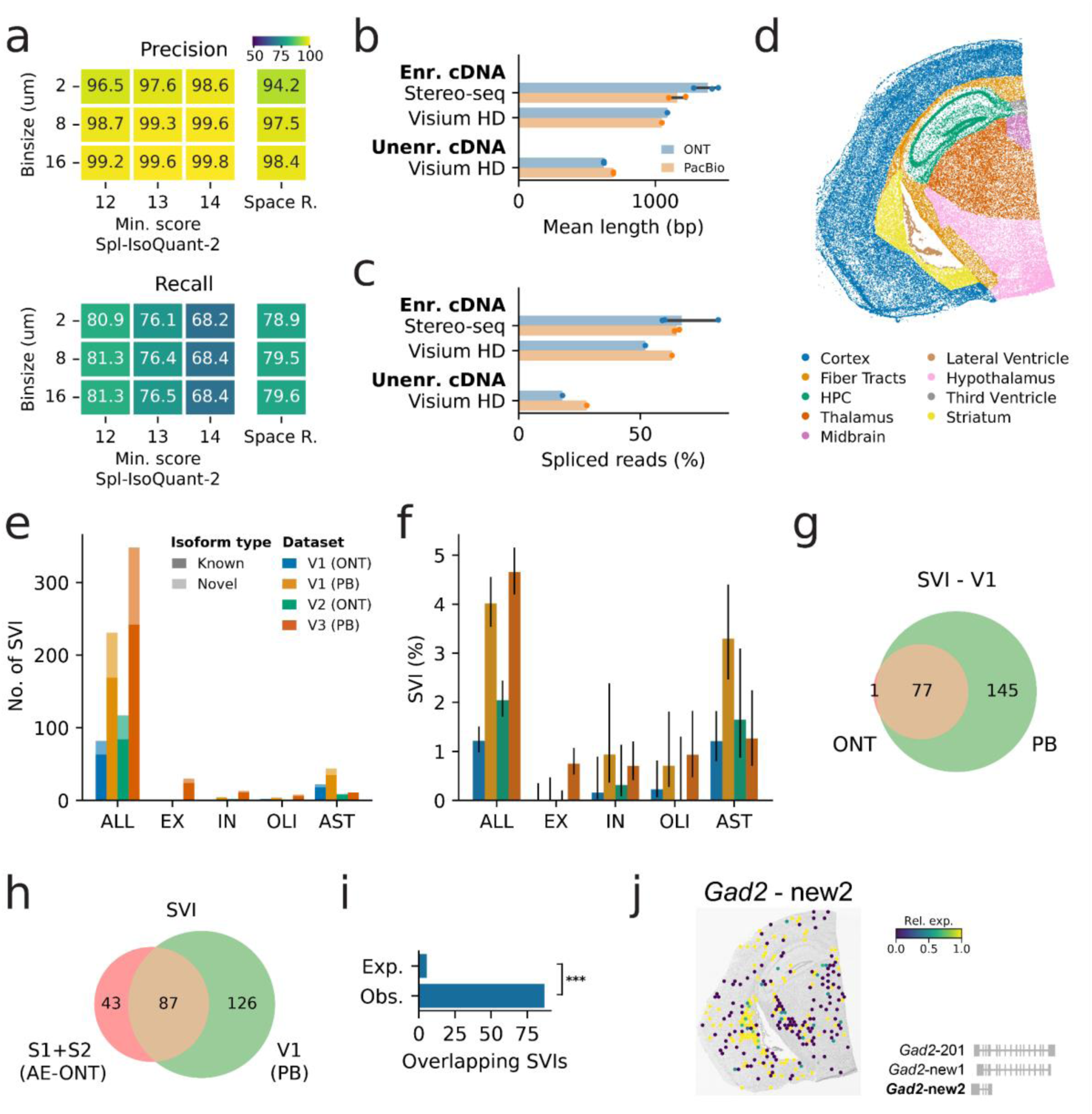
a) Barcode calling precision and recall of Spl-IsoQuant-2 and Space Ranger on simulated Visium HD data using different while grouping the spots in different bin sizes. Different minimal alignment scores (min. score) were tested for Spl-IsoQuant-2. **b)** Average read length and **c)** percentage of spliced reads in the Stereo-seq and Visium HD data. Datasets are grouped by whether enriched (enr.) (long-molecule selection and exome enrichment) or unenriched (unenr.) cDNA was sequenced. **d)** Region annotation of the Visium HD V1 sample. **e)** Number and **f)** percentage of significant spatially variable isoforms (FDR < 0.05). **g)** Overlap of significant SVIs between the V1 ONT and V1 PB data. Only isoforms tested in both datasets are considered. **h)** Overlap of significant SVIs between Sample1+2 (AE) ONT and V1 (PB) data. Only isoforms tested in both datasets are considered. **i)** Observed and expected overlap of SVIs (p = 3.5e-94, two-sided Fisher’s exact test). **j)** Relative expression of *Gad2*-new2 in V1 PB. Every hexagon shows the mean relative expression of the underlying cells. Relative expression indicates the fraction of long reads for that gene (*Gad2*) that are assigned to our isoform of interest (new2). A hexagon is only plotted when there are underlying cells expressing *Gad2* (Methods). The H&E staining is shown in the background. The schematic diagram in the corner shows the gene structure of *Gad2* isoforms. HPC = hippocampus, SVI = spatially variable isoforms, ALL = all cells, EX = excitatory neurons, IN = inhibitory neurons, OLI = oligodendrocytes, AST = astrocytes.

V2 and V3 were generated without long-molecule and exome selection, resulting in a shorter overall read length and a lower percentage of spliced reads compared to V1 and the Stereo-seq data (**Fig 5b-c, Tables S21-25**). As a consequence, fewer reads have an isoform assigned in V3-ONT compared to V1-ONT (7.2M vs. 8.4M), even though four ONT flow cells were used for V3-ONT compared to one for V1-ONT. For both ONT and PB, applying long-molecule selection and exome enrichment thus greatly reduces the sequencing cost per informative read (**Table S26-27**).

Similar to the Stereo-seq-based samples, the Visium HD slides are coronal sections of the adult mouse brain but cut slightly more anteriorly. As a result, all slides contain some similar brain regions, such as cortex and hippocampus, but other regions, for example, midbrain, striatum, and hypothalamus, are specific to certain slides (**Fig 5d, S29-30**). Using RCTD, we assigned cell types to the segmented cells in V1 and 8μm bins in V2 and V3 (**Fig S28-30**) (see Methods). In all Visium HD samples, some cells or 8μm bins (27.1% for V1, 55.5% for V2, and 29.3% for V3) could not be annotated using RCTD because of the low read depth, most likely due to the use of long-read data as input (**Fig S31-33**).

Next, we applied Spl-IsoFind to the Visium HD data to detect SVIs (**Fig 5e, Table S28-32**). Compared to the Stereo-seq-based data, relatively more SVIs were detected in astrocytes and fewer in excitatory neurons, which is most likely explained by the different cell type proportions in the Stereo-seq-derived and Visium HD-derived data (**Fig 4c, 5f**). We detected more SVIs in the PB data compared to ONT. For V1, 4,676 genes were tested in both the ONT and PB data (**Fig S34a**). Among these shared tested isoforms, all except one SVIs detected in the ONT dataset were significant in the PB data as well, whereas multiple SVIs were detected exclusively in the PB data (**Fig 5g**). We observed a similar pattern when comparing SVIs detected in V2-ONT and V3-PB, which represent very similar coronal sections. Again, the SVIs greatly overlapped and most platform-specific SVIs were detected only in the PB data (**Fig S34b**). This suggests a higher sensitivity in the PB datasets, which is most likely explained by the higher accuracy of barcode assignments and isoform assignments (**Table S33**). We also looked at whether the same pattern was present in the Stereo-seq data, although the PB Stereo-seq data had lower read depth. Here, we did not see a clear advantage for PB, which could be due to the lower PB sequencing depth or differences in barcode structure between platforms (**Fig S34c**).

In total, 4,976 isoforms were tested in both Sample1+2 AE and V1 (PB), of which 130 and 213 were significant, respectively (**Fig S35**). The significant overlap of 87 isoforms between the two experiments indicates strong reproducibility across platforms and biological replicates (p = 3.5e-94, two-sided Fisher’s exact test) (**Fig 5h-i**). For instance, *Arpp19* had four significant isoforms in Sample 1+2 (AE), three of which were also detected in all Visium HD samples (**Fig S36**). Different SVIs between the two Stereo-seq and Visium HD data can reflect regional specificity. For example, two *Gad2* isoforms were significant only in V1 (PB), V2 (ONT), and V3 (PB), likely due to their expression in the striatum, which is absent in the Stereo-seq samples (**Fig 5j, S37**). The two *Gad2* SVIs are both newly discovered isoforms using IsoQuant. *Gad2*-new1 is predicted to encode a 471 aa protein that lacks part of the N-terminal region compared to the canonical isoform *Gad2*-201, which is not spatially variable in our data. *Gad2*-new2 is predicted to encode a shorter 189 aa product. Highly similar protein sequences are annotated in human, with strong sequence conservation (97.9% for the 471 aa isoform and 88.2% for the 189 aa isoform). The 189 aa isoform lacks an in-frame stop codon and is annotated in human as a likely target of nonsense-mediated decay (NMD), suggesting that it may represent a regulatory rather than protein-coding transcript. Prior work in the human cortex has reported developmental and disease-associated shifts in full-length canonical versus NMD-associated GAD2 transcripts^65^.

## Discussion

Here, we present Spl-ISO-Seq2, and the accompanying software packages Spl-IsoQuant-2 and Spl-IsoFind, which allows studying spatially variable isoforms (SVIs) at single-cell and thus a cell-type-specific resolution for the first time. Spl-ISO-Seq2 has spot sizes far smaller than any known cells in the mouse brain. Thus, the vast majority of long reads can be attributed to a single cell, even achieving single-cell resolution for smaller cell types such as oligodendrocytes, which are thought to be 6-8μm in the mouse brain^66,67^. As such we can pinpoint isoforms that show cell-type-specific spatial variation, some of which have not been shown before through spatial transcriptomics. Some notable examples of cell-type-specific SVIs include *Rps24*, with regional isoform regulation specific to oligodendrocytes, and *Snap25* for excitatory neurons.

Furthermore, our developed software, Spl-IsoQuant-2 and Spl-IsoFind, is versatile and can be applied to data generated using all single-cell and spatial protocols. Spl-IsoQuant-2 has been tested on long reads derived from 10x 3’ single-cell, 10x Visium, Visium HD 3’, and Curio Biosciences’ Slide-seqV2. Moreover, Spl-IsoQuant-2 can take into account any user-specified molecule structure. Even though the Visium HD data was generated from a more anterior coronal cut, many spatially variable isoforms were detected in both datasets. Applying exome enrichment and long-molecule selection to Visium HD samples dramatically improves read length and number of spliced reads leading to broadly comparable results with respect to similarly treated Stereo-seq cDNA. For both protocols, Visium HD and Stereo-seq, exome enrichment and long-molecule selection allow for similar results while using one quarter of the flow cells compared to naively sequencing cDNA.

Previously, we and others showed that two mutually exclusive exons of *Snap25* exhibit spatially defined inclusion^4,19^. However, these initial spatial long-read approaches were based on the 10X Genomics’ Visium technology. Due to spot sizes of ∼60μm, which exceed the size of most cells in mouse and human brain, we could not conclusively determine the cell-type contribution to these spatially variable isoforms. Using Spl-ISO-Seq2, which has single-cell resolution, we revealed that *Snap25* region-specific isoform expression is indeed observed when only considering excitatory-neuron spots.

Moreover, we complement our new wet lab and barcoding methods with a new tool, Spl-IsoFind, to detect splicing regulation that does not conform with well-established anatomical regions. Using our previously developed *n* x 2 isoform tests^4^ we could easily test for genes exhibiting differentially used isoforms between predefined brain regions such as cortex vs. hippocampus, or between cortical layers. However, many other spatially-derived isoform patterns may not conform to known brain structures. Here, we implement a region-agnostic detection method, based on Moran’s I^33^ and contrast it to genes detected by comparing pre-defined brain regions. Using predefined regions increases the number of overall detected events in comparison to the region-agnostic approach. On the other hand, the region-agnostic approach detects events that were not observable with the predefined-regions approach. Indeed, in addition to the previously detected cortical pattern of *Snap25*, this gene also shows a switch in exon inclusion in a restricted area of the midbrain – a region that gave no prior indication of being of specific interest for splicing investigation. Thus, the region-agnostic approach has value in terms of finding isoform regulation specific to unknown areas.

In addition to region- and cell-type-specific differences, several genes, including *SNAP25*, exhibit sex-dependent splicing^56–58^, raising the possibility that SVIs identified in the male mouse brain may not fully translate to females. Using human single-nucleus long-read RNA sequencing data from three female and three male hippocampal samples, we identified 229 exons with sex-specific inclusion differences, predominantly in excitatory neurons. Although this analysis is limited by small sample size and substantial inter-individual variability, it suggests that spatial and sex-dependent splicing regulation can intersect. These findings highlight sex as an important biological variable and underscore the need to study this in more detail.

A current limitation of Spl-IsoFind is its low detection rate, which is partially explained by read depth. For example, we discovered more than three times as many SVIs in Sample 1, where we used two ONT flow cells, compared to Sample 2, which was sequenced using only one. Besides sequencing depth, tissue placement affected the number of informative reads: in Sample 2, many reads mapped to the opposite hemisphere and were excluded from downstream analysis. We therefore recommend only sequencing the region of interest. In general, our saturation analysis showed that cell-type-agnostic SVIs begin to plateau at approximately one PromethION flow cell per coronal section, whereas cell-type-specific SVIs do not show clear saturation. We therefore recommend using one to two PromethION flow cells or PB SMRT cells, depending on whether the primary goal is global SVI discovery or cell-type-resolved analyses. Finally, for PB sequencing, library configuration strongly influences the number of sequenced and informative reads. Samples 1 and 2 were prepared using 16 - fold MAS-Seq concatenation, whereas V1 used 12-fold concatenation (Kinnex). Because long-molecule selection produced relatively long molecules (**Fig 5b**), 16-fold concatenation likely generated molecules exceeding optimal sequencing length and reduced efficiency. Matching the fold concatenation to read length, as in V1, substantially improved throughput. Alternatively, the statistical power could be increased computationally by imputing isoform-level expression using matched single-cell data. Similar approaches have been developed for short-read spatial transcriptomics^68–70^, although applying such methods would require careful validation to avoid false positives. In addition, optimizing the neighborhood size or combining results across multiple neighborhood sizes to avoid relying on a single definition may further increase detection power.

The number of cell-type-specific SVIs could further be improved by decreasing the number of doublets in the data, thereby increasing the number of cells per cell type. However, due to 10µm slices, it is not always possible to completely exclude the presence of long reads from two distinct cells at the same cartesian (x,y) coordinate but varying in the “z” direction. Glial cells especially often form doublets^23^, which may explain our low number of singlets for oligodendrocytes and astrocytes and the correspondingly low number of detected SVIs. Current thresholds for detecting doublets might be overly strict, but we prioritize minimizing false positives, as is also reflected in our Spl-IsoQuant-2 parameters.

Furthermore, cell segmentation is currently based on nuclear staining only, expanding each segmented nucleus outward to a set radius or until it touches a neighboring cell. Because cells vary in size and morphology, this approach has clear limitations. Using additional stains to mark full cell boundaries could improve segmentation accuracy and thus reduce the number of doublets in future work.

Interesting extensions to Spl-ISO-Seq2 would include simultaneous measurements of proteins and/or open chromatin as has been done before in single-cells^10,71^. This could reveal regulatory relationships such as the interplay between promoters and alternative transcription-start-site usage as well as the interplay between RNA and protein isoforms, and how these relationships may be regulated spatially.

In summary, our wet lab and dry lab methods increase the resolution of spatial and single-cell isoform research. By combining high-resolution spatial data with isoform detection at the cell-type level, our approach represents a crucial step forward in understanding the spatially regulated mechanisms of splicing in complex tissues at a real single-cell resolution.

## Methods

### Ethics Statement

All experiments were conducted in accordance with relevant NIH guidelines and regulations, related to the Care and Use of Laboratory Animals tissue. Animal procedures were performed according to protocols approved by the Research Animal Resource Center at Weill Cornell College of Medicine.

### Animals and Tissue Processing

1 male C57BL P56 mouse was anesthetized with isoflurane and sacrificed via decapitation following IACUC protocols for Stereo-seq and subsequent sequencing. The brain was quickly removed, embedded in a cryocube with Tissue-Tek OCT (VWR 25608-930) and frozen at - 80C. The brain was cut at 10um thick using a Dakewe 6250 Cryostat Microtome and immediately mounted onto the Stereo-seq Chip Slide.

### Stereo-seq Protocol

After tissue sections (10um) were adhered to the Stereo-seq chip surface they were incubated at 37C for 5 minutes. Then, the sections were fixed in methanol and incubated for 30 minutes at -20C. Subsequently, the same sections were stained with nucleic acid dye (Thermo fisher, Q10212) and imaging was performed with a EVOS m7000 microscope prior to in situ capture at the channel of GFP. Tissue permeabilization, tissue removal, reverse transcription, and cDNA release was then performed based on Stereo-seq transcriptomics set for chip-on-a-slide user manual. cDNA was then amplified for 15 cycles and cleaned up using SPRI beads (Beckman Coulter, California, USA). cDNA quality was assessed by TapeStation High Sensitivity D5000 (Agilent Technologies Inc., California, USA) and quantified by Qubit™ dsDNA Quantification Kit (Thermofisher, Massachusetts, USA). Final library was constructed based on the manufacturer’s protocol. The quality was then assessed by TapeStation High Sensitivity D1000 (Agilent Technologies Inc., California, USA) and quantified by Qubit™ dsDNA Quantification Kit.

### DNB Sequencing

Short-read sequencing of final libraries was performed on the DNBSEQ-T7RS using the DNBSEQ-T7RS Stereo-seq Visualization Reagent Set (T7 STO FCL PE100) with a configuration of Read1: 50 (26-40 dark) cycles, Read2: 100 cycles, Barcode: 10 cycles.

### Linear PCR of cDNA

We first performed a linear/asymmetric PCR on the cDNA aliquoted for long-read experiments with the purpose of appending additional basepairs to the 3’ end of reads in order to avoid any TSO-TSO artifacts. The first asymmetric PCR used primer 5’-CTTCCGATCTATGGCGACCTTATCAG-3’ with the following settings: 95 °C for 3 min, 12 cycles of 98 °C for 20 s, 64°C for 30 s and 72 °C for 60 s. The resulting cDNA product was cleaned up with 0.8× SPRIselect beads (Beckman Coulter, B23318) and washed with 80% ethanol twice. This cDNA was then inputted into a symmetric PCR again using primer 5’-CTTCCGATCTATGGCGACCTTATCAG-3’ and TSO primer 5’-CTGCTGACGTACTGAGAGGC-3’ with 95 °C for 3 min, 8 cycles of 98 °C for 20 s, 64°C for 30 s and 72 °C for 60 s. Amplified cDNA was again purified with 0.8× SPRIselect beads (Beckman Coulter, B23318) and washed with 80% ethanol twice.

### Exome Enrichment of cDNA

500 ng of the cDNA collected from the previous step was used as input for the exome enrichment. Exome capture was performed as previously described to selectively amplify spliced molecules^5^. Agilent probe kit SureSelectXT Mouse All Exon (Agilent, 5190-4641) and the reagent kit SureSelectXT Mouse All Exon library (Agilent, G9611A) were used according to the manufacturer’s manual. First, the block oligonucleotide mix was made by combining 1 ul of LPCR primer (5’-CTTCCGATCTATGGCGACCTTATCAG-3’) and 1 ul of TSO primer (5’-CTGCTGACGTACTGAGAGGC-3’) at 200 ng/ul. We next combined 5 ul of 50-100 ng/ul of linearly amplified cDNA (see section above) and diluted it with 2 ul blockmix and 2 ul nuclease free water (NEB, AM9937). This mix was incubated on a thermocycler at 95 °C for 5 min, 65 °C for 5 min and 65 °C on hold. Next, 20 ul of SureSelect Hyb1, 0.8 ul of SureSelect Hyb2, 8.0 ul of SureSelect Hyb3 and 10.4 ul of SureSelect Hyb4 were combined at room temperature to make hybridization mix. Once the cDNA block mix reached to 65 °C on hold, 5 µl of probe mix, 1.5 µl of nuclease-free water, 0.5 µl of 1:4 diluted RNase Block and 13 µl of the hybridization mix were added to the cDNA block oligo mix and incubated for 24 h at 65 °C. M-270 Streptavidin Dynabeads (Thermo Fisher Scientific, 65305) were washed three times and resuspended with 200 µl of binding buffer after 24 hours of incubation. The incubated cDNA-block mix was mixed with the M270 Dynabeads and placed on a Hula mixer for 30 minutes. After this incubation time, the bead buffer was replaced with 200 µl of wash buffer 1 (WB1) followed by another 15-min incubation with low speed. Next, the WB1 was replaced with WB2, and the tube was transferred to the thermocycler for the next round of incubation (10 minutes). WB2 was replaced 2 more times at 10 minute intervals with the pre-heated WB2. Following this last incubation, beads were resuspended in 18 ul nuclease free H20 and then the spliced cDNA, which is attached to the beads, was amplified as described below.

### 3.3K Gene Set Enrichment of cDNA

We also enriched linear-PCR amplified cDNA with a custom gene set of 3,378 mouse ortholog genes related to various brain diseases and synaptic function, as we have published previously (**Table S34**)^10^. All experimental procedures remained the same as the exome enrichment and the “SureSelectXT Mouse All Exon” probes were replaced with the custom set (Agilent Part No. 5191-6900).

### Long-molecule Capture and Amplification

The long-molecule selection of spliced cDNA was performed as previously published^20^. Spliced cDNA bound with the M-270 Dynabeads was amplified with primers (LPCR: 5’-CTTCCGATCTATGGCGACCTTATCAG-3’; TSO: 5’-CTGCTGACGTACTGAGAGGC-3’) and KAPA-HiFi enzyme by using the following PCR protocol: 95°C for 3 min, 8 cycles of 98°C for 20 s 64°C for 60 s and 72°C for 3 min. The amplified cDNA was isolated from M-270 beads as supernatant and followed by a size selection/purification with 0.48× SPRIselect beads (in 1.25M NaCl-20% PEG buffer) and then eluted in 25ul EB buffer. All 25 ul size-selected spliced cDNA was used as template for the second round amplification of 6-8 cycles with the same PCR conditions suggested above. The product of the second round PCR was size selected/purified with 0.48× SPRIselect beads (in 1.25M NaCl-20% PEG buffer) and then eluted in 50 ul EB buffer.

### Oxford Nanopore Library Preparation

For the 2 samples, ∼75 fmol exome enriched and long-molecule selected cDNA was used as input into the ONT Ligation Sequencing Kit (ONT, SQK-LSK114), and Sample 1 (AE) was sequenced twice. Library construction was performed according to the manufacturer’s protocol (Nanopore Protocol, Amplicons by Ligation, version ACDE_9163_v114_revO_29Nov2022). The ONT library was loaded onto a PromethION sequencer by using PromethION Flow Cell (ONT, FLO-PR114M) and sequenced for 72 h. Base-calling was performed with high-accuracy settings on the Oxford Nanopore MinKNOWUI dorado basecaller.

### PacBio Library Preparation

Exome enriched and long-molecule selected cDNA was also sequenced with PacBio using the PacBio Mas-Seq for 10X 3’ Concatenation kit (102-659-600). As this kit was meant for cDNA coming from the 10X Chromium Next GEM Single Cell 3’ kit, we performed an initial PCR such that the correct sequences would be added to the ends of each molecule such that the 3’ and 5’ ends resembled those from 10X (10x_PR1_Stereo: 5’-CTACACGACGCTCTTCCGATCTCTTCCGATCTATGGCGACCTTATCAG-3’; 10x_PR2_Stereo: 5’-AAGCAGTGGTATCAACGCAGAGCTGCTGACGTACTGAGAGGC-3’).

The PacBio Mas-Seq for 10X 3’ Concatenation kit was used following PacBio’s published protocol. The resulting library was sequenced using the PacBio Revio using 1 SMRT cell per sample. Reads were split by identifying known read-linking sequences using skera (version 1.4.0).

### Tissue isolation and reverse transcription for exon validation

To validate spatially variable isoforms found by comparing two regions, we isolated the midbrain region from the remaining portion of the brain from three P56 mice. RNA was isolated from 30-50mg of these regions using the Qiagen miRNA Tissue/Cells Advanced Mini Kit (Qiagen Cat. no. 217684) following the established protocol. RNA was frozen at -80C. Next, we performed reverse transcription using the SuperScript III First-strand Synthesis System (Invitrogen, 18080051) for RT-PCR kit following the manufacturer’s protocol. We combined 5uL RNA with 0.5uL random hexamers, 0.5uL oligodT, 1uL dNPTs, and 3uL H20. This mix was incubated at 65C for 5 minutes followed by being placed on ice for 1 minute. We next added 2 uL 10x Buffer, 4 uL MgCl2, 2uL DTT, 1uL RNAse Out, and 1 uL SuperScriptIII to the samples and incubated at 25C for 10 minutes, 50C for 50 minutes, and 85C for 5 minutes. The cDNA was then stored at -20 until further use.

### PCR exon validation

cDNA was amplified using target primers (see below) using the following settings: 95 °C for 3 min, 26 cycles of 98 °C for 20 s, 64°C for 30 s and 72 °C for 60 s. Resulting cDNA was inputted onto an Agilent Tapestation System 4200 using D1000 ScreenTapes following the manufacturer’s protocol.

### Primers used for QPCR Exon Validation

Nptn

5’-CTCTTGCTGGTCTCTGGCTC-3’ and 5’-GTGGCAGTGAGTTCTACCCC-3’

Phactr1

5’-CTGTCGGAAATGGAGCCTGT-3’ and 5’-CGGAGGCGACAGTTTCACT-3’

Gapdh

ReadyMade_GAPDH_Fwd (IDT catalog #51-01-07-12): 5’-ACCACAGTCCATGCCATCAC-3’

ReadyMade_GAPDH_Rvs (IDT catalog #51-01-07-13): 5’-TCCACCACCCTGTTGCTGTA-3’

### Tapestation Gel Quantifications

Tapestation gel images were saved and quantified in FIJI using established gel quantification methods^72^. Values per sample were normalized to that sample’s Gapdh levels. Exon inclusion values were calculated using 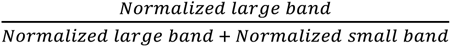 per sample and brain region. For *Nptn*, normalized exon inclusion = 539 base pair (bp) band/(191 bp+539 bp band). For *Phactr1*, normalized exon inclusion = 345 base pair (bp) band/(345 bp+138 bp band).

### Long-read processing

For upstream analysis of PB and ONT reads we implemented Spl-IsoQuant-2 — a new version of the Spl-IsoQuant pipeline (based on IsoQuant^24^). Spl-IsoQuant-2 is capable of processing long-read data obtained with the Stereo-seq and 10x Visium HD spatial protocols, as well as 10x Visium, Curio Biosciences Slide-seqV2, 10x 3’ v2/v3 single-cell protocols. Additionally, Spl-IsoQuant-2 implements a universal barcode calling algorithm that allows a user to process long-read data obtained with any custom protocol (not limited to the ones described above) by providing a molecule description file (see below).

Spl-IsoQuant-2 starts with identifying barcode and UMI sequences in every read sequence. Then, the reads are mapped to the reference genome with minimap2^73^ in spliced mode and assigned to known isoforms and genes using existing IsoQuant algorithms. Finally, Spl-IsoQuant removes PCR duplicates and performs gene and transcript quantification. Below we provide details on every step of the pipeline.

To account for unannotated isoforms, in this work we first ran standard bulk IsoQuant pipeline using Gencode version 36 mouse basic reference annotation and combination of all Stereo-seq long-read data (3 ONT and 2 PB samples, 391 million reads in total). Default settings specific for ONT data were chosen to avoid potential false positives. The generated GTF, which contained 28’891 known and 26’445 novel isoforms, was further provided into the Spl-IsoQuant-2 pipeline as the reference annotation. To enable a fair comparison with the third-party data from Lebrigand et al. we similarly performed novel isoform discovery by treating the data as bulk and subsequently running spatial analysis using an updated gene annotation.

### Stereo-seq barcode detection

Due to similarity of primers at 3’ and 5’ ends Stereo-seq protocol multiple barcoded cDNAs may be concatenated. To account for this fact, Spl-IsoQuant-2 implements an algorithm that is capable of detecting multiple barcoded cDNAs within a single read. The algorithm first identifies the poly(T) sequence, the 3’ primer (STOmics sequencing primer in Stereo-seq notation) and the linker (fixed sequence between the barcode and the UMI sequence, see protocol). Further, it detects the closest TSO primer following the detected poly(T) stretch (**Fig 2b**) and repeats this procedure iteratively for the remaining part of the read. Iterative primer detection is then repeated for the reverse-complemented read sequences, thus allowing for reverse concatenation. The resulting set of non-overlapping sequences located between 3’ primer and 5’ TSO are outputted as individual cDNA sequences.

These individual subsequences are then used for detecting barcode and UMI sequences (CIDs and MIDs in Stereo-seq notation respectively) (**Fig 2c**), as well as the remaining pipeline steps, such as read alignment and feature quantification. The barcode sequence extracted from a cDNA (located between the 3’ primer and the linker, see **Fig 2c**) is matched against the set of barcodes from the input whitelist. Only cDNA with barcodes with the alignment score above a certain threshold S_min_ are reported (see details below); cDNAs with non-matching barcodes are rejected. UMI sequence is reported as a read subsequence located between the linker and the start of the poly(T) stretch.

Here we exploit the same pattern matching as in the previous version of Spl-IsoQuant^20^, which is based on short k-mer indexing and conventional Smith-Waterman local alignment (via the ssw-py library^74^). To efficiently detect sequences from a given set (e.g. barcodes) or a fixed sequence (e.g. linker or primer), we use k-mer indexing: a hash table where each key is a k-mer and the value is the set of known sequences containing it. Given a read, we extract all of its k-mers and use the index to retrieve potential matches. To “parse” the read according to the molecule structure, we exploit three separate indices: for the TSO primer (23 bp, k=8), the 3’ primer (26 bp, k=6) and the linker (15 bp, k=5). These indices are used to quickly detect potential positions of a fixed sequence in a read, which are then verified via Smith-Waterman local alignment.

Once the positions of the primer and the linker are identified, the barcode sequence is extracted (between the primer and the linker) and potential matches are identified among the known barcodes using a k-mer index constructed for the input barcode whitelist (25 bp, k=14). While smaller k-mer length increases sensitivity for detecting the correct known barcode even in the presence of sequencing errors, it also raises the chance of spurious matches, especially in large barcode sets (>400 million for Stereo-seq protocol). Therefore, k-mer matching serves only to shortlist potential matches. Top N (N=10 by default) potential matches with the highest number of shared k-mers are validated using Smith-Waterman local alignment. Extracted barcode sequence is required to have an alignment score at least S_min_ (22 by default, classic alignment scoring system: +1 for match, -1 for indel/mismatch) with the closest candidate from the whitelist. In this case the barcode from the whitelist is reported as the correct barocode. If two or more known barcodes have the same highest alignment score or all potential matches have scores below S_min_, no barcode is reported for this cDNA. Thus, the k-mer index reduces the search space and limits the number of costly alignment operations needed for barcode identification, while the alignment itself helps to filter out possible false positive matches and uncertain candidates.

Finally, to enable processing of Stereo-seq data typically featuring extremely large barcode whitelists (>450 million) we substantially reworked the algorithm implementation (see performance statistics below).

### Benchmarking Stereo-seq barcode detection

To evaluate the barcode calling algorithm, we simulated ONT data using a modified version of Trans-NanoSim^75^, available at https://github.com/andrewprzh/lrgasp-simulation. This version improves read truncation procedures, enabling realistic ONT truncation probabilities and non-truncated reads, and was previously used for IsoQuant benchmarking^24^. The error model was trained with actual sequencing data from this project (ONT sequencing, Sample 2), showing a sequencing error rate of 3.8% (1.5% mismatches, 1.1% insertions, 1.2% deletions), ensuring accurate recall and precision estimates. No read truncation was induced during the simulation process.

In this work we assessed the barcode calling method using two simulated datasets: (1) the one where each read contains a single cDNA sequence, and (2) another one where each read can contain from 1 to 4 concatenated cDNAs. Molecule templates were created using Gencode reference transcript sequences according to the Stereo-seq molecule structure (**Fig 2c**). Reference transcripts were randomly selected based on the expression profile derived from the Sample 2. For the second dataset, we randomly concatenated between 1 and 4 cDNA templates. The probability of each cDNA benign reverse-complemented prior to concatenation is 0.4. While these parameters might not exactly match the ones for real data, the resulting simulated dataset contained a substantial fraction of concatenated reads (likely exceeding average concatenation levels observed in real data, see Supplementary Table S3 for details), thus allowing to accurately assess the algorithm capabilities. In both experiments we simulated 10 million reads.

We ran Spl-IsoQuant-2 on the simulated datasets and compared the detected barcodes against the true barcodes attached during simulation. Correctly identified barcodes were considered true positives, incorrectly reported barcodes as false positives, and undetected barcodes as false negatives. For reads containing multiple cDNAs, each cDNA was counted individually (**Fig 2e**).

We also tried running Flexiplex on our data as it allows to specify custom molecule structure with user-defined primers. Unfortunately, after running for 5 days in 16 threads on a dataset containing just 1 million ONT reads it was able to detect less than 20 thousand barcodes (<2% of reads with detected barcodes). Similarly, it was not able to provide any reasonable result for simulated data (2 detected barcodes for the entire dataset). In contrast, as mentioned above, Spl-IsoQuant-2 allows indexing ∼450 million barcodes in less than 10 minutes, and processes 60/10 million long reads per hour without/with deconcatenation respectively in 16 threads. Spl-IsoQuant-2 bacode detection rate is ∼55% on real data and ∼75% on simulated data.

### Barcode detection for other spatial and single-cell protocols

Spl-IsoQuant-2 also implements barcode detection algorithms for other spatial (10x Visium, Visium HD 3’, Curio Biosciences’ Slide-seqV2) and single-cell (10x 3’ v2 and v3) protocols by adapting the concept described above. To facilitate future biotechnological innovations we also implemented a universal barcode calling algorithm suitable for any custom protocol. While Flexiplex allows specifying custom adapter sequences, Spl-IsoQuant allows a user to describe any custom molecule structure, including sophisticated cases with multiple barcodes (such as 10x Visium HD), barcodes separated by a linker sequence in the middle (such as Curio Biosciences’s Slide-seqV2 spatial protocol), or the same barcode being attached to the cDNA sequence more than once. This functionality enables upstream processing of long-read data sequenced via any custom single-cell or spatial sequencing methods without developing novel software.

The main complication for barcode calling in 10x Visium HD data is the similarity between the barcode sequences of neighbouring spots and variable barcode lengths (14-16bp), not typical for any other protocol. To keep high precision we decided to keep a rather high alignment score cutoff (13). Using simulated 10x Visium HD data with the same error rate as in Stereo-seq experiments (3.8%) we show that Spl-IsoQuant-2 outperforms existing SpaceRanger pipeline adapted for long-reads by 10x Genomics (**Fig 5a**, **Table S33**). For 2μm and 8μm resolutions our algorithm generates 2.4x and 3.5x fewer false positives compared to SpaceRanger with just slightly lower recall. On PB simulated data it achieves nearly perfect results with 99.76% precision and 99.03% recall.

### Read processing and UMI deduplication

Overall, the Spl-IsoQuant-2 pipeline is similar to its previous version. Individual cDNA sequences extracted from raw reads are mapped to the reference genome using minimap2 with IsoQuant default settings (-x splice -k 14 --junc-bed annotation.bed -a --MD -Y --secondary=yes for ONT and -x splice -k 14 --junc-bed annotation.bed -a --MD -Y --secondary=yes for PB). Spl-IsoQuant then assigns these mapped reads to reference genes and isoforms using the IsoQuant standard algorithm^20^. Multimapped reads are resolved by selecting the most consistent alignment with the reference annotation or discarding them. Reads uniquely assigned to a known gene undergo further processing to remove PCR duplicates.

To eliminate PCR duplicates, reads are grouped by gene and barcode. Within each group, reads with identical or highly similar UMI sequences (edit distance ≤ 4) are considered duplicates. We use an iterative algorithm to identify and assign reads to representative UMIs, selecting the read covering the most splice junctions as the representative and ignoring others. These UMI-filtered reads are then used for downstream analyses, such as exon and transcript quantification and differential splicing analysis. While this method may occasionally remove non-duplicate reads due to similar UMIs, the randomness of UMI sequences minimizes this impact on the analysis.

### Comparison PacBio and ONT isoform assignment

For reads with a barcode-UMI-gene combinations sequenced with both PB and ONT, we compared the assigned isoforms by Spl-ISO-Seq2 only when both platforms assigned an isoform.

### Cell segmentation and barcode to cell assignment

Cells were segmented based on the ssDNA staining and short-read gene expression using the SAW^76^ pipeline (version 7). For nucleus segmentation, SAW version 7 relies on a built-in deep-learning model trained on nucleus-stained images to segment nuclei from the ssDNA staining. After segmentation, cell boundaries are estimated by expanding each nucleus by up to 10 barcodes (corresponding to 5 μm) or until the expanded boundaries of neighboring cells collide. This generates several outputs including a masking image indicating the cell boundaries, a barcode-to-position mapping file, and a GEF (gene expression file) that contains the CellID for each x and y coordinate combination. However, this GEF only includes coordinates that have at least one short-read assigned. If we used this file directly to map coordinates (and thus barcodes) to the cells, we would lose the coordinates that only have long-reads but no short-reads assigned. To ensure that all coordinates inside a segmented cell were assigned a CellID, we generated an artificial GEM (gene expression matrix) with one short-read for each x and y coordinate. Using the `saw convert bin2cell` function and the previously obtained masking file, we converted this artificial GEM into a new cell-binned GEF. This new GEF includes the CellID for all coordinates inside a cell.

### (Sub)region annotation

Regions and Subregions were annotated manually by using a combination of gene expression markers, clustering analyses, cell type distribution, and known brain region architecture. Regions based on these criteria were outlined using the Polygon function, turned into masks, and uploaded such that all spots within the masked region were annotated as so.

### Cell-type identification

We used Robust Cell Type Decomposition (RCTD) (V2.2.1)^37^ to assign cell types to segmented cells and identify singlets and doublets. For the Spl-ISO-Seq2 datasets, we input gene expression matrices obtained from the short-read sequencing data. For the Visium HD datasets, short-read data were unavailable, so we constructed gene expression matrices using the long reads. We used a reference^9^ file including six P56 mouse single-cell samples from multiple brain regions including hippocampus, visual cortex, striatum, and thalamus. Previously annotated “doublets” from the single-cell data were removed prior to analysis. We ran RCTD in “doublet” mode allowing each segmented cell to be modeled as consisting of either one or two cell types. We ran RCTD with default parameters following published tutorials (https://github.com/dmcable/spacexr/blob/master/vignettes/spatial-transcriptomics.Rmd). Spots designated as “singlets” were assigned as cell types per sample.

### Slide alignment

We aligned Sample 1 and Sample 2 using the `spateo.align.morpho_align()` function from the spateo^28^ package (version 1.1.0) with parameters beta=0.1 and lambdaVF=1.0. This greatly improved the alignment compared to naively overlapping the slides (**Fig 1d, S38**)

### Spatially variable isoform tests using predefined regions

We used the isoform assignments from Spl-IsoQuant-2 to detect spatially variable isoforms instead of using full-length reads only. For each region of interest, we counted the number of unique reads per isoform. Afterwards, we used the `DiffSplicingAnalysis()` function from the ScisorSeqR^4^ package (version 0.1.9) to identify genes that exhibit differential isoform usage between the predefined brain regions using the default settings (minNumReads = 25, numIsoforms = 10). Using these default settings, genes with less than 25 reads are filtered out. NumIsoforms means that we tested a maximum of 10 isoforms per gene and grouped the remaining isoforms in a rest group, which results in an 11x2 count matrix at most. The returned p-values are calculated using a chi-squared test and corrected for multiple testing using Benjamini Hochberg correction. Genes with FDR <= 0.05 and delta Pi >= 0.1 are considered significant. Delta Pi was calculated for each gene as the sum of change in percent isoform from the top two isoforms detected. We tested all pairwise combinations for the general brain regions, hippocampal subregions, and cortical layers using all cells and per cell type.

### Differential exon testing using predefined comparisons

We called differentially expressed exons as we have previously^10,20^ done. After executing Spl-IsoQuant on the long-read sequencing data, which mapped, barcoded, and spliced reads while correcting for UMIs, we generated an AllInfo file. This file contains detailed metadata for each read, including: read identifier, gene name, comparison group (such as specific cell type or brain region), barcode, UMI, intron chain, transcription start site (TSS), polyadenylation site, exon chain, transcript classification (known or novel), and intron count. You can find further information on AllInfo files at: https://github.com/noush-joglekar/scisorseqr ^4^.

AllInfo files were split based on the specific comparisons being analyzed. These were analyzed using the casesVcontrols() function in the scisorATAC R package (https://github.com/careenfoord/scisorATAC)^10^, with a threshold of 10 minimum reads per exon, to identify differentially included exons using standard parameters.

The process to detect alternative exons involved the following steps:

1. Read Selection We used reads from both comparison groups (e.g., cortex barcodes vs. midbrain barcodes) that contain introns and adhere to canonical splice site rules.
2. Alternative Exon Identification For each candidate exon, we tallied: A: Reads supporting both splice junctions of the exon B: Reads supporting the left splice site, ending within the exon on the right C: Reads supporting the right splice site, starting within the exon on the left D: Reads skipping the exon (but assigned to the same gene) E: Reads overlapping the exon in any fashion
3. Filtering Criteria

○ Exclude exons where A + B + C + D < 10, due to insufficient read coverage
○ Exclude exons where (A + B + C + D)/(A + B + C + D + E) < 0.8, indicating insufficient clarity between inclusion and exclusion
○ Calculate PSI (Percent Spliced In):

▪ leftPSI = (A + B) / (A + B + D)
▪ rightPSI = (A + C) / (A + C + D)
▪ Keep exons where both PSI values are within [0.05, 0.95]
4. Cluster-Specific PSI Calculation For exons that passed all prior filters:

○ Calculate group-specific PSI = (A + B + C) / (A + B + C + D), only if A + B + C + D ≥ 10
○ Construct a 2x2 contingency table using exon inclusion and skipping counts for each group (e.g., layer 4 old vs. young)
○ Retain tables where at least 3 of the 4 cells have expected values ≥ 5
5. Statistical Testing

Each retained exon is tested for association between group identity and exon inclusion using a two-sided Fisher’s exact test. Resulting p-values are adjusted for multiple comparisons via the Benjamini-Yekutieli correction.

### Comparison to Visium coronal brain slides

We downloaded FASTQ files from the coronal brain slides published by Lebrigand et al.^19^ from the Sequence Read Archive under accession numbers SRX8673336 and SRX8673337. We processed these reads using Spl-IsoQuant-2 and used ScisorSeqR as described earlier to detect genes that exhibit differential isoform usage. Since these datasets were annotated at a finer anatomical level, we mapped their original region annotation to the region annotations we used (**Table S15**). Finally, we detected spatially variable isoforms using Spl-IsoFind as described below.

### Sex-specific exon-inclusion analysis

We investigated sex-specific splicing differences using a single-nucleus long-read dataset consisting of hippocampal samples from three males and three females^9^. We downloaded FASTQ files here: https://knowledge.brain-map.org/data/ASP3B09DZ8PXDUYSHDH. We processed these using Spl-IsoQuant-2 and tested for sex-specific exon inclusion differences by aggregating reads per sex and performing exon-level comparisons within each cell type (see differential exon testing using predefined comparisons). We used the cell-type labels defined in the original publication. Exons were considered significant if they passed a BY-adjusted p-value threshold of 0.05 and showed an absolute ΔPSI > 0.1 between males and females. To reduce the influence of outlier individuals, we additionally required that at least two out of three individuals per sex had sufficient read coverage (≥10 reads) and that all individuals with sufficient coverage agreed on the direction of the effect.

### Spatially variable isoform tests using Moran’s I

We used the `esda.Moran` function from the ESDA^77^ python package (version 2.4.3) to calculate Moran’s I and the corresponding p-value using a one-tailed permutation test based on 1e5 permutations. The input data for this analysis consists of the relative expression of the tested isoform in every cell. For the spatial weights, we focused only on cells that express the corresponding gene in the long-read data. The weight for the *k* (default=10) nearest neighbors of each cell was set to 1, while all other neighbors were assigned a weight of 0.

To reduce the number of tested isoforms, we only included isoforms that met the following criteria:

- The corresponding gene is expressed in at least 5\**k* (default=50) cells in the long-read data.
- At least 20 cells express the tested isoform and at least 20 cells express another isoform (to ensure some minimum expression of the isoform of interest and variance in the data).
- The fraction of cells expressing the tested isoform should be between 0.05 and 0.95 (to ensure some variance in the data).

The p-values are corrected for multiple testing using the Benjamini-Yekutieli method. Isoforms are considered significant if FDR <= 0.05 and a Moran’s I score > 0.01.

When comparing to Geary’s C, we replaced the `esda.Moran` function by `esda.Geary`. Both options are available in the Spl-IsoFind package.

### Using simulations to assess Moran’s I precision and recall

For all simulations, we used the original coordinates of the cells from Sample 1. To assess precision, we generated random data by assigning values of 0 and 1 randomly to all cells. We simulated different imbalances in the distribution of 0s and 1s (50-50, 40-60, 30-70, 20-80, 10-90, 5-95). To evaluate precision, we simulated relative expression values based on four patterns:

1. *Frontiers*: A clear border is simulated between cells with a value of 1 and cells with a value of 0.
2. *Square*: A square region within the tissue is filled with 1s, with surrounding cells assigned 0. Different square sizes (200, 500, 1000, 2000 µm) were tested.
3. *Square - surrounded by random points*: Similar to the square pattern, but the square is surrounded by cells with random values (either 0 or 1).
4. *Line - surrounded by random points*: Instead of a square, a horizontal or vertical line of 1s is simulated, surrounded by random points on both sides. Different line widths (100, 250, 500 µm) were tested.

Each scenario was simulated 100 times and downsampled to different cell numbers (50, 100, 500, 1000, 5000, 10000 cells). After downsampling, Moran’s I and the corresponding p-value were calculated as described earlier. If downsampling resulted in violating any of the predefined criteria for Moran’s I (e.g., fewer than 20 cells per category), we did not calculate the Moran’s I score or p-value for those cases.

### GO-enrichment analysis

We used the enrichGO() function in clusterProfiler v4.18.4^78^ with the BP ontology and an FDR cutoff of q < 0.05. We used the simplify() function to remove redundancy among the terms.

### Saturation analysis

For S1 (AE), S2 (AE), and S1 (3.3K), we randomly subsampled the sequenced reads to 10%, 20%,…, 90% of the total read count. We repeated each subsampling step ten times to account for stochastic variation.

### Cell-type-constrained permutations

By default, we permute all cells randomly, which means that the relative expression values of each cell are shuffled across all cells, regardless of their assigned cell type. To investigate which SVIs are caused by differences in cell-type abundances, we performed cell-type-constrained permutations. In this approach, only cells with assigned cell-type labels could be used. We used cells that RCTD labeled as ‘singlet’ or ‘doublet_certain’. For these doublets, we assigned a label based on the RCTD assigned doublet probabilities For singlets, we used the assigned cell-type labels, and for doublets, we included only those for which both cell types were clearly identified. During each permutation, the doublets were assigned a cell-type label with probabilities based on their assigned doublet probabilities. Next, we randomly shuffle the values of the cells within each specific cell type. Due to the increased computational time, we performed 1e4 permutations (1e4) instead of the usual 1e5.

In Figure 4e, we compared the p-values from this cell-type-constrained permutation method to those obtained using normal random permutations. Since the original p-values were calculated using all cells and 1e5 permutations, we recalculated the normal p-values as well using 1e4 permutations and only cells with an assigned cell type.

### Spatial hexagon plots

To avoid overlapping cells or very small cells, we decided to plot spatial isoform expression using spatial hexagon plots. Every hexagon shows the mean relative expression of the underlying cells. Relative expression indicates the fraction of long reads for a gene that are assigned to our isoform of interest. As such, a hexagon is only plotted when there are underlying cells expressing the corresponding gene. A relative expression of 1 thus indicates that all underlying long reads express the isoform of interest, while a relative expression of 0 indicates that all underlying long reads express another isoform.

### Visium HD 3’ long-read sequencing

For V1, we received cDNA from 10x Genomics. 10x Genomics generated the cDNA as follows. A mouse brain (Male, 8 weeks, RIN=8) was obtained from Charles River Laboratories. A 10 µm section was taken with a cryostat (Epredia CryoStar NX70). Tissue preparation, sectioning, H&E staining, and imaging followed the Visium HD 3’ Fresh Frozen Tissue Preparation Handbook (CG000804). Generation of cDNA followed the Visium HD 3’ Spatial Gene Expression User Guide (CG000805).

We did exome enrichment and long molecule selection as previously published^20^. Spliced cDNA bound with the M-270 Dynabeads was amplified with primers (LPCR: 5′-CTACACGACGCTCTTCCGATCT-3′; TSO: 5′-AAGCAGTGGTATCAACGCAGAG-3′ and KAPA-HiFi enzyme by using the following PCR protocol: 95°C for 3 min, 8 cycles of 98°C for 20 s 64°C for 60 s and 72°C for 3 min. The amplified cDNA was isolated from M-270 beads as supernatant and followed by a size selection/purification with 0.48× SPRIselect beads (in 1.25M NaCl-20% PEG buffer) and then eluted in 25ul EB buffer. All 25 ul size-selected spliced cDNA was used as template for the second round amplification of 6-8 cycles with the same PCR conditions suggested above. The product of the second round PCR was size selected/purified with 0.48× SPRIselect beads (in 1.25M NaCl-20% PEG buffer) and then eluted in 50 ul EB buffer.

For ONT sequencing, ∼100 fmol exome enriched and long-molecule selected cDNA was used as input into the ONT Ligation Sequencing Kit (ONT, SQK-LSK114). Library construction was performed according to the manufacturer’s protocol (Nanopore Protocol, Amplicons by Ligation, version DCS_9187_v114_revK_10Dec2025). The ONT library was loaded onto PromethION Flow Cell (ONT, FLO-PR114M) and sequenced on PromethION sequencer for 72 h. Base-calling was performed with high-accuracy settings on the Oxford Nanopore MinKNOWUI Dorado basecaller.

For PB sequencing, 35 ng exome enriched and long-molecule selected cDNA was used as input into the Kinnex 16S rRNA kit (PacBio, cat #103-072-000). Library construction was performed according to the PacBio’s protocol from Kinnex PCR to ABC step. The PacBio library was sequenced in 1 SMRTcell. The fastq was demultiplexed with Skera. For V2, we downloaded publicly available Visium HD 3’ ONT long-read data from a P56 male mouse brain from https://epi2me.nanoporetech.com/visium_hd_2025.06/.

For V3, we downloaded publicly available Visium HD 3’ PacBio long-read data from a P56 male mouse brain from: https://downloads.pacbcloud.com/public/dataset/Kinnex-single-cell-RNA/DATA-RevioSPRQ-Kinnex-VisiumHD-mouseBrain/1-Sreads/.

### Visium HD 3’ data processing

The algorithm for barcode detection for Visium HD 3’ ONT data was implemented based on the Visium HD 3’ molecule structure in a similar manner as the described Stereo-seq barcode calling method, but without read deconcatenation (see above). K-mer size of 7 was used for detecting potential positions of R1 primer (22 bp) and both barcode parts (14-16 bp each). For alignment-based barcode validation, the default minimal score S_min_=13 was chosen for each of the barcode parts.

The implemented barcode finding algorithm was benchmarked and compared against Space Ranger on Visium HD 3’ mouse simulated data. The data was generated using Trans-NanoSim^75^ in a similar way as Stereo-seq simulated data (3.8% error rate, see above), but without cDNA concatenation (i.e. one read contains a single cDNA molecule). The reported barcodes were converted to Visium HD slide spot ids and assessed at three resolution levels: 2 μm, 8 μm and 16 μm (**Fig 5a**).

For V1, we used Space Ranger (v 4.0.1) for cell segmentation. A high-resolution microscope image is not available for V2 and V3. As an alternative for cell segmentation, we used the 8μm bins and used those as a proxy for cells. For all Visium HD datasets, we ran RCTD on long-read gene expression data. Afterwards, we applied Spl-IsoFind as described above.

### Code availability

Spl-IsoQuant-2 is an open source software and is openly available at https://github.com/algbio/spl-IsoQuant.

Spl-IsoFind is available at https://github.com/tilgnerlab/Spl-IsoFind

## Data availability

Processed data is available for interactive visualization at spatialisoforms.weill.cornell.edu.

### Funding statement and acknowledgements

We thank Wodan Ling for carefully evaluating the statistical validity of Spl-IsoFind. We thank Adrian Tan, Chendong Pan, Aihong Liu, Seongeun Oh and Jenny Xiang from the Genomics Resources Core Facility at Weill Cornell Medicine for performing RNA sequencing. We also thank Weill Cornell Medicine Scientific Computing Unit (SCU) for use of their computational resources.

Figure 1a, 4a, and S19 were created using BioRender.com. Supported by: MIRA R35 GM152101-01 (H.U.T), Brain Initiative grant 1RF1MH121267-01 (H.U.T.), NIDA U01 DA053625-01 (H.U.T among others), NIDA grant 2T32DA039080 (J.H.), NSF GRFP # 2139291 (C.F.), the Feil Family Foundation (H.U.T.), European Union (European Research Council, SCALEBIO, 101169716. Views and opinions expressed are however those of the author(s) only and do not necessarily reflect those of the European Union or the European Research Council. Neither the European Union nor the granting authority can be held responsible for them) (A.D.P, A.I.T.), Research Council of Finland grants No. 322595, 346968, 358744 (A.D.P, A.I.T., R.P.), NIGMS Maximizing Investigators’ Research Award (MIRA) R35 GM138152 (I.H.).

### Competing interests

H.U.T. has presented at user meetings of 10× Genomics, Oxford Nanopore Technologies, and Pacific Biosciences, which in some cases included payment for travel and accommodations.

H.U.T. has recently agreed to consult for ISOgenix Ltd., for work unrelated to the present manuscript. The other authors declare no competing interests.

## Notes

### Competing Interest Statement

H.U.T. has presented at user meetings of 10x Genomics, Oxford Nanopore Technologies, and Pacific Biosciences, which in some cases included payment for travel and accommodations. H.U.T. has recently agreed to consult for ISOgenix Ltd., for work unrelated to the present manuscript. The other authors declare no competing interests.

### Summary of Updates

Updated results. Added extra Visium HD data.

