## Supplement for "Spatial isoform sequencing at sub-micrometer single-cell resolution reveals novel patterns of spatial isoform variability in brain cell types"

**Table S1.** Stereo-seq short-read statistics

| | Sample 1 | Sample 2 |
|---|---|---|
| Number of segmented cells | 91,690 | 108,845 |
| Mean number of barcodes per cell | 753 | 642 |
| Median number of barcodes per cell | 702 | 586 |
| Mean number of UMIs per cell | 1,353 | 925 |
| Median number of UMIs per cell | 1,144 | 792 |
| Mean number of genes per cell | 678 | 499 |
| Median number of genes per cell | 614 | 452 |

**Table S2.** Stereo-seq ONT barcode calling statistics

| | Sample 1 (AE) | Sample 1 (3.3K) | Sample 2 (AE) |
|---|---|---|---|
| Total reads | 207,282,773 | 87,520,126 | 58,799,050 |
| cDNAs extracted | 218,287,258 | 88,514,020 | 62,775,129 |
| PolyT detected | 182,228,343 | 56,831,671 | 54,111,424 |
| Primer detected | 159,898,493 | 39,694,971 | 48,318,281 |
| Linker detected | 160,091,513 | 39,749,743 | 48,380,796 |
| Barcode detected | 130,360,866 | 31,182,006 | 40,354,173 |
| PolyT detected | 83.48% | 64.21% | 86.20% |
| Primer detected | 73.25% | 44.85% | 76.97% |
| Linker detected | 73.34% | 44.91% | 77.07% |
| Barcode detected | 59.72% | 35.23% | 64.28% |

**Table S3.** Stereo-seq ONT barcode calling statistics: the effect of de-concatenation barcode calling algorithm

| Dataset | S1 (AE) | S2 (AE) |
|---|---|---|
| Total reads | 207,282,775 | 58,799,051 |
| cDNAs extracted | 218,287,260 | 62,775,130 |
| Barcoded cDNAs | 129,178,416 | 40,354,173 |
| Barcoded cDNAs without deconcatenation | 118,173,931 | 36,378,094 |
| Reads with >= 2 cDNAs | 10,458,285 | 3,903,874 |
| Barcoded cDNAs in these reads | 21,462,770 | 7,879,953 |
| Additional barcoded cDNAs recovered | 11,004,485 | 3,976,079 |
| **Additional barcoded cDNAs recovered (%)** | **9.31%** | **10.93%** |

**Table S4.** Stereo-seq PacBio barcode calling statistics

| | Sample 1 (AE) | Sample 2 (AE) |
|---|---|---|
| Total reads | 19,878,737 | 17,627,912 |
| cDNAs extracted | 20,137,382 | 17,969,619 |
| PolyT detected | 17,365,347 | 15,487,476 |
| Primer detected | 16,951,711 | 15,368,317 |
| Linker detected | 16,952,851 | 15,369,386 |
| Barcode detected | 15,195,233 | 13,981,555 |
| PolyT detected | 86.23% | 86.19% |
| Primer detected | 84.18% | 85.52% |
| Linker detected | 84.19% | 85.53% |
| Barcode detected | 75.46% | 77.81% |

**Table S5.** Stereo-seq ONT read-to-isoform assignment and PCR deduplication statistics

| | Sample 1 (AE) | Sample 1 (3.3K) | Sample 2 (AE) |
|---|---|---|---|
| Total reads | 207,282,773 | 87,520,126 | 58,799,050 |
| cDNAs extracted | 218,287,258 | 88,514,020 | 62,775,129 |
| Barcode detected | 130,360,866 | 31,182,006 | 40,354,173 |
| **Of them** | | | |
| Uniquely assigned | 100,571,599 | 27,204,522 | 30,483,380 |
| Uniquely assigned and spliced | 79,030,635 | 25,712,877 | 24,159,223 |
| Unique gene-barcode pairs | 35,128,134 | 7,500,561 | 14,910,661 |
| Reads after deduplication | 36,728,127 | 7,923,634 | 15,295,364 |
| Spliced reads after deduplication | 25,916,277 | 7,016,938 | 11,134,293 |
| Barcode detected | 59.72% | 35.23% | 64.28% |
| **Of them** | | | |
| Uniquely assigned | 77.15% | 87.24% | 75.54% |
| Uniquely assigned and spliced | 60.62% | 82.46% | 59.87% |
| Reads after deduplication | 26.95% | 24.05% | 36.95% |
| Spliced reads after deduplication | 28.17% | 25.41% | 37.90% |

**Table S6.** Stereo-seq PacBio read-to-isoform assignment and PCR deduplication statistics

| | Sample 1 (AE) | Sample 2 (AE) |
|---|---|---|
| Total reads | 19,878,737 | 17,627,912 |
| cDNAs extracted | 20,137,382 | 17,969,619 |
| Barcode detected | 15,195,233 | 13,981,555 |
| **Of them** | | |
| Uniquely assigned | 12,427,208 | 11,407,337 |
| Uniquely assigned and spliced | 9,852,363 | 9,344,667 |
| Unique gene-barcode pairs | 8,479,274 | 7,453,247 |
| Reads after deduplication | 8,534,498 | 7,489,921 |
| Spliced reads after deduplication | 6,500,576 | 5,845,919 |
| Barcode detected | 75.46% | 77.81% |
| **Of them** | | |
| Uniquely assigned | 81.78% | 81.59% |
| Uniquely assigned and spliced | 64.84% | 66.84% |
| Reads after deduplication | 55.80% | 53.31% |
| Spliced reads after deduplication | 56.17% | 53.57% |

**Table S7.** Stereo-seq ONT statistics after PCR deduplication

| | Sample 1 (AE) | Sample 1 (3.3K) | Sample 2 (AE) |
|---|---|---|---|
| Spliced reads after deduplication | 25,916,277 | 7,016,938 | 11,134,293 |
| Overlapping with segmented cell in correct hemisphere | 10,741,771 | 3,054,136 | 3,104,550 |
| Isoform assigned | 8,006,816 | 2,062,768 | 2,406,135 |
| Unique genes detected | 22,434 | 12,198 | 18,003 |
| Mean number of reads per gene | 479 | 250 | 172 |
| Unique isoforms detected | 49,860 | 22,060 | 36,273 |
| Mean number of reads per isoform | 161 | 94 | 66 |
| Mean number of spliced reads per cell | 177 | 50 | 54 |
| Median number of spliced reads per cell | 140 | 39 | 44 |
| Mean percentage of spliced reads assigned to a gene per cell | 100% | 100% | 100% |
| Mean number of genes detected per cell | 152 | 43 | 50 |
| Median number of genes detected per cell | 125 | 35 | 42 |
| Mean percentage of spliced reads assigned to an isoform per cell | 75.5% | 67.2% | 77.4% |
| Mean number of isoforms detected per cell | 116 | 30 | 39 |
| Median number of isoforms detected per cell | 95 | 24 | 33 |

**Table S8.** Stereo-seq Pacbio statistics after PCR deduplication

| | Sample 1 (AE) | Sample 2 (AE) |
|---|---|---|
| Spliced reads after deduplication | 6,500,576 | 5,845,919 |
| Overlapping with segmented cell in correct hemisphere | 2,855,135 | 1,668,435 |
| Isoform assigned | 2,283,296 | 1,339,239 |
| Unique genes detected | 17,312 | 15,980 |
| Mean number of reads per gene | 165 | 104 |
| Unique isoforms detected | 34,879 | 30,142 |
| Mean number of reads per isoform | 65 | 44 |
| Mean number of spliced reads per cell | 47 | 29 |
| Median number of spliced reads per cell | 37 | 23 |
| Mean percentage of spliced reads assigned to a gene per cell | 100% | 100% |
| Mean number of genes detected per cell | 45 | 28 |
| Median number of genes detected per cell | 36 | 23 |
| Mean percentage of spliced reads assigned to an isoform per cell | 79.8% | 80.2% |
| Mean number of isoforms detected per cell | 36 | 22 |
| Median number of isoforms detected per cell | 28 | 19 |

**Table S9.** A comparison of read statistics between samples generated using the Stereo-seq, Visium HD, and Visium platforms. Informative reads are defined as UMI-deduplicated, spliced reads with an assigned isoform and overlapping a segmented cell or spot in the correct hemisphere. However, for the Lebrigand et al paper, a subset, containing 44M and 99M reads, respectively, was uploaded to SRA. The uploaded data likely represent reads already enriched for informative molecules, as the number of reads after UMI deduplication is consistent with those reported in their Supplementary Materials.

| | Stereo-seq Sample 1 (AE - ONT) | Visium HD V1 (ONT) | Visium HD V1 (PacBio) | Visium CBS1 | Visium CBS2 |
|---:|---:|---:|---:|---:|---:|
| Sequenced reads | 207,282,773 | 98,713,075 | 126,023,947 | 173,996,000 | 286,458,673 |
| Analysed reads | 207,282,773 | 98,713,075 | 126,023,947 | 43,899,878 | 98,892,086 |
| Barcode detected | 130,360,866 | 49,089,229 | 84,196,089 | 36,625,448 | 80,219,056 |
| Uniquely assigned | 100,571,599 | 38,561,039 | 66,968,834 | 31,032,886 | 67,505,940 |
| Uniquely assigned and spliced | 79,030,635 | 25,987,614 | 53,617,739 | 19,972,628 | 48,768,593 |
| Reads after deduplication | 36,728,127 | 22,313,578 | 18,616,621 | 17,581,867 | 18,489,755 |
| Spliced reads after deduplication | 25,916,277 | 14,740,254 | 13,620,397 | 10,929,678 | 12,189,522 |
| Informative reads | 8,006,816 | 8,354,635 | 6,982,558 | 8,204,667 | 8,749,685 |

**Table S10.** Differential relative isoform expression results for all cells for all tested genes. See Supplementary File.

**Table S11.** Differential relative isoform expression results for excitatory neurons for all tested genes. See Supplementary File.

**Table S12.** Differential relative isoform expression results for inhibitory neurons for all tested genes. See Supplementary File.

**Table S13.** Differential relative isoform expression results for oligodendrocytes for all tested genes. See Supplementary File.

**Table S14.** Differential relative isoform expression results for astrocytes for all tested genes. See Supplementary File.

**Table S15.** Mapping between the original and new labels for the Lebrigand et al. dataset.

| **Original label** | **New label** |
|---|---|
| CA1/CA2, CA2, DG, Hippocampus area | Hippocampus |
| Iscortex-1, Isocortex-2, Olfactory area, Retrosplenial area, | Cortex |
| Fiber tracts | White Matter |
| Thalamus | Thalamus |
| Midbrain, Hypothalamus | Midbrain |

**Table S16.** Spl-IsoFind results for all cells for all tested genes. See Supplementary File.

**Table S17.** Spl-IsoFind results for excitatory neurons for all tested genes. See Supplementary File.

**Table S18.** Spl-IsoFind results for inhibitory neurons for all tested genes. See Supplementary File.

**Table S19.** Spl-IsoFind results for oligodendrocytes for all tested genes. See Supplementary File.

**Table S20.** Spl-IsoFind results for astrocytes for all tested genes. See Supplementary File.

**Table S21.** Visium HD cell segmentation statistics

| | V1 |
|---:|---:|
| Number of segmented cells | 77,821 |
| Mean number of barcodes per cell | 46 |
| Median number of barcodes per cell | 42 |

**Table S22.** Read length statistics for Stereo-seq and Visium HD

| Sample | Total reads | Mean length, bp | N50, bp | N90, bp |
|---|---|---|---|---|
| **Stereo-seq ONT** | | | | |
| Sample 1 (AE) | 207,282,773 | 1,282 | 1,409 | 851 |
| Sample 1 (3.3K) | 87,520,126 | 1,416 | 1,486 | 1,032 |
| Sample 2 (AE) | 58,799,050 | 1,462 | 1,659 | 971 |
| **Stereo-seq PacBio** | | | | |
| Sample 1 (AE) | 19,878,737 | 1,099 | 1,270 | 737 |
| Sample 2 (AE) | 17,627,912 | 1,223 | 1,511 | 814 |
| **Visium HD V1 ONT** | | | | |
| | 98,713,075 | 1,089 | 1,193 | 702 |
| **Visium HD V1 PacBio** | | | | |
| | 126,023,947 | 1,049 | 1,096 | 762 |
| **Visium HD V2 ONT** | | | | |
| PBC44593 | 161,221,491 | 622 | 767 | 341 |
| PBC44760 | 132,795,836 | 626 | 769 | 342 |
| PBC49262 | 140,386,479 | 624 | 768 | 340 |
| PBC52788 | 165,541,042 | 624 | 771 | 338 |
| **Visium HD V3 PacBio** | | | | |
| | 108,063,589 | 694 | 743 | 443 |
| | 107,529,335 | 694 | 743 | 443 |
| | 102,934,038 | 693 | 743 | 443 |
| | 101,261,931 | 694 | 744 | 444 |

**Table S23.** Visium HD barcode calling statistics using Spl-IsoQuant-2

| | V1 (ONT) | V1 (PB) | V2 (ONT) | V3 (PB) |
|---|---|---|---|---|
| Total reads | 98,713,075 | 126,023,947 | 599,944,848 | 545,812,840 |
| Barcode detected | 49,089,229 | 84,196,089 | 268,552,874 | 388,488,422 |
| PolyT detected | 90,224,485 | 124,208,263 | 584,711,646 | 542,561,819 |
| R1 detected | 77,716,763 | 120,032,663 | 520,667,619 | 521,687,965 |
| Barcode detected | 49.73% | 66.81% | 44.76% | 71.18% |
| PolyT detected | 91.40% | 98.56% | 97.46% | 99.40% |
| R1 detected | 78.73% | 95.25% | 86.79% | 95.58% |

**Table S24.** Visium HD read-to-isoform assignment and PCR deduplication statistics.

| | V1 (ONT, 2μm) | V1 (PB, 2μm) | V2 (ONT, 8μm) | V3 (PB, 8μm) |
|---|---|---|---|---|
| Total reads | 98,713,075 | 126,023,947 | 599,944,848 | 419,788,893 |
| Barcode detected | 49,089,229 | 84,196,089 | 268,552,874 | 304,292,333 |
| **Of them** | | | | |
| Uniquely assigned | 38,561,039 | 66,968,834 | 180,099,678 | 286,182,637 |
| Uniquely assigned and spliced | 25,987,614 | 53,617,739 | 49,918,226 | 86,331,762 |
| Unique gene-barcode pairs | 21,728,795 | 18,504,392 | 44,088,597 | 72,985,533 |
| Reads after deduplication | 22,313,578 | 18,616,621 | 47,600,985 | 80,401,514 |
| Spliced reads after deduplication | 14,740,254 | 13,620,397 | 10,844,620 | 19,965,368 |
| Barcode detected | 49.73% | 66.81% | 44.76% | 72.49% |
| **Of them** | | | | |
| Uniquely assigned | 78.55% | 79.54% | 67.06% | 94.05% |
| Uniquely assigned and spliced | 52.94% | 63.68% | 18.59% | 28.37% |
| Reads after deduplication | 45.46% | 22.11% | 17.72% | 26.42% |
| Spliced reads after deduplication | 30.03% | 16.18% | 4.04% | 6.56% |

**Table S25.** Visium HD statistics after PCR deduplication

| | V1 (ONT) | V1 (PB) | V2 (ONT) | V3 (PB) |
|---|---|---|---|---|
| Spliced reads after deduplication | 14,740,254 | 13,620,397 | 10,844,620 | 19,965,368 |
| Isoform assigned | 8,354,635 | 6,982,558 | 7,186,605 | 12,604,203 |
| Unique genes detected | 21,211 | 20,263 | 27,283 | 29,319 |
| Mean number of reads per gene | 533 | 521 | 392 | 666 |
| Unique isoforms detected | 42,357 | 38,178 | 46,382 | 50754 |
| Mean number of reads per isoform | 197 | 183 | 155 | 248 |
| Mean number of spliced reads per cell/8um bin | 145 | 136 | 21 | 50 |
| Median number of spliced reads per cell/8um bin | 111 | 103 | 17 | 35 |
| Mean percentage of spliced reads assigned to a gene per cell/8um bin | 100% | 100% | 100% | 100% |
| Mean number of genes detected per cell/8um bin | 127 | 121 | 20 | 46 |
| Median number of genes detected per cell/8um bin | 102 | 96 | 17 | 33 |
| Mean percentage of spliced reads assigned to an isoform per cell/8um bin | 73.9% | 66.0% | 67.3% | 64.6% |
| Mean number of isoforms detected per cell/8um bin | 96 | 80 | 13 | 30 |
| Median number of isoforms detected per cell/8um bin | 76 | 64 | 11 | 21 |

**Table S26.** Estimated costs for the individual steps during each experiment. Per type of experiment, we also indicated which steps were included. The total cost per experiment is shown in Table S27. Shared steps (e.g., the Stereo-seq slide) are included for multiple experiments, even when the same slide was used in practice to get a fair estimate of the cost per read (e.g., as in Sample 1 (AE) and Sample 1 (3.3K)). The reported costs are an estimate and may vary across institutes depending on available infrastructure and vendor discounts.

| | Est. cost | S1/S2 (AE) ONT | S1/S2 (AE) PB | S1 (3.3K) ONT | V1 ONT | V1 PB | V2 ONT | V3 PB |
|---|---|---|---|---|---|---|---|---|
| Stereo-seq slide (including SR seq.) | $5000 | 1x | 1x | 1x | | | | |
| Visium HD slide (excluding SR seq.) | $4500 | | | | 1x | 1x | 1x | 1x |
| Long-mol. selection | $35 | 1x | 1x | 1x | 1x | 1x | | |
| Exome enrichment | $155 | 1x | 1x | 1x | 1x | 1x | | |
| 3378 gene enrichment | $350 | | | 1x | | | | |
| ONT library prep | $120 | 1x | | 1x | 1x | | 1x | |
| PromethION flow Cell | $1040 | 1-2x | | 1x | 1x | | 4x | |
| PacBio library prep | $475 | | 1x | | | 1x | | 1x |
| PacBio SMRT Cell | $1700 | | 1x | | | 1x | | 4x |

**Table S27.** Estimated total cost per experiment and cost per sequencing output. Costs are reported per 1M total reads and per 100K informative reads. Informative reads are spliced reads with an assigned isoform overlapping a segmented cell in the correct hemisphere and thus suitable for downstream analysis. Estimates are calculated for the complete workflow and for the long-read sequencing part alone.

| | Including array prep | | | Long-read sequencing only | | | |
|---|---|---|---|---|---|---|---|
| | Est. total cost | Cost per 1M seq. reads | Cost per 100K informative reads | Est. total cost | Cost per 1M seq. reads | Cost per 100K informative reads | |
| S1 (AE) - ONT | $7,390 | $36 | $92 | $2,390 | $12 | $30 | |
| S1 (AE) - PB | $7,365 | $368 | $323 | $2,365 | $118 | $104 | S1 (AE) - PB was sequenced using PacBio Mas-Seq with 16-fold concatenation resulting in fewer reads compared to V1 - PB, which was sequencing with the Kinnex kit with 12-fold concatenation |
| S1 (3.3K) - ONT | $6,700 | $76 | $325 | $1,700 | $19 | $82 | |
| S2 (AE) - ONT | $6,350 | $108 | $264 | $1,350 | $23 | $56 | For S2 (AE) - ONT, many reads were mapped to cells in the other hemisphere that we discarded. This largely accounts for the difference compared to S1 (AE) - ONT |
| S2 (AE) - PB | $7,365 | $409 | $550 | $2,365 | $131 | $177 | S2 (AE) - PB was sequenced using PacBio Mas-Seq with 16-fold concatenation resulting in fewer reads compared to V1 - PB, which was sequencing with the Kinnex kit with 12-fold concatenation |
| V1 - ONT | $5,850 | $59 | $70 | $1,350 | $14 | $16 | |
| V1 - PB | $6,865 | $54 | $98 | $2,365 | $19 | $34 | |
| V2 - ONT | $8,780 | $15 | $122 | $4,280 | $7 | $60 | In comparison to V1-ONT, this did not use exome enrichment and long-molecule selection. As a result, it needed more flow cells, raising the cost |
| V3 - PB | $11,775 | $28 | $93 | $7,275 | $17 | $58 | In comparison to V1-PB, this did not use exome enrichment and long-molecule selection. As a result, it needed more flow cells, raising the cost |

**Table S28.** Spl-IsoFind results for all cells for all tested genes in the Visium HD data. See Supplementary File.

**Table S29.** Spl-IsoFind results for excitatory neurons for all tested genes in the Visium HD data. See Supplementary File.

**Table S30.** Spl-IsoFind results for inhibitory neurons for all tested genes in the Visium HD data. See Supplementary File.

**Table S31.** Spl-IsoFind results for oligodendrocytes for all tested genes in the Visium HD data. See Supplementary File.

**Table S32.** Spl-IsoFind results for astrocytes for all tested genes in the Visium HD data. See Supplementary File.

**Table S33.** Statistics for Visium HD 3' simulated ONT and PacBio data

| | Data | ONT | | PacBio | |
|---|---|---|---|---|---|
| | Error rate | 3.8% | | 0.1% | |
| | Method | SpaceRanger | Spl-IsoQuant-2 | SpaceRanger | Spl-IsoQuant-2 |
| 2um | Precision | 94.18% | 97.58% | 99.61% | 99.76% |
| | Recall | 78.88% | 76.08% | 99.999% | 99.26% |
| 8um | Precision | 97.50% | 99.29% | 99.88% | 99.93% |
| | Recall | 79.45% | 76.39% | 99.999% | 99.26% |

**Table S34.** List of 3,378 targeted genes.

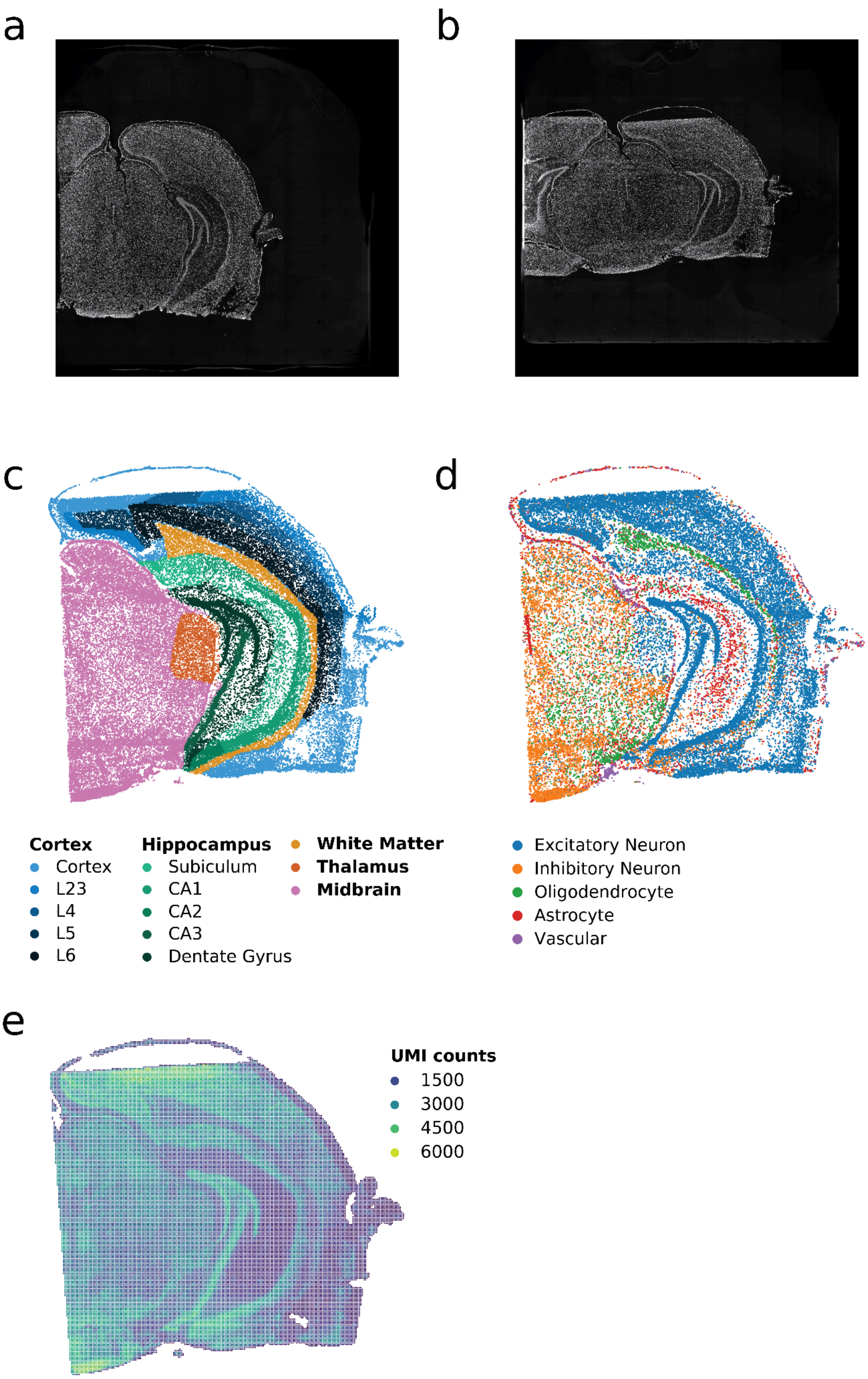

**Figure S1. a-b)** ssDNA staining of **a)** Sample 1 and **b)** Sample 2. **c)** (Sub)region annotation of Sample 2. **d)** Cell-type annotations of Sample 2. Only cells annotated as singlets and cell types with >100 cells are plotted. **e)** UMI counts per bin (50x50spots) for Sample 2.

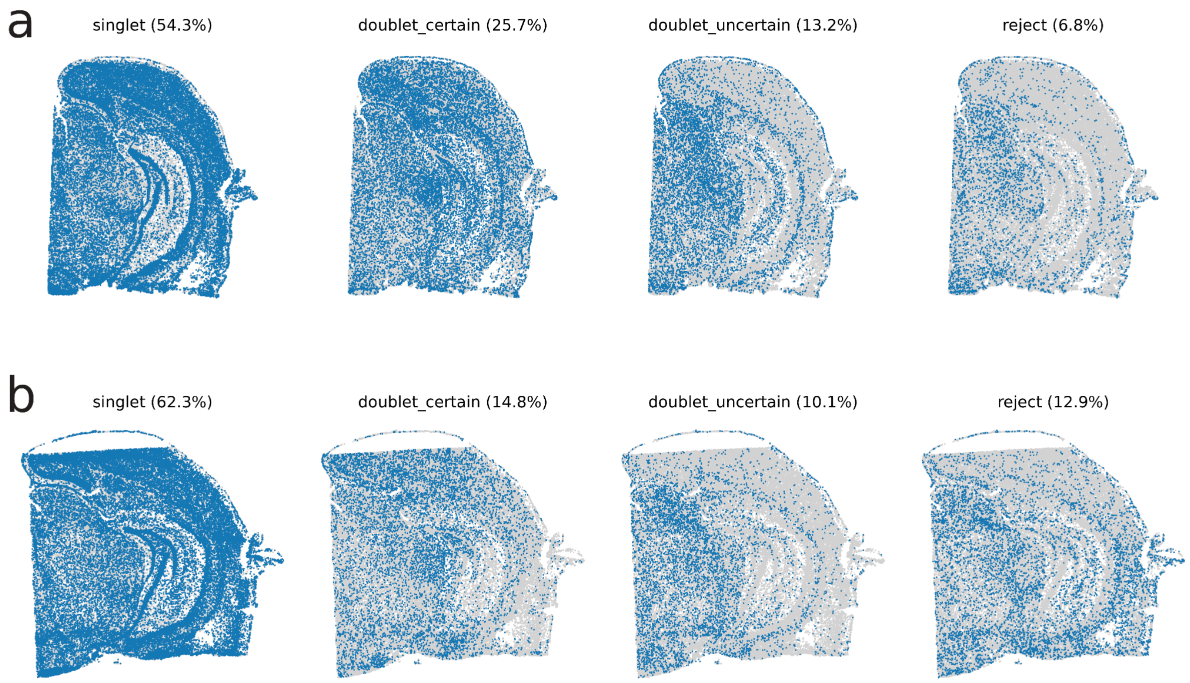

**Figure S2** RCTD cell-assignment distribution in **a)** Sample 1 and **b)** Sample 2.

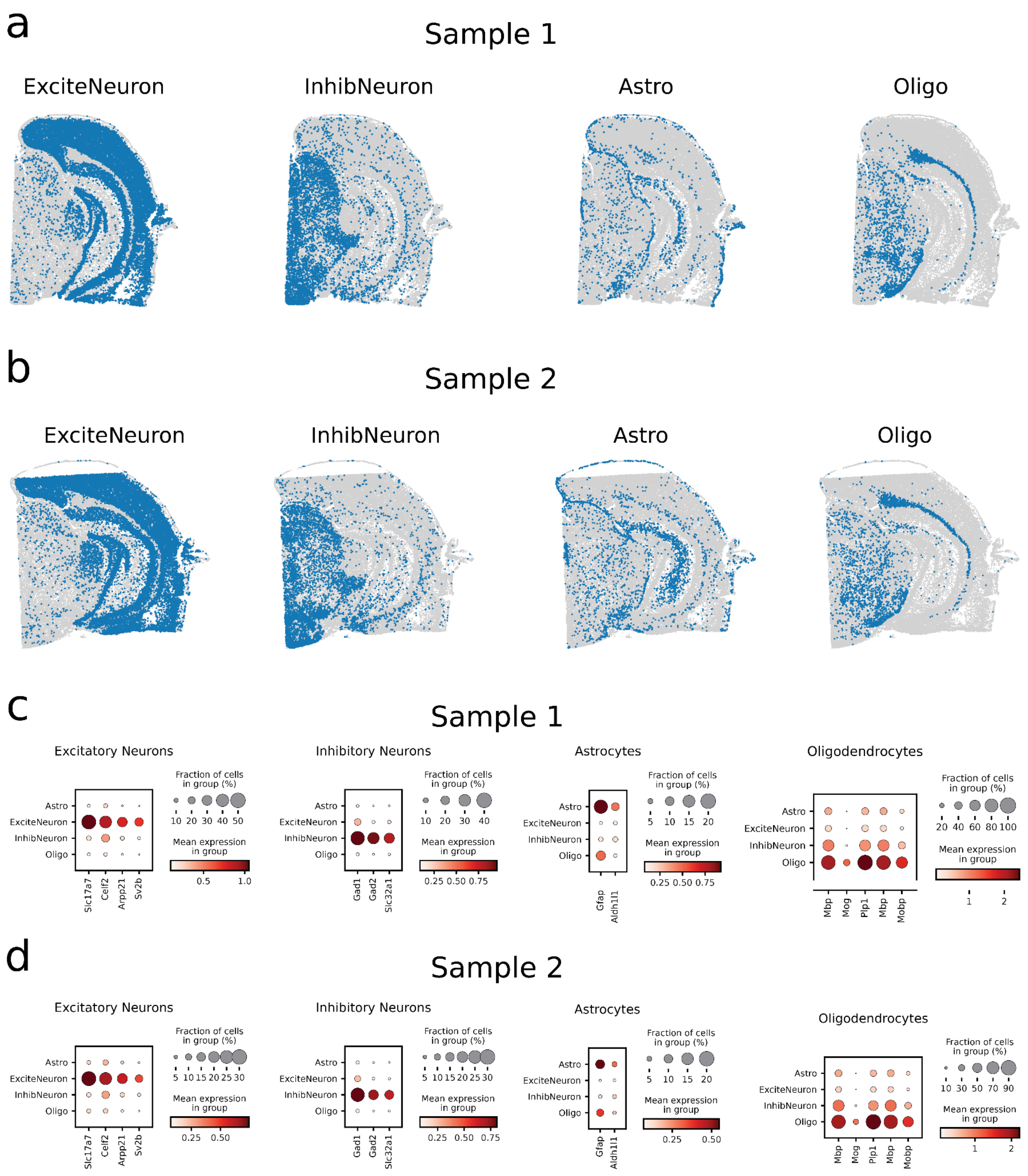

**Figure S3. a-b)** Cell-type distribution and **c-d)** marker gene expression of the four major cell types in **a,c)** Sample 1 and **b,d)** Sample 2. ExciteNeuron = Excitatory Neurons, InhibNeurons = Inhibitory Neurons, Astro = Astrocytes, Oligo = Oligodendrocytes

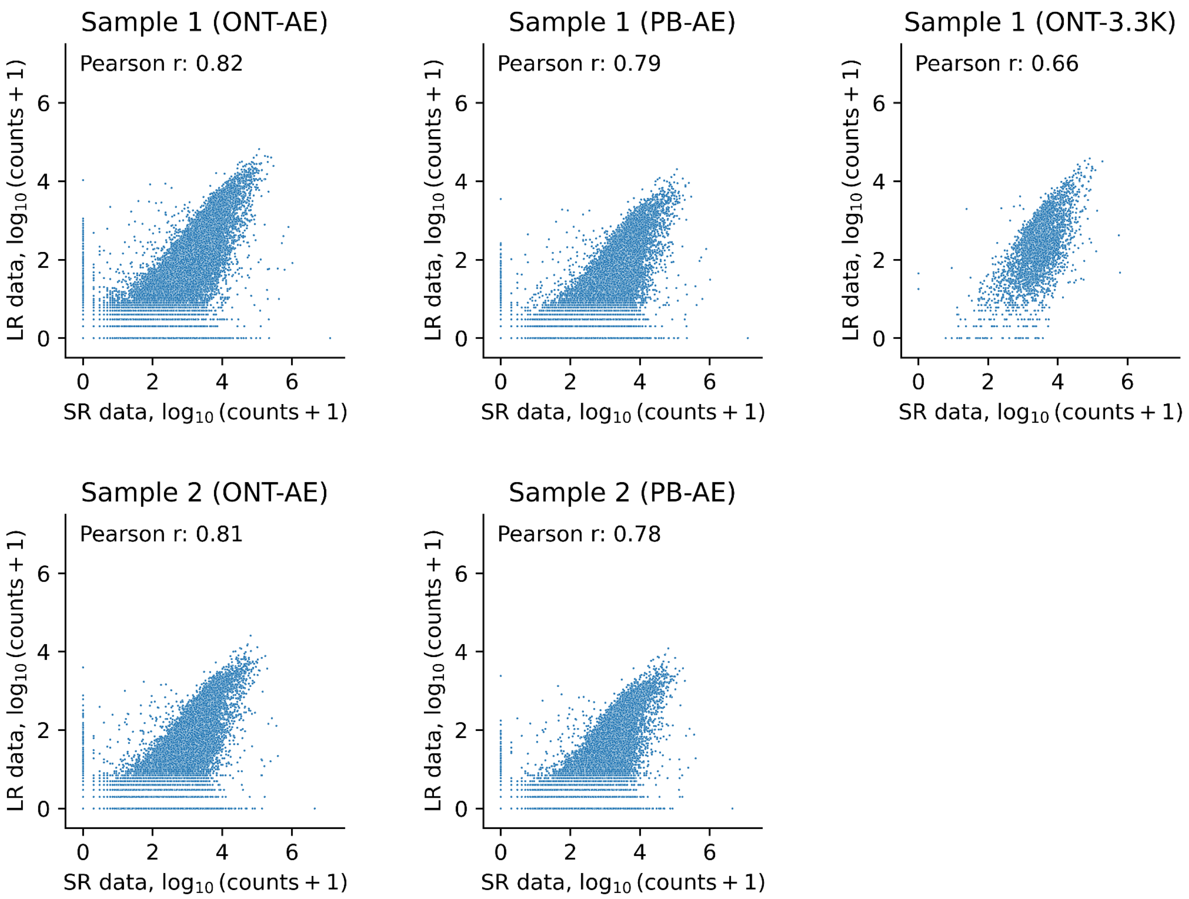

**Figure S4.** Correlation between gene expression as measured using the long-read and short-read sequencing. Counts are log10 transformed. Pearson correlation is indicated in the top left corner. Only the targeted genes are shown for Sample1 (3.3K). LR = long-read, SR = short-read, AE = allexome, PB = pacbio, 3.3K = 3.3K targeted dataset.

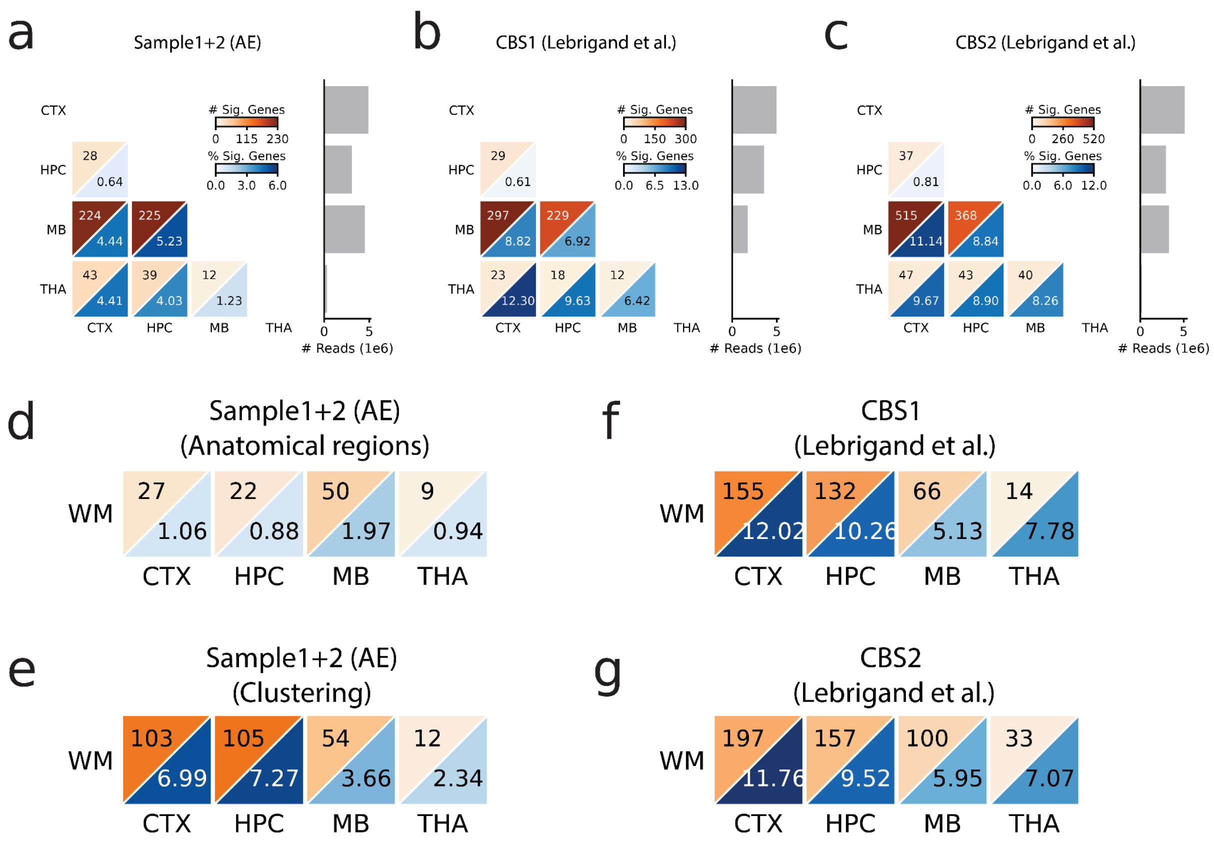

**Figure S5.** Number and percentage of genes changing isoform expression across predefined brain regions using all cells in **a)** combined Sample 1+2 (AE), **b)** Coronal Brain Section 1 (CBS1) from Lebrigand et al., and **c)** Coronal Brain Section 2 (CBS2) from Lebrigand et al. The barplot on the right shows the number of long reads used during the analysis per brain region. **d-g)** Number and percentage of genes changing isoform expression across predefined brain regions when comparing white matter to other brain regions in **d-e)** combined Sample 1+2 (AE) using regions defined based on **d)** anatomical locations and **e)** clustering the cells, **f)** Coronal Brain Section 1 (CBS1) from Lebrigand et al., and **g)** Coronal Brain Section 2 (CBS2) from Lebrigand et al..

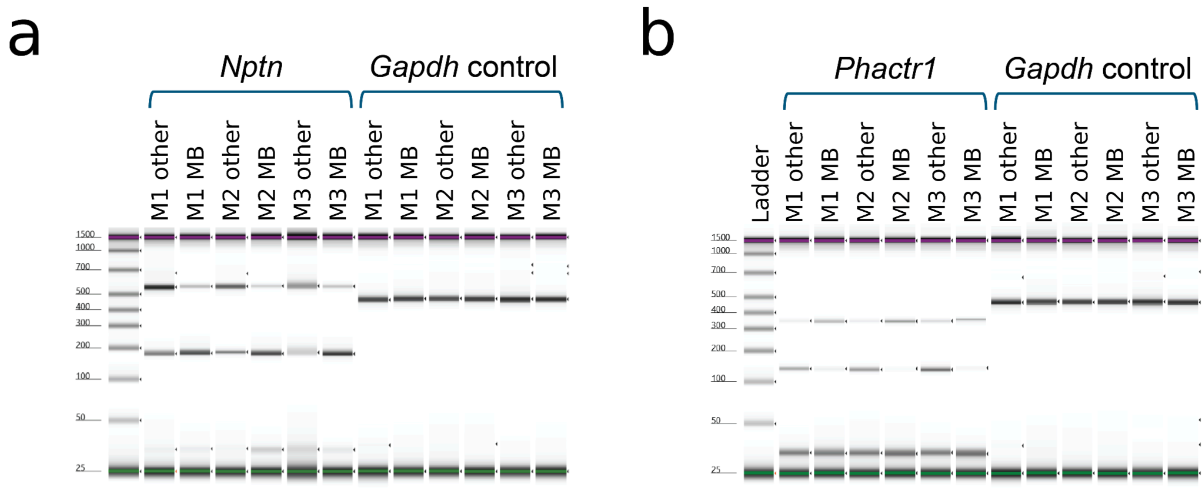

**Figure S6. a-b)** Tapestation gel of PCR amplified products using exon-of-interest-spanning **a)** *Nptn* and **b)** *Phatcr1* specific primers or *Gapdh* specific primers. Mx = Mouse sample, MB = isolated bulk midbrain, other = bulk brain with midbrain removed.

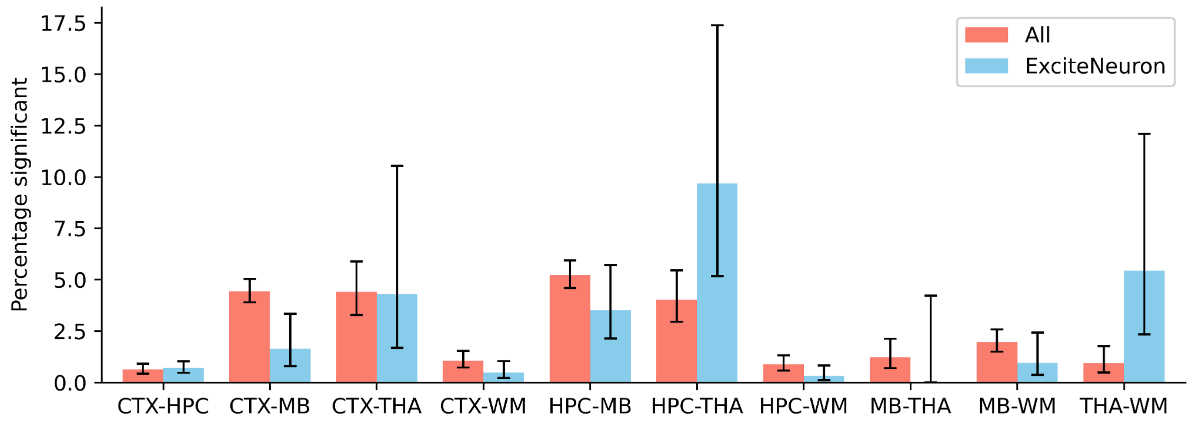

**Figure S7.** Percentage of significant genes for each comparison using all cells and only excitatory neurons. Error bars indicate 95% confidence intervals. CTX = cortex, HPC = hippocampus, MB = midbrain, THA = thalamus, WM = white matter,

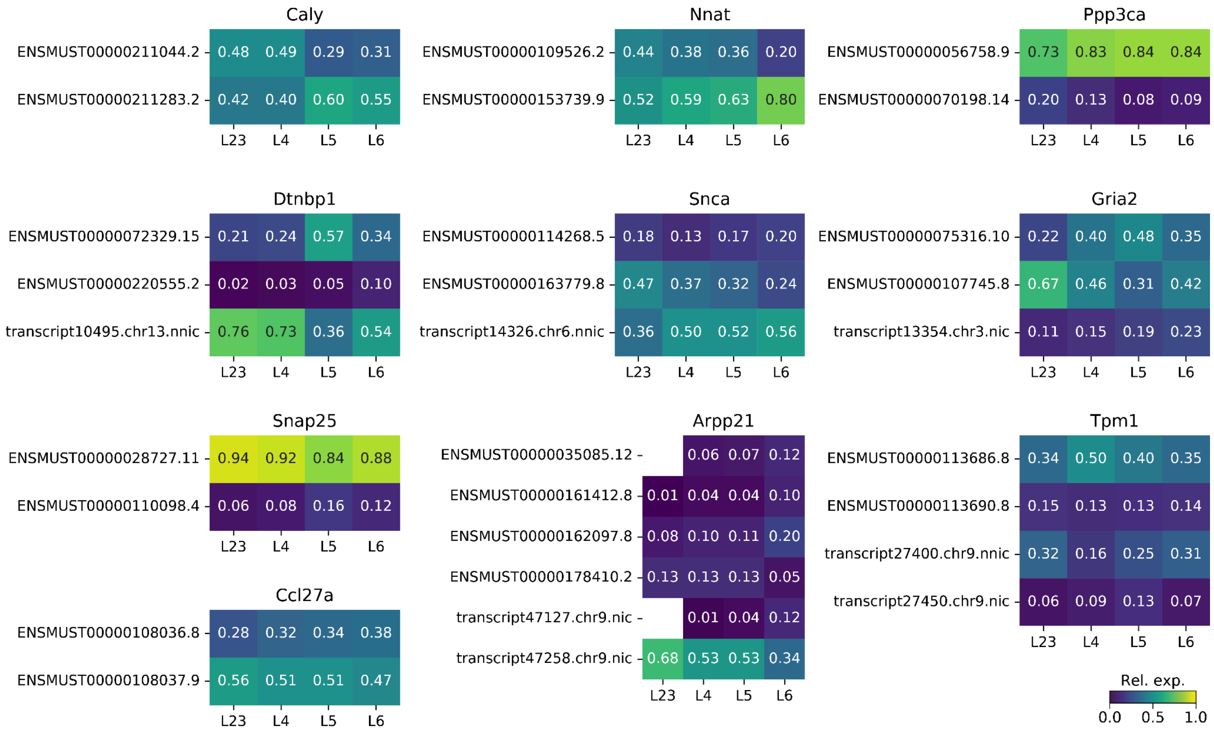

**Figure S8.** For all genes showing layer-specific isoform abundance in excitatory neurons, a heatmap is showing the relative expression of each isoform in the different layers. Only isoforms with relative expression ≥ 0.1 in at least one layer are plotted.

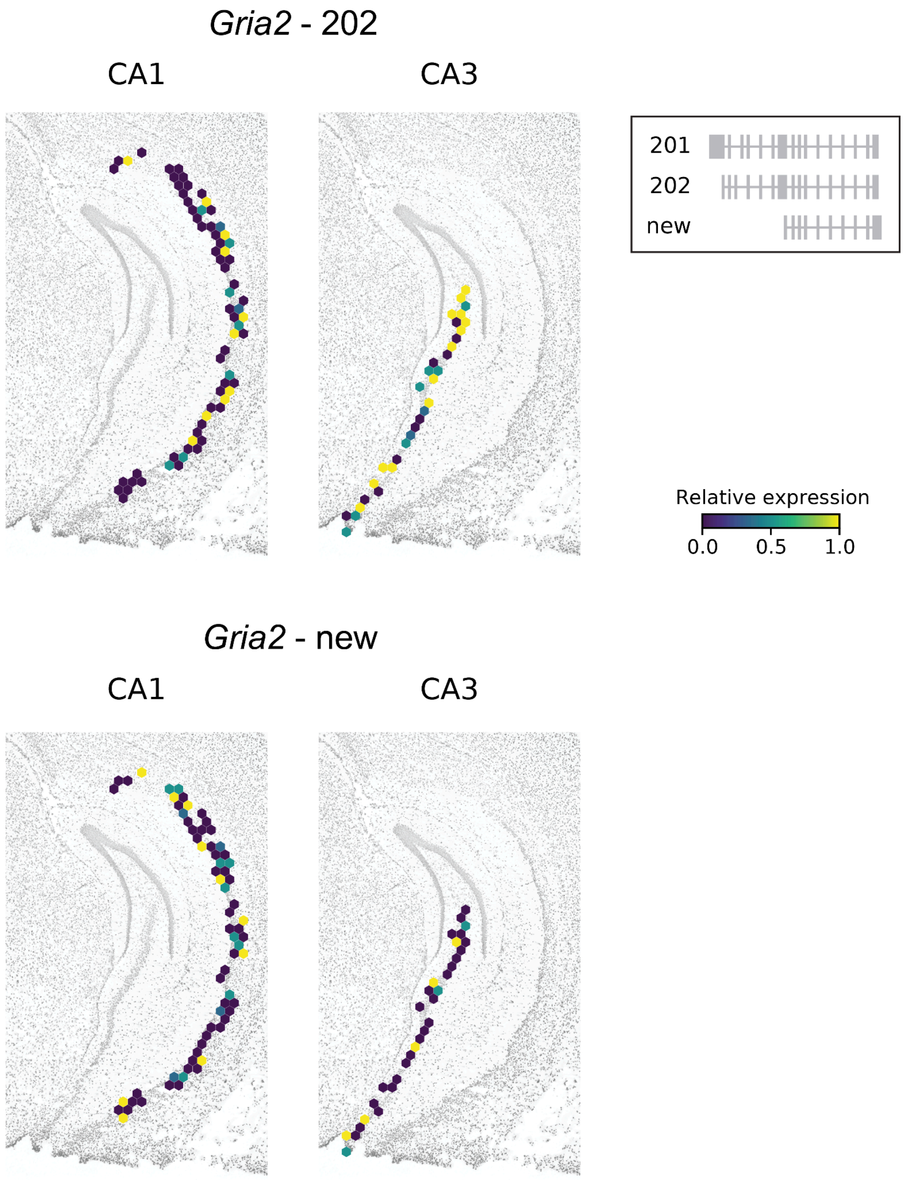

**Figure S9.** Relative expression of *Gria2* isoform 202, and a new transcript in excitatory neurons in different regions of the hippocampus of Sample 1. Every hexagon shows the mean relative expression of the underlying cells (Methods). *Gria2* shows differential isoform changes between CA1 and CA3.

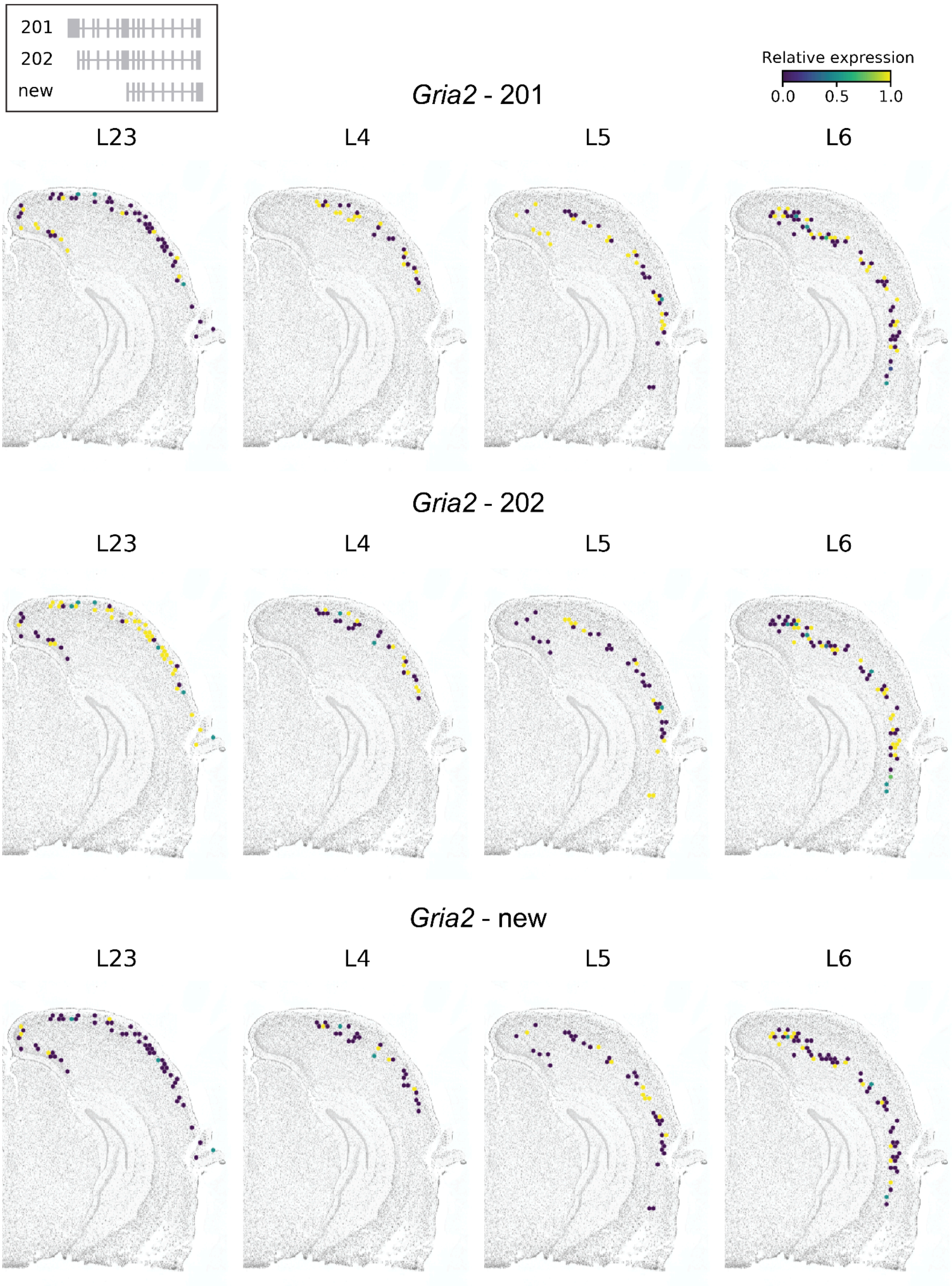

**Figure S10.** Relative expression of *Gria2* isoform 201, 202, and a new transcript in excitatory neurons in different cortical layers of Sample 1. Every hexagon shows the mean relative expression of the underlying cells (Methods). *Gria2* shows differential isoform changes between L23 and L5.

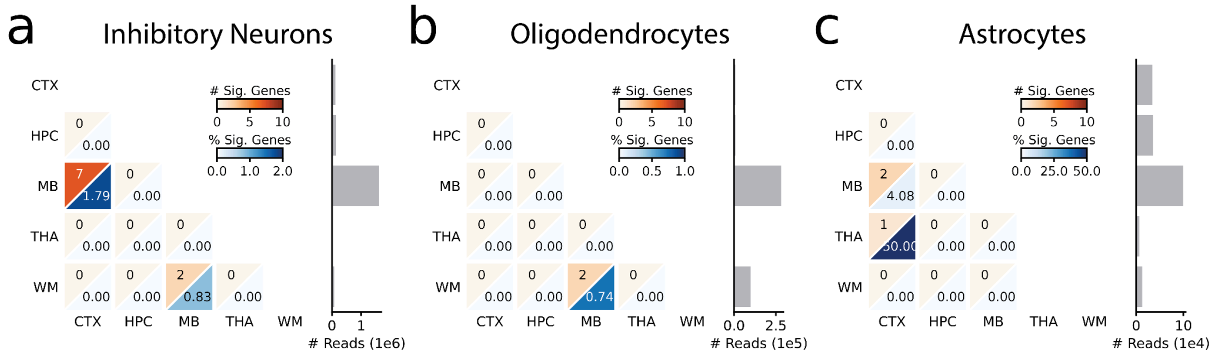

**Figure S11.** Number and percentage of genes changing isoform expression in **a)** inhibitory neurons, **b)** oligodendrocytes, and **c)** astrocytes across brain regions. Barplot on the right shows the number of long reads used during the analysis per brain region. CTX = cortex, HPC = hippocampus, MB = midbrain, THA = thalamus, WM = white matter.

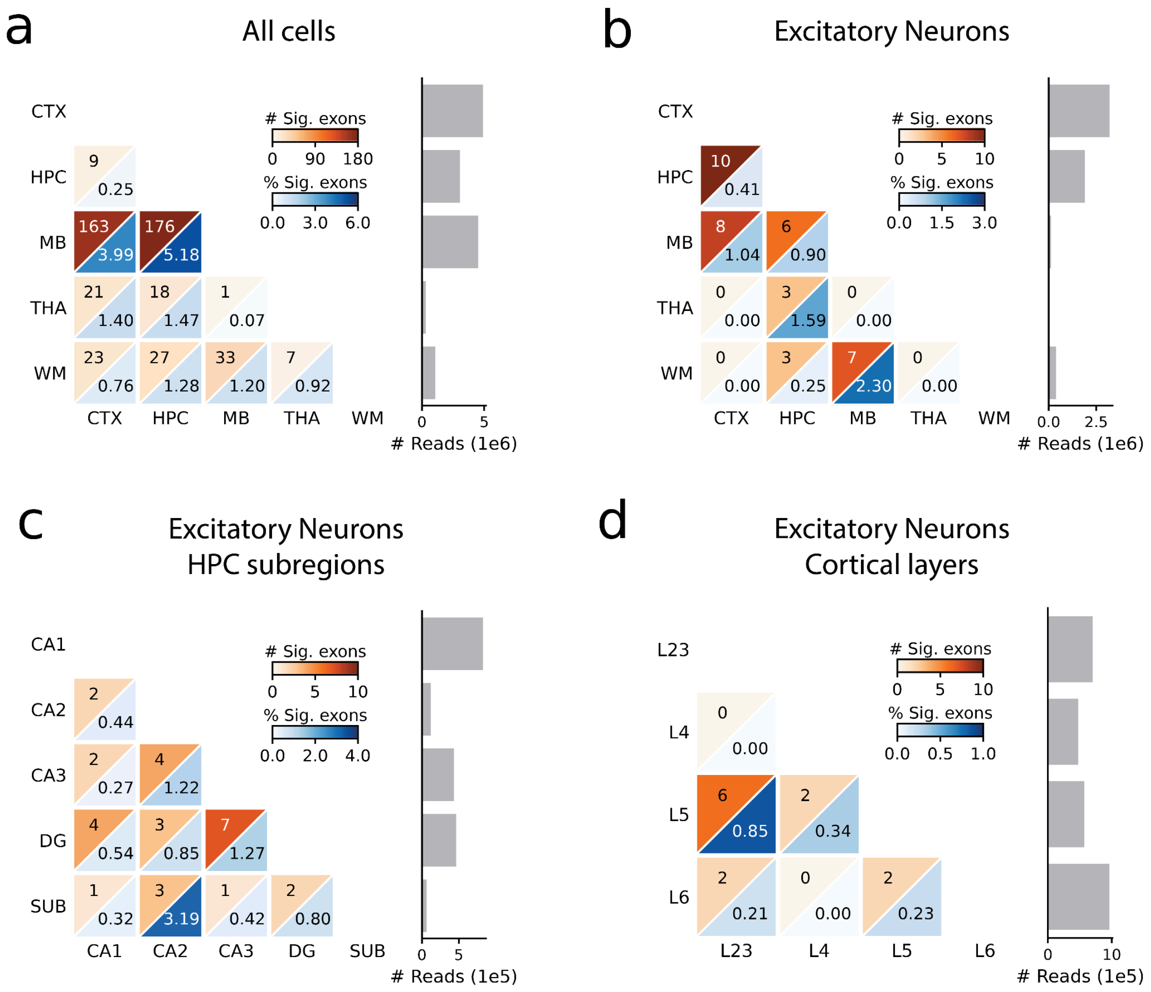

**Figure S12. a)** Number and percentage of exons changing inclusion across predefined brain regions using all cells in combined Sample 1+2 (AE). Barplot on the right shows the number of long reads used during the analysis per brain region. **b-d)** Number and percentage of exons changing inclusion in excitatory neurons across (sub)regions. CTX = cortex, HPC = hippocampus, MB = midbrain, THA = thalamus, WM = white matter, DG = Dentate Gyrus, SUB = Subiculum

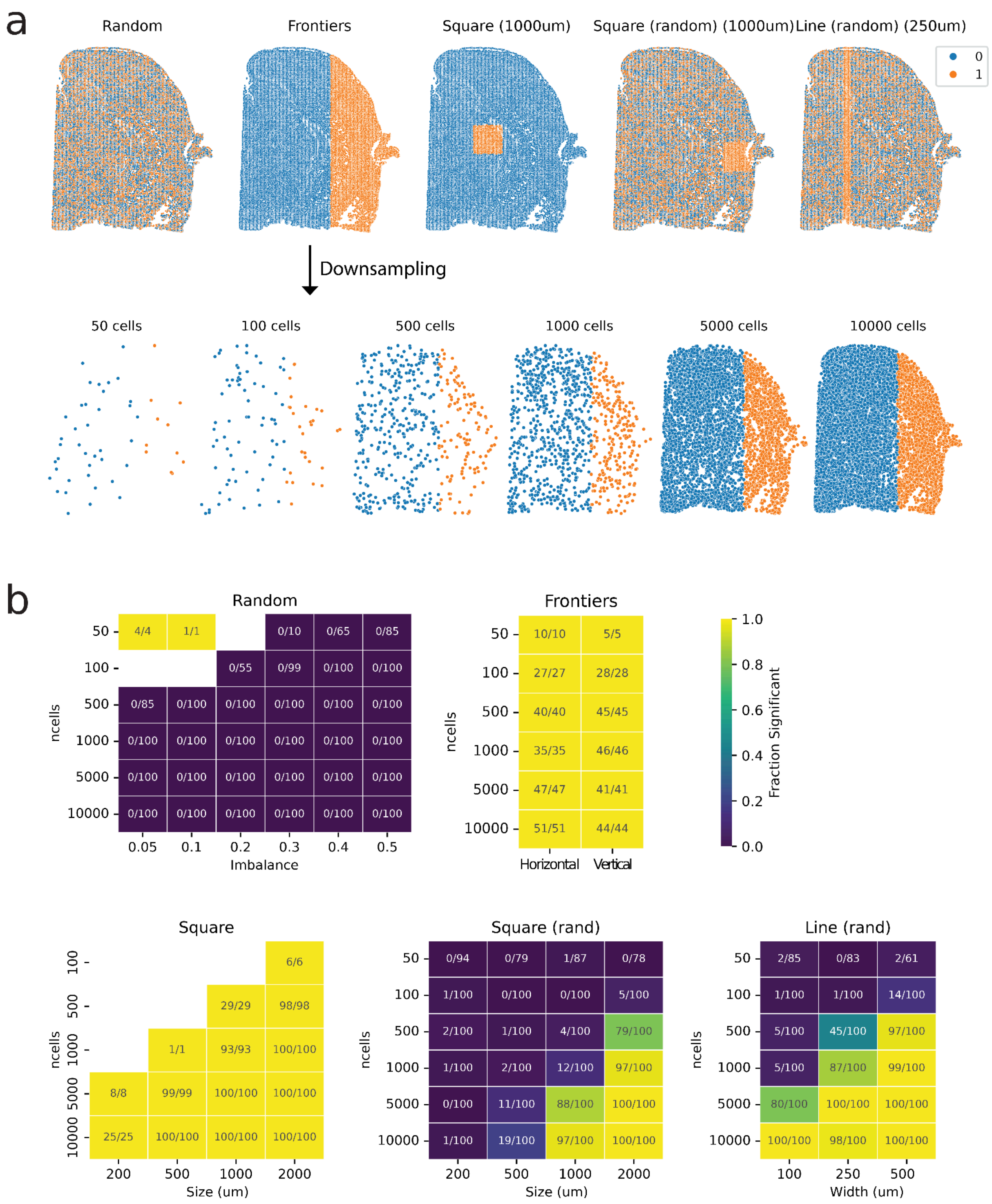

**Figure S13. a)** Example simulation for every category, including a downsampling example for the frontiers simulation. **b)** Heatmaps indicating the number of significant simulations per category. Square simulations with 50 cells never fulfilled the Moran's I testing criteria (see Methods) and are thus missing from the heatmap.

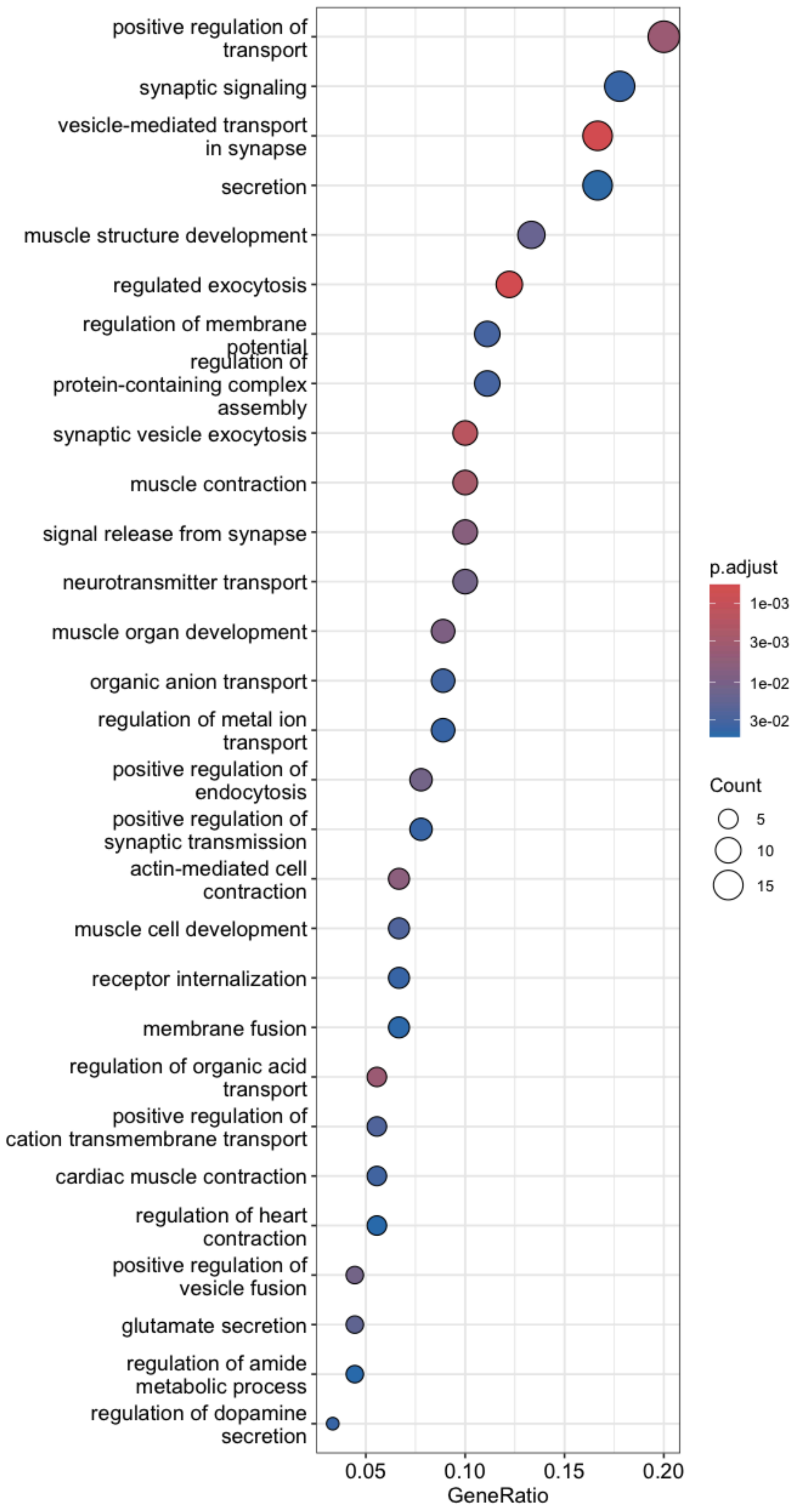

**Figure S14.** Significantly enriched biological processes (BP) in SVIs in the combined Sample 1+2 (AE) identified using GO enrichment analysis.

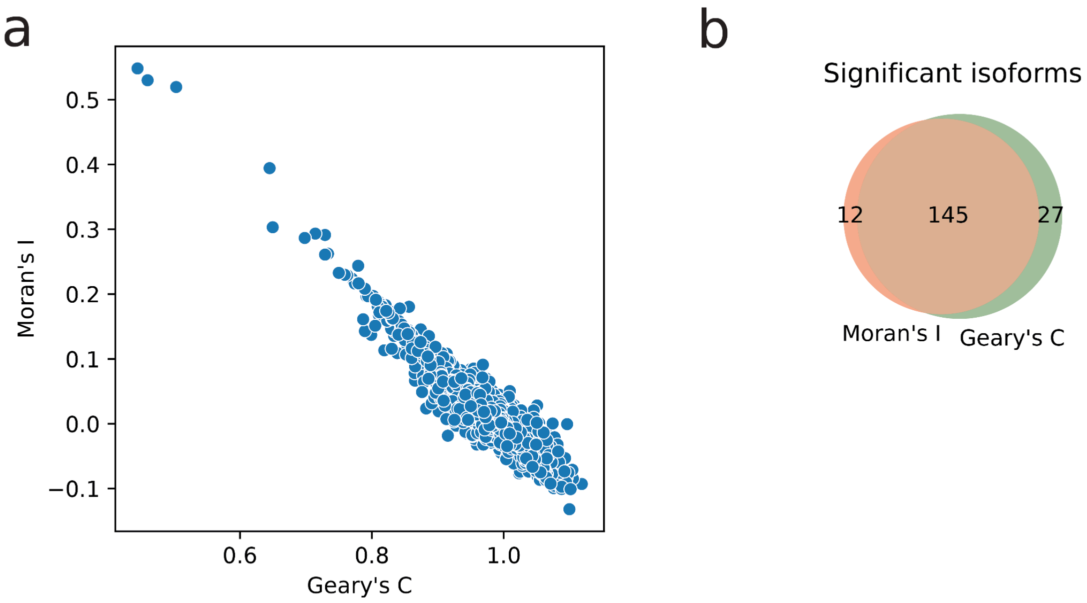

**Figure S15. a)** Geary's C vs. Moran's I score for the tested isoforms in the combined Sample 1+2 (AE) data. **b)** Venn diagram showing the overlap of significant isoforms when using Moran's I or Geary's C.

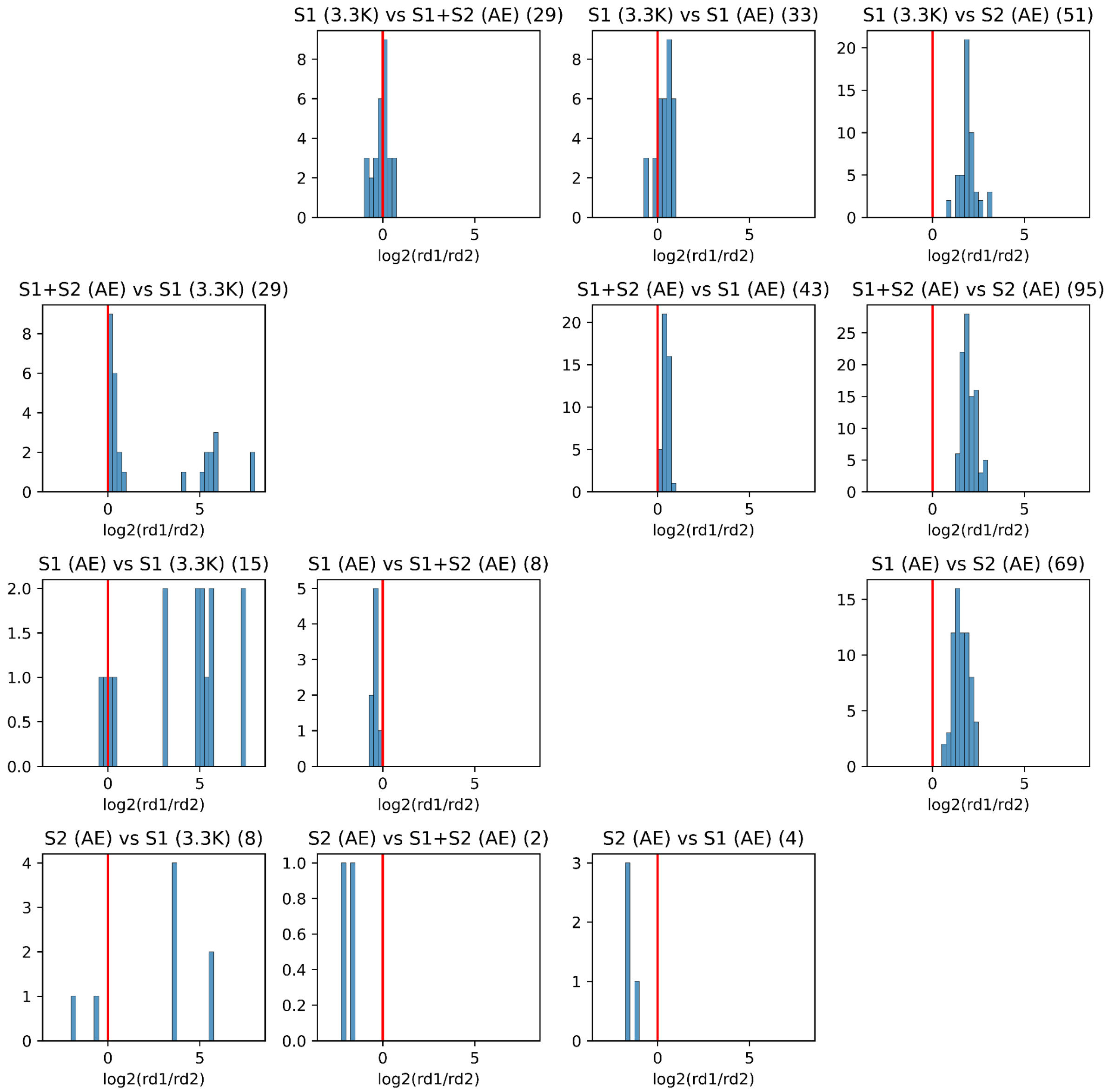

**Figure S16.** For every pairwise combination of datasets (e.g. S1 (3.3K) vs S1+S2 (AE)), we count the number of isoforms significant in the first dataset and not significant in the other. The comparison and number of isoforms are mentioned in the title. For these isoforms, we compare the read depth in the first dataset (rd1) compared to the second dataset (rd2). A value larger than zero indicates that the discordance is potentially explained by the difference in read depth between the samples.

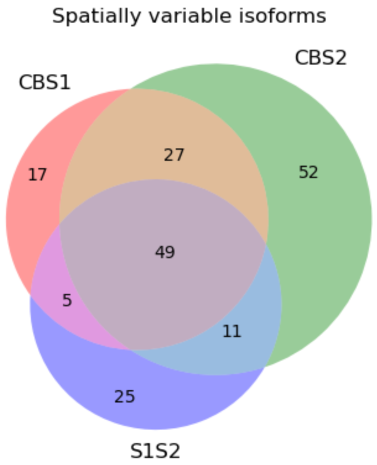

**Figure S17.** Overlap between the detected SVIs in our combined Sample 1+2 (AE) data, CBS1 and CBS2 from Lebrigand et al.

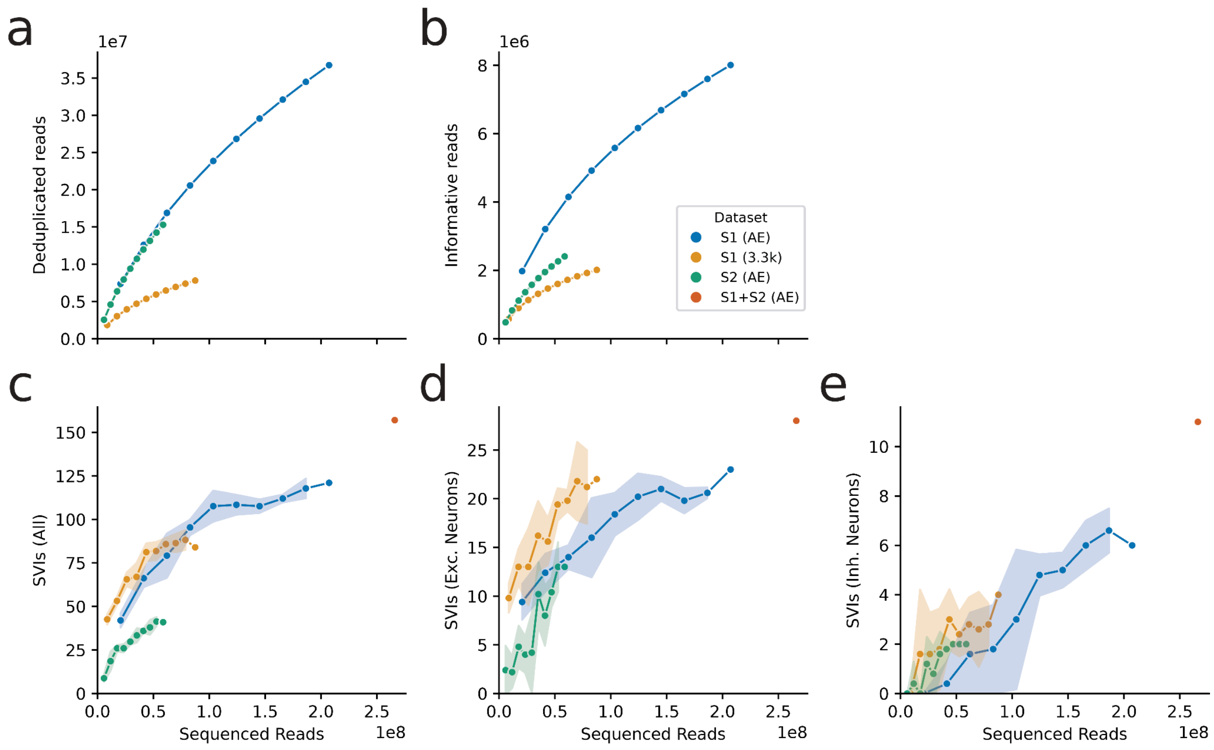

**Figure S18**. Saturation analysis for the Spl-ISO-Seq2 data. The plots show the number of downsampled sequenced reads vs. **a)** deduplicated reads, **b)** informative reads (spliced reads with an assigned isoform that overlap a segmented cell in the correct hemisphere), **c-e)** spatially variable isoforms (SVIs) detected in **c)** all cells, **d)** excitatory neurons, and **e)** inhibitory neurons. Lines show the mean across 10 downsampling experiments; shaded regions indicate ±1 standard deviation.

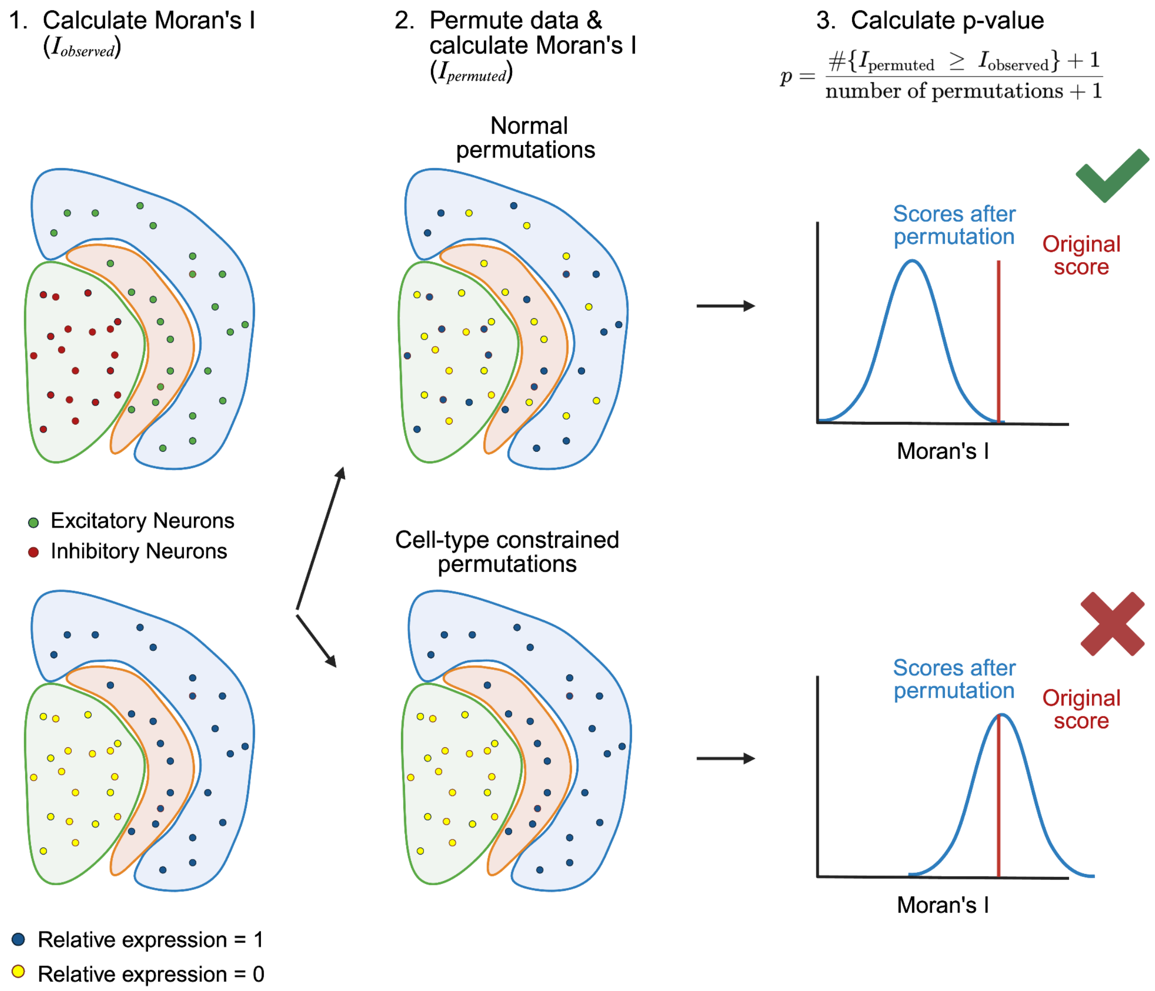

**Figure S19.** Schematic example to explain the difference between normal and cell-type constrained permutations. In the example, the spatial isoform pattern (high relative expression in the blue and orange region vs. low relative expression in the green region) is caused by a different cell-type composition (the blue and orange region only contains excitatory neurons, while the green region contains inhibitory neurons). During our normal permutation tests, the relative expression values of all cells are shuffled, causing the isoform to be significant. However, during the cell-type constrained permutations, only values within a cell type are shuffled and as such this example isoform is not significant anymore.

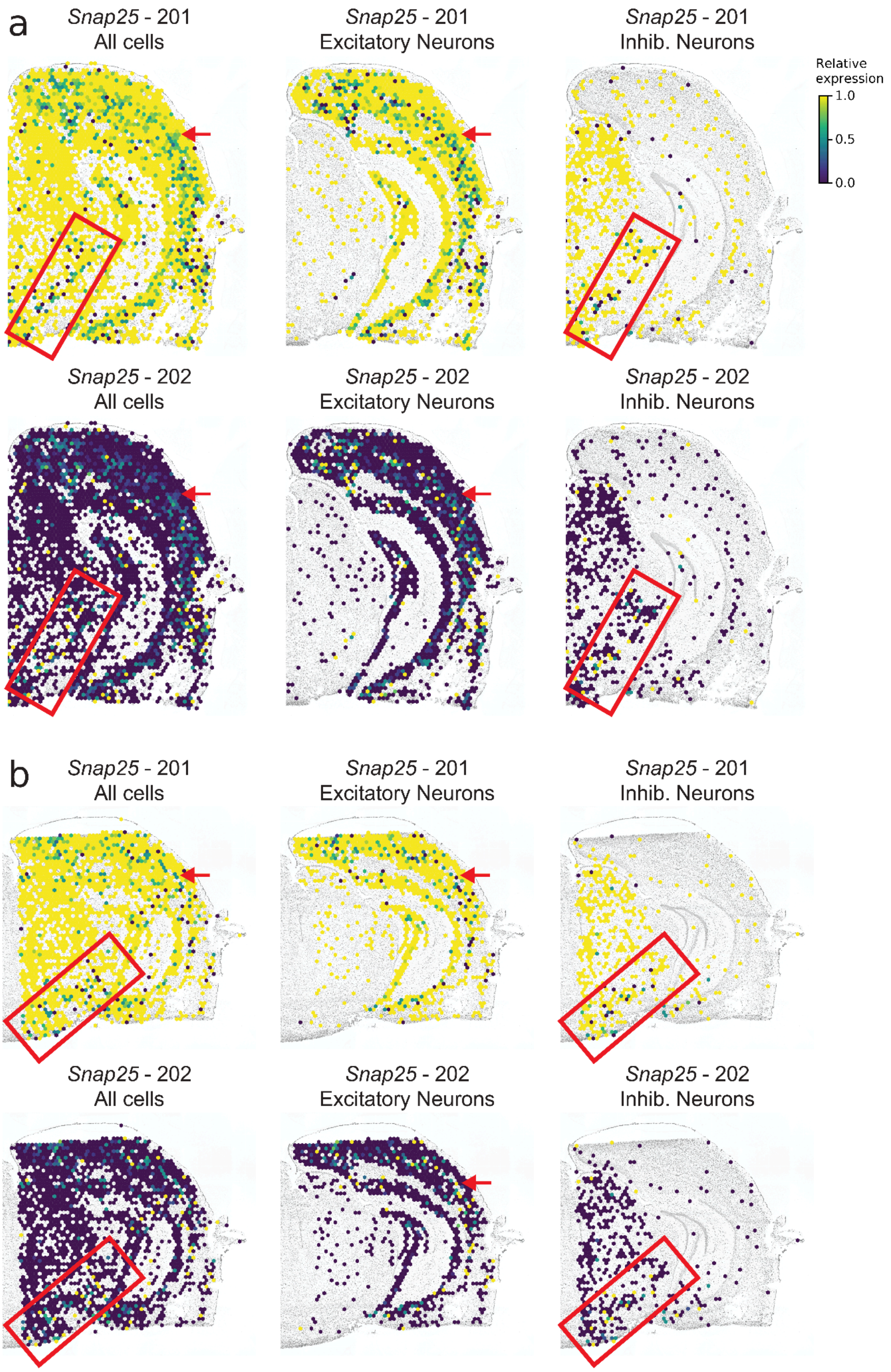

**Figure S20.** Spatial distribution of *Snap25* isoforms in all cells, excitatory and inhibitory neurons in **a)** Sample 1 (AE) and **b)** Sample 2 (AE). Every hexagon shows the mean relative expression of the underlying cells (Methods). Since 201 and 202 are the only isoforms for *Snap25* in our data, the relative expression values add up to 1. The red boxes and arrows indicate decreased expression of isoform 201 and thus increased expression of 202.

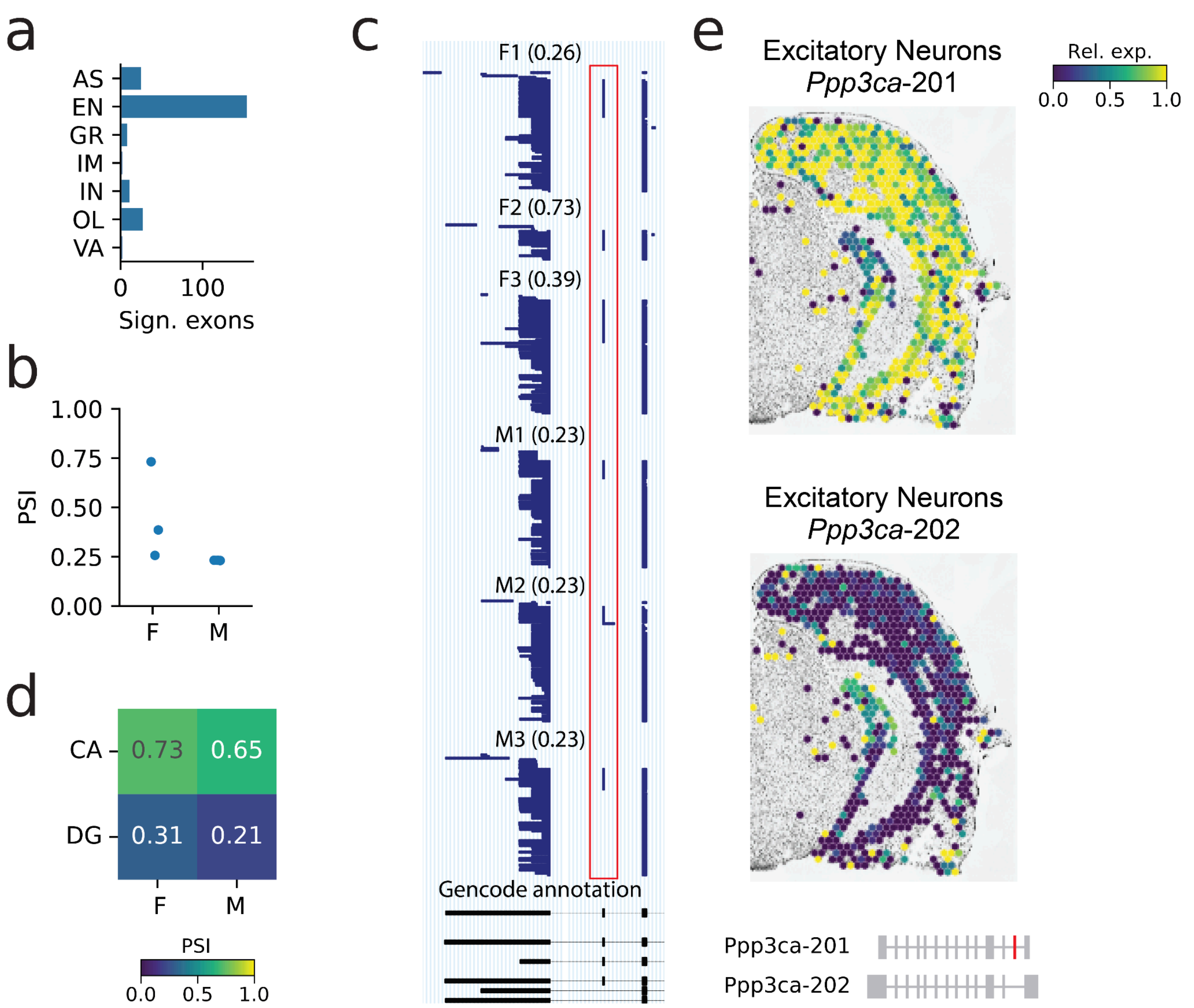

**Figure S21. a)** Number of significant sex-specific exons per cell type. **b)** PSI value for exon chr4_101029166_101029195_- in *PPP3CA* for each individual in excitatory neurons. **c)** Single-cell long-reads for *PPP3CA* measured in excitatory neurons per individual. Each line in the top six tracks represents a single cDNA molecule. The exon, chr4_101029166_101029195_-, in the red box is sex-specific. The bottom black track shows the Gencode annotation. The PSI value for each individual is indicated between brackets. **d)** PSI value for exon chr4_101029166_101029195_- in *PPP3CA* in excitatory neurons in the Dentate Gyrus (DG) and Cornu Ammonis (CA) for females (F) and males (M). **e)** Relative expression of *Ppp3ca*-201 and *Ppp3ca*-202 in excitatory neurons in Sample 1 (AE). Every hexagon shows the mean relative expression of the underlying cells. The ssDNA staining is shown in the background. The schematic diagram in the corner shows the gene structure of *Ppp3ca*-201 and *Ppp3ca*-202, the two most common isoforms in the data. The exon highlighted in red is the main difference between the two isoforms. This is the same exon that shows sex-specific inclusion differences, noting that PPP3CA is encoded on the positive strand in human and on the negative strand in mouse. AS = Astrocytes, EN = Excitatory Neurons, GR = Granule cells, IM = Immune cells, IN = Inhibitory Neurons, OL = Oligodendrocytes, VA = Vascular cells.

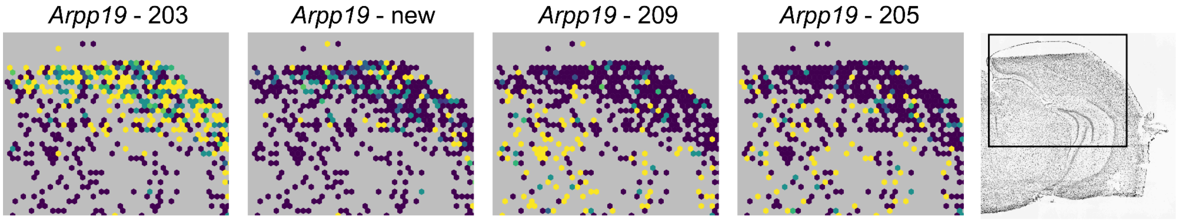

**Figure S22.** Spatial distribution of the four significant isoforms of *Arpp19* in Sample 2 (AE). Every hexagon shows the mean relative expression of the underlying cells (Methods).

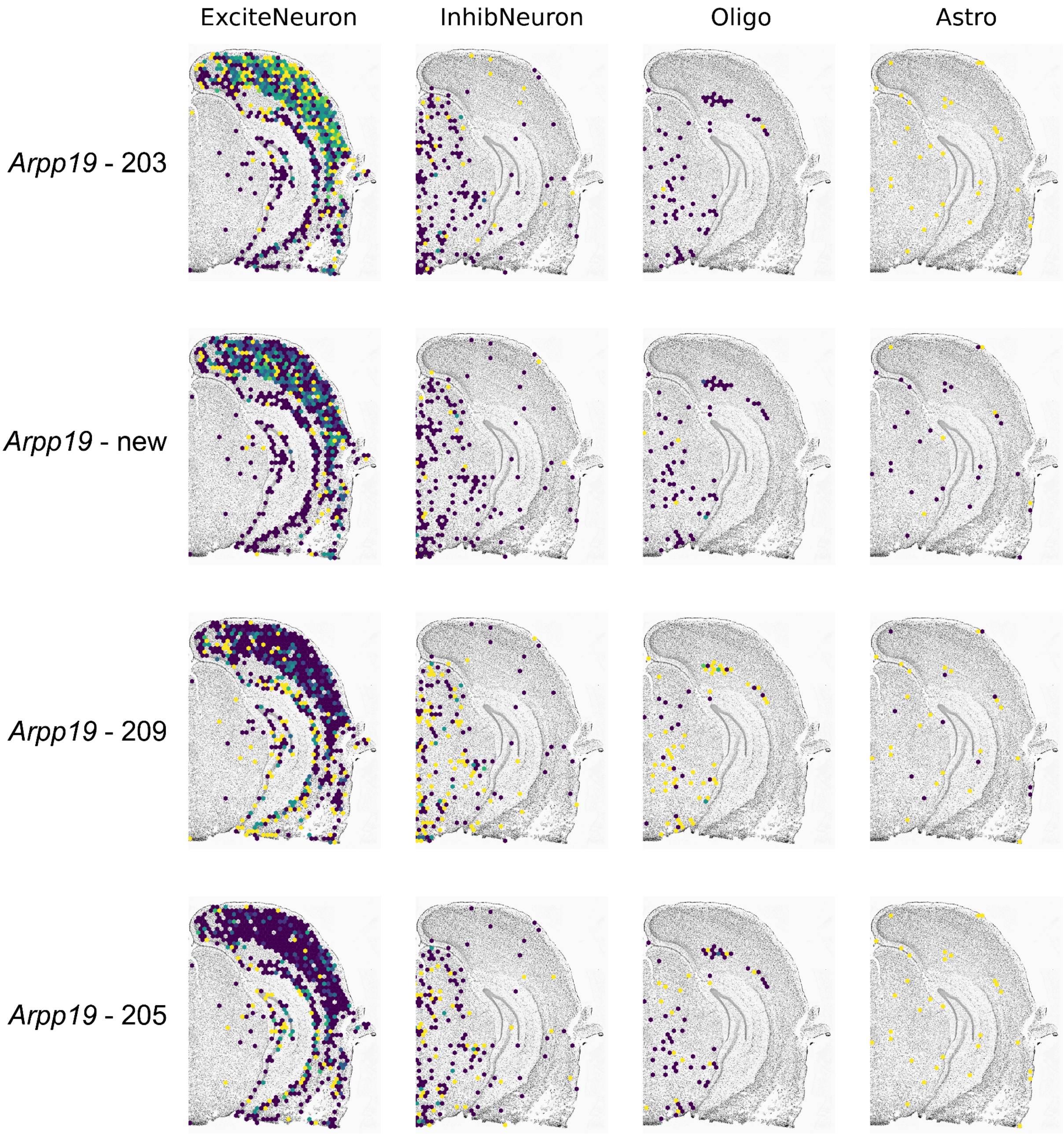

**Figure S23.** Spatial distribution of *Arpp19* isoforms per cell type in Sample 1 (AE). Every hexagon shows the mean relative expression of the underlying cells (Methods).

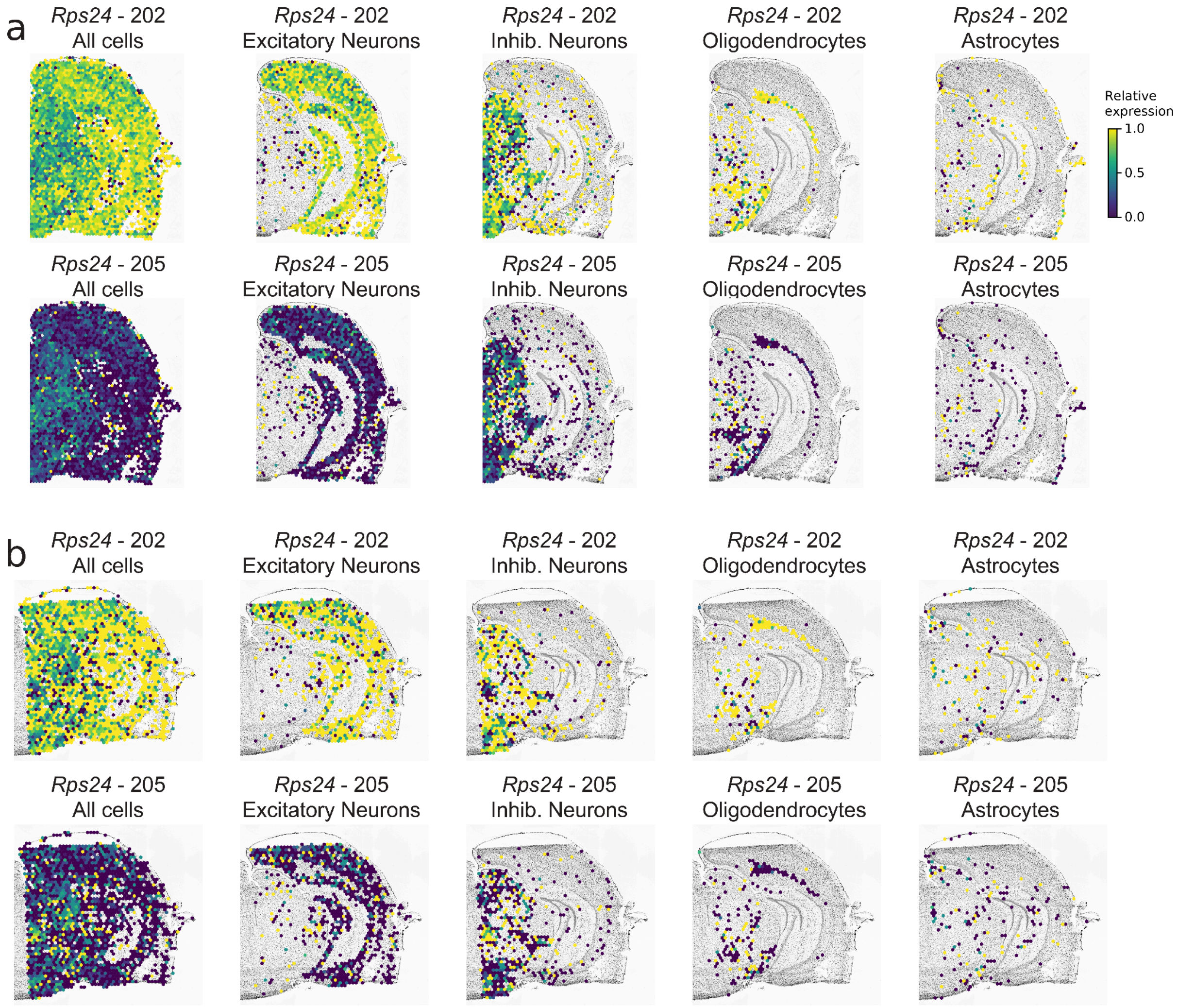

**Figure S24.** Spatial distribution of two isoforms of *Rps24* in all cells and per cell type in **a)** Sample 1 (AE) and **b)** Sample 2 (AE). Every hexagon shows the mean relative expression of the underlying cells (Methods).

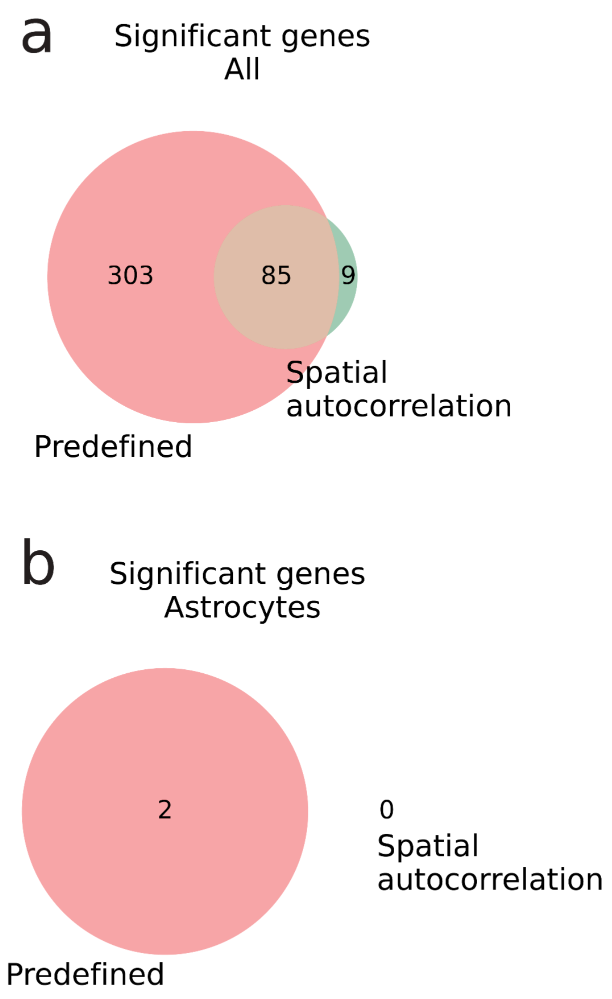

**Figure S25.** Overlap between the significant genes found using both methods for **a)** all cells, **b)** inhibitory neurons, **c)** astrocytes, and **d)** oligodendrocytes in the Sample 1 (AE) + Sample 2 (AE) data.

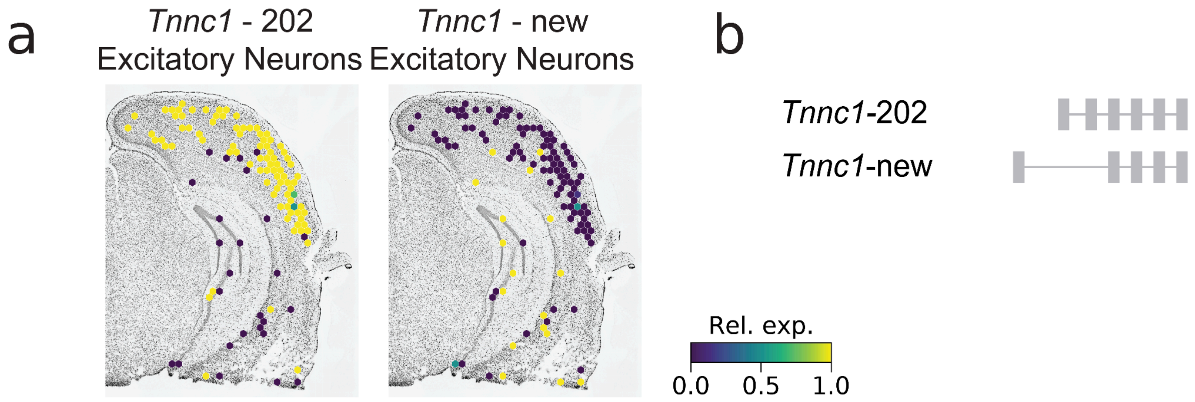

**Figure S26. a)** Spatial distribution of two *Tnnc1* isoforms in excitatory neurons. Every hexagon shows the mean relative expression of the underlying cells (Methods). Isoform 202 is more expressed in the top of the figure, while the novel isoform is more expressed at the bottom of the figure. **b)** Schematic gene diagram of the two isoforms.

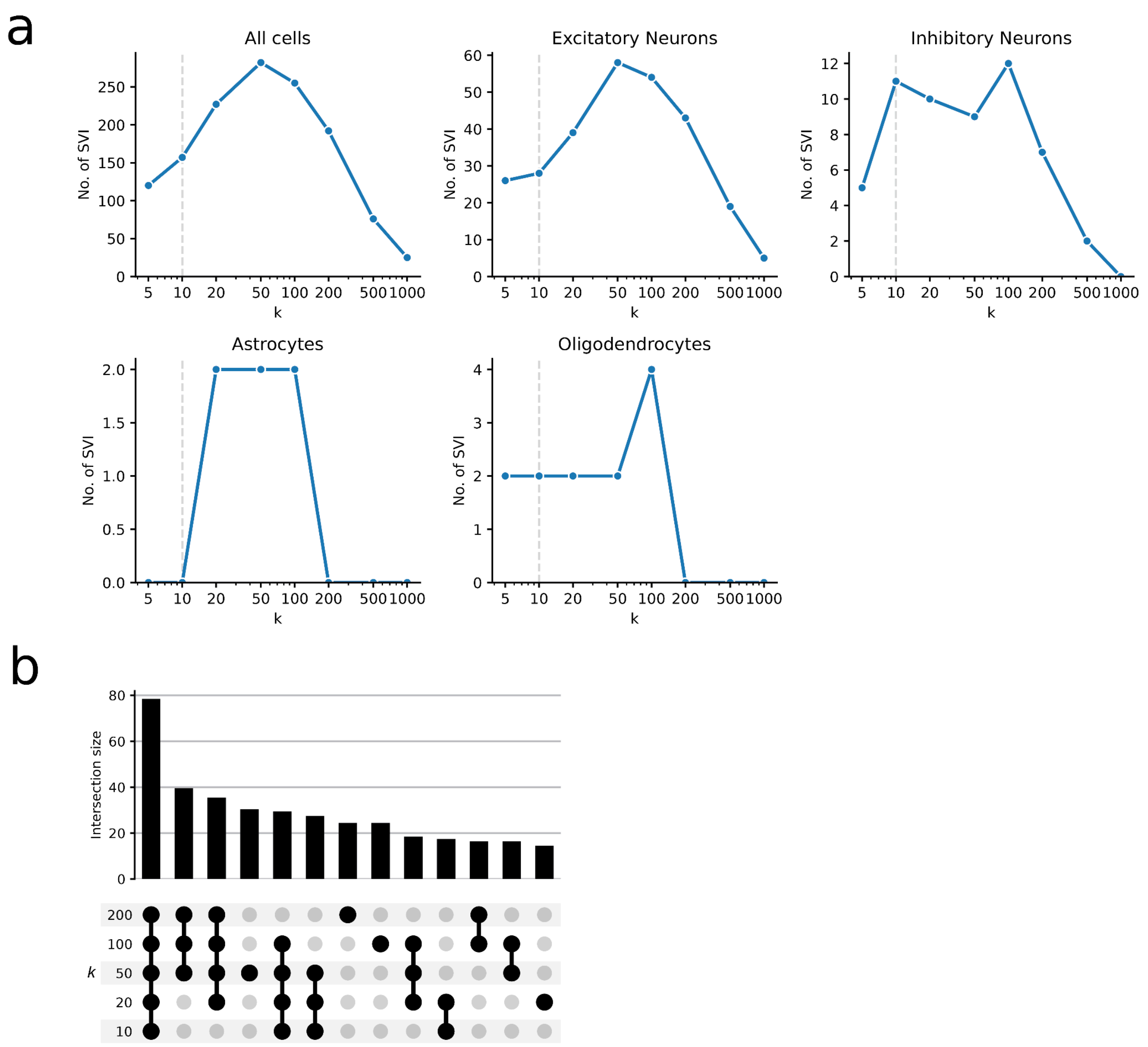

**Figure S27. a)** Number of significant SVIs detected by Spl-IsoFind using different numbers of nearest neighbors (*k*) when using all cells and for each cell type. **b)** The overlap in SVIs when using a different number of neighbors when using all cells.

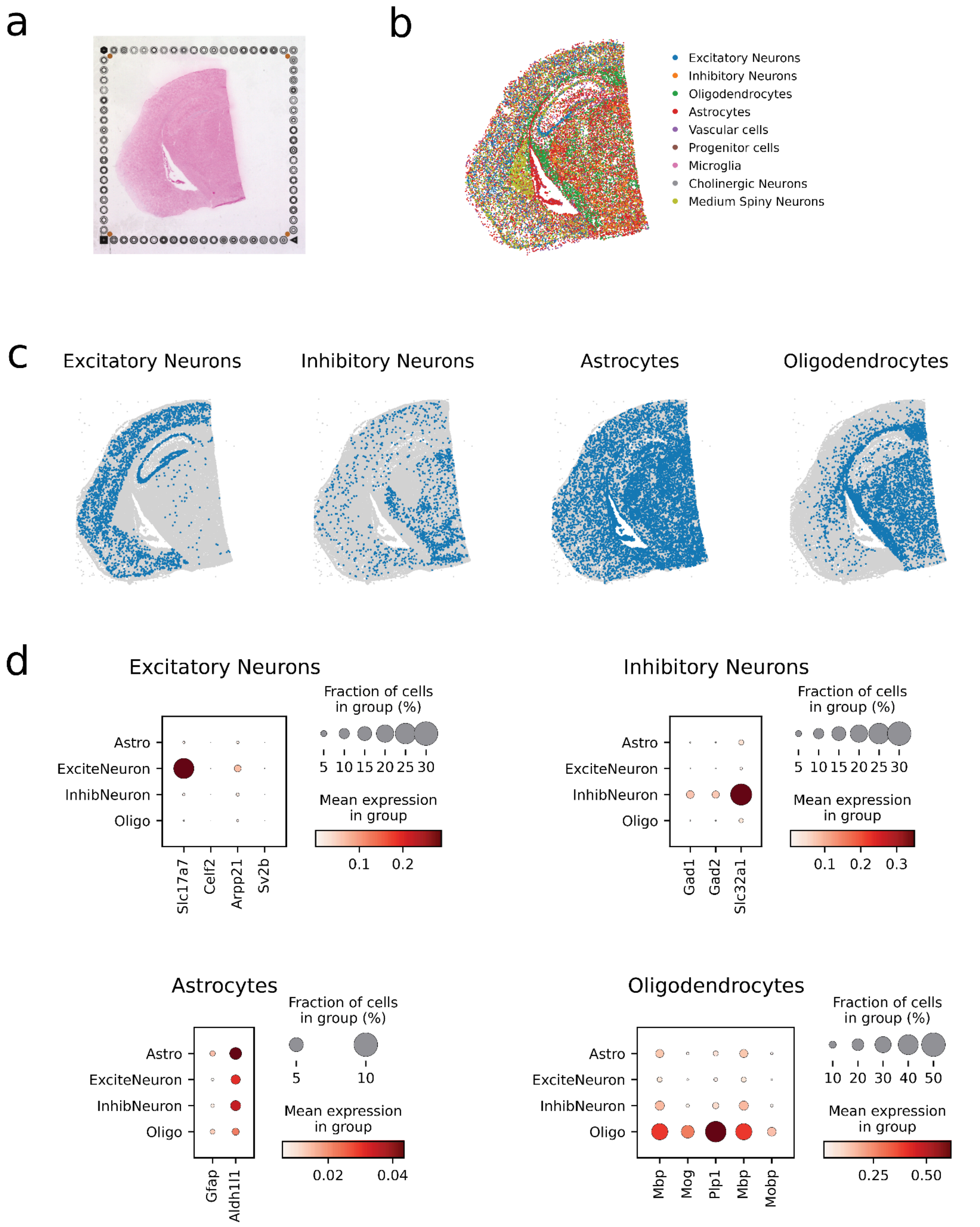

**Figure S28. a)** CytAssist image of the V1 brain slide. **b)** Cell-type annotations of the sample. Only cells annotated as singlets and cell types with >100 cells are plotted. **c)** Cell-type distribution and **d)** marker gene expression of the four major cell types in the Visium HD sample.

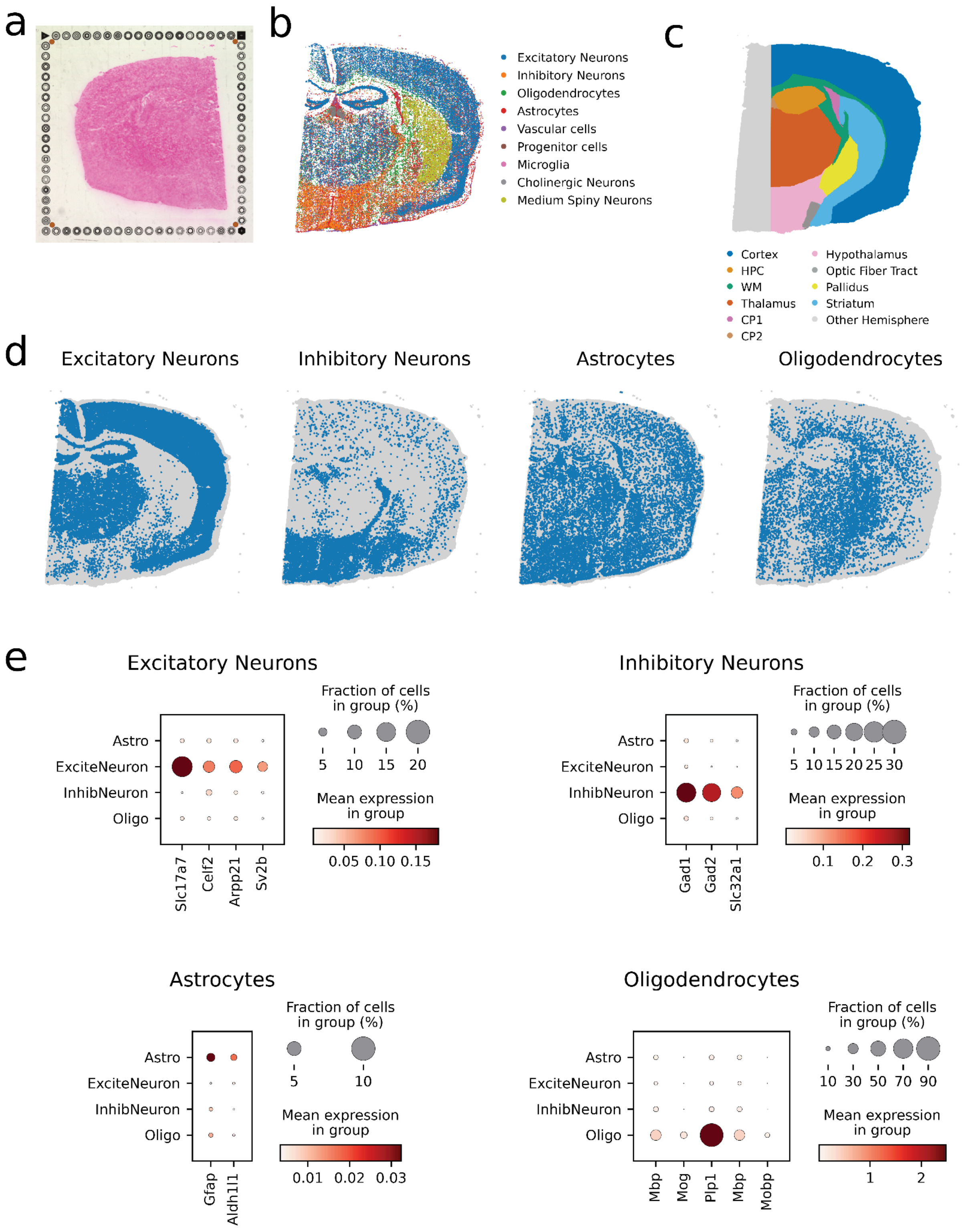

**Figure S29. a)** CytAssist image of the V2 brain slide. **b)** Cell-type annotations of the sample. Only cells annotated as singlets and cell types with >100 cells are plotted. **c)** Brain region annotation. **d)** Cell-type distribution and **e)** marker gene expression of the four major cell types in the Visium HD sample.

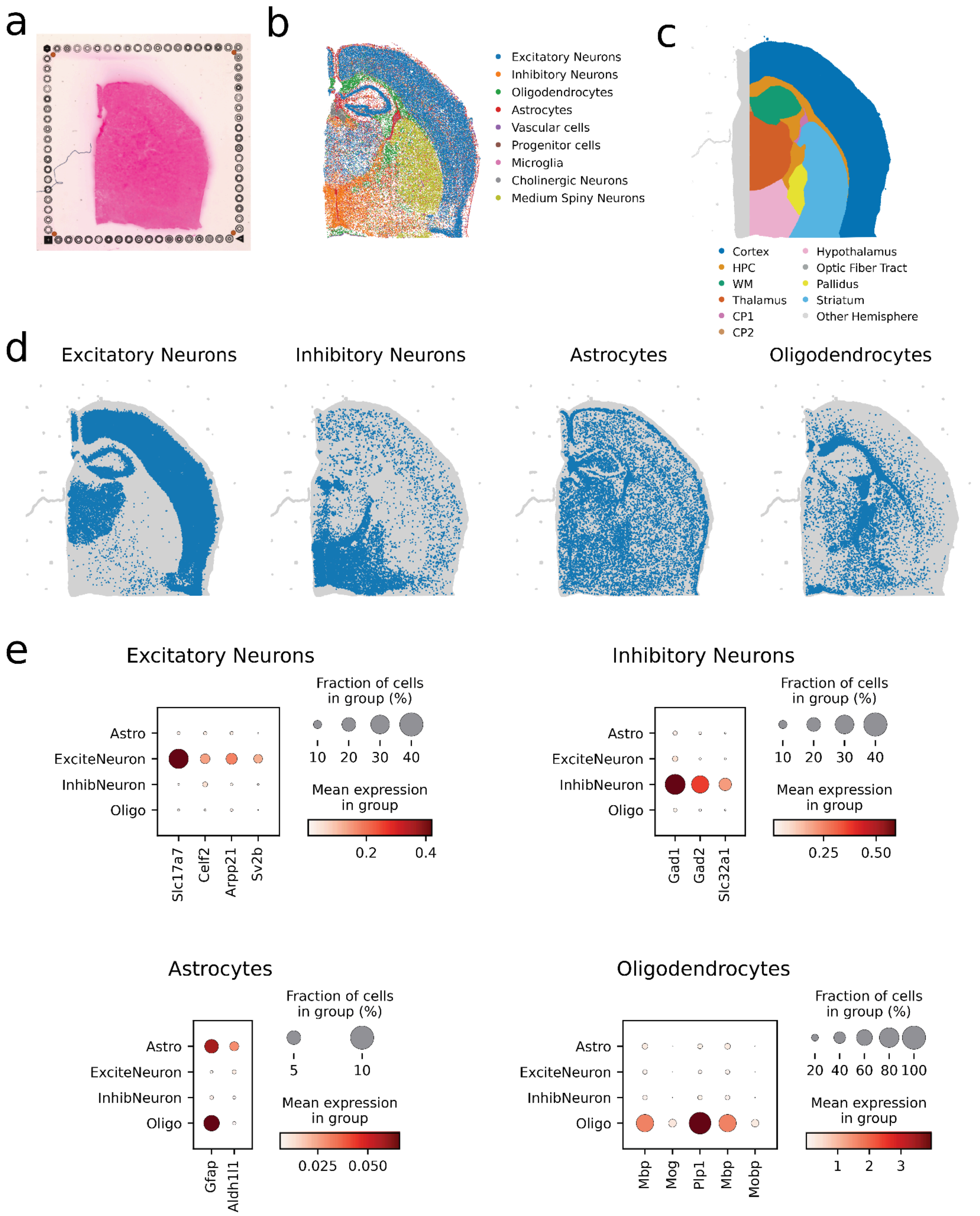

**Figure S30. a)** CytAssist image of the V3 brain slide. **b)** Cell-type annotations of the sample. Only cells annotated as singlets and cell types with >100 cells are plotted. **c)** Brain region annotation. **d)** Cell-type distribution and **e)** marker gene expression of the four major cell types in the Visium HD sample.

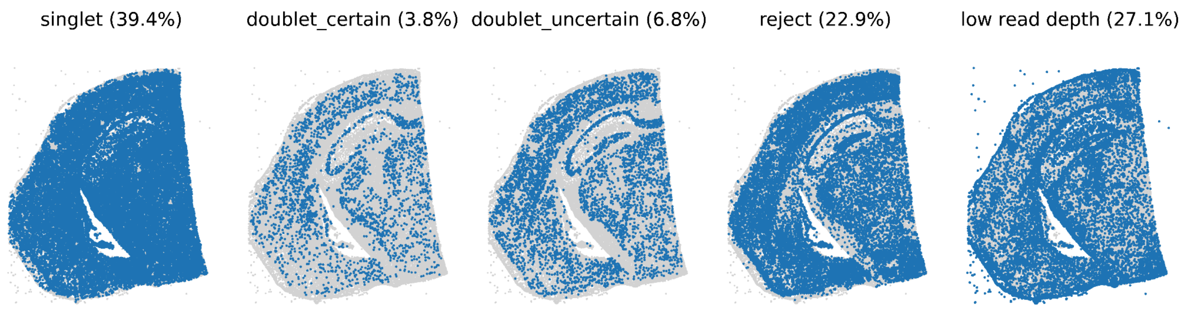

**Figure S31.** RCTD cell-assignment distribution in the V1 sample.

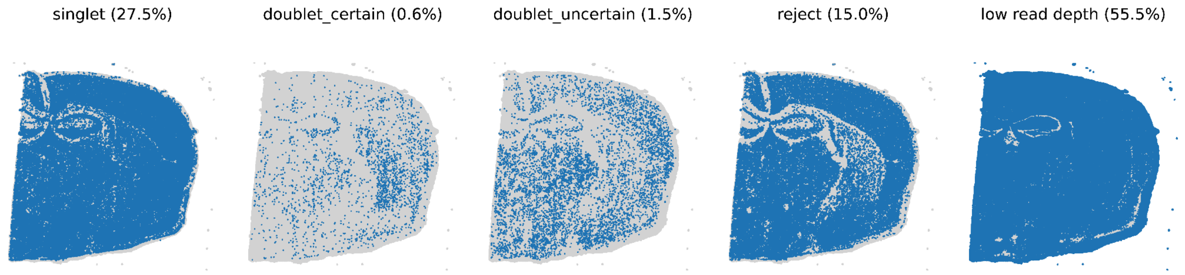

**Figure S32.** RCTD cell-assignment distribution in the V2 sample.

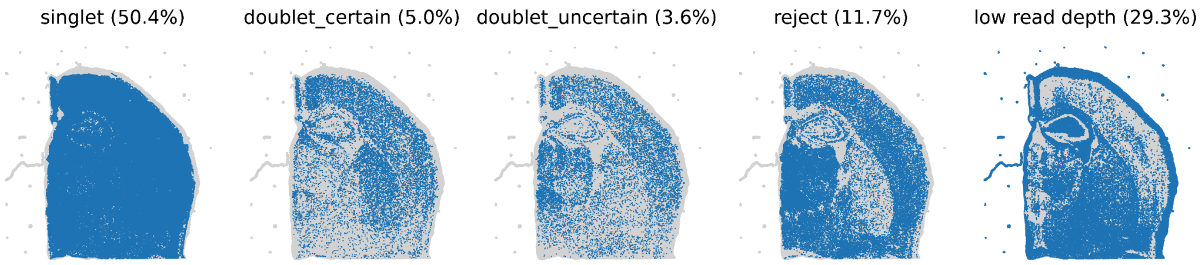

**Figure S33.** RCTD cell-assignment distribution in the V3 sample.

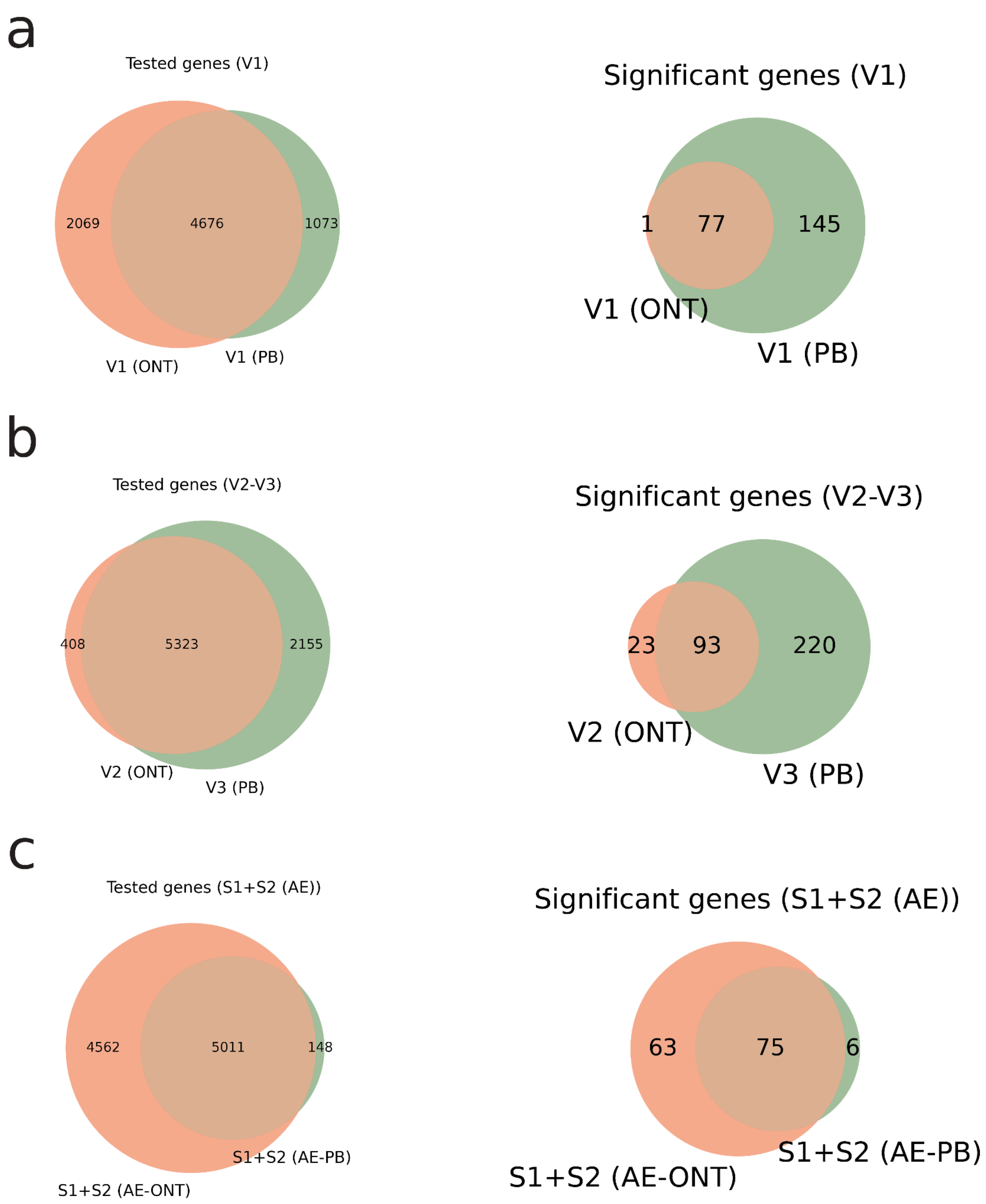

**Figure S34.** Overlap tested and significant genes ONT and PacBio

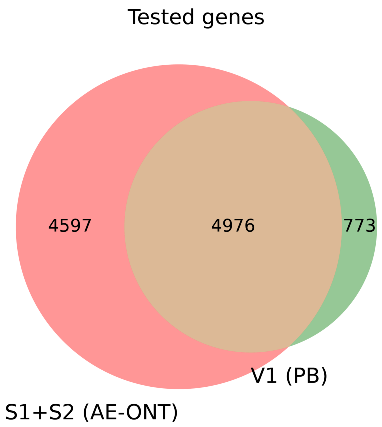

**Figure S35.** Overlap of significant SVIs between Sample1+2 (AE) ONT and Visium HD V1=PB data.

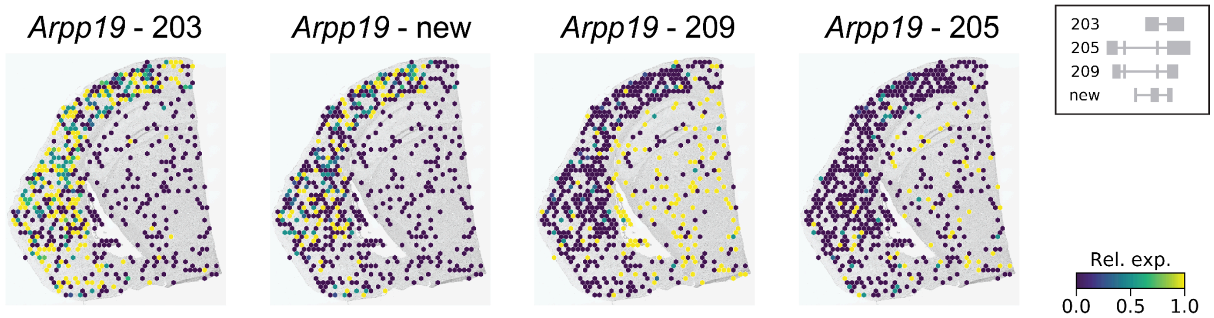

**Figure S36.** Spatial distribution of the four significant isoforms of *Arpp19* in the Visium HD data. Every hexagon shows the mean relative expression of the underlying cells (Methods).

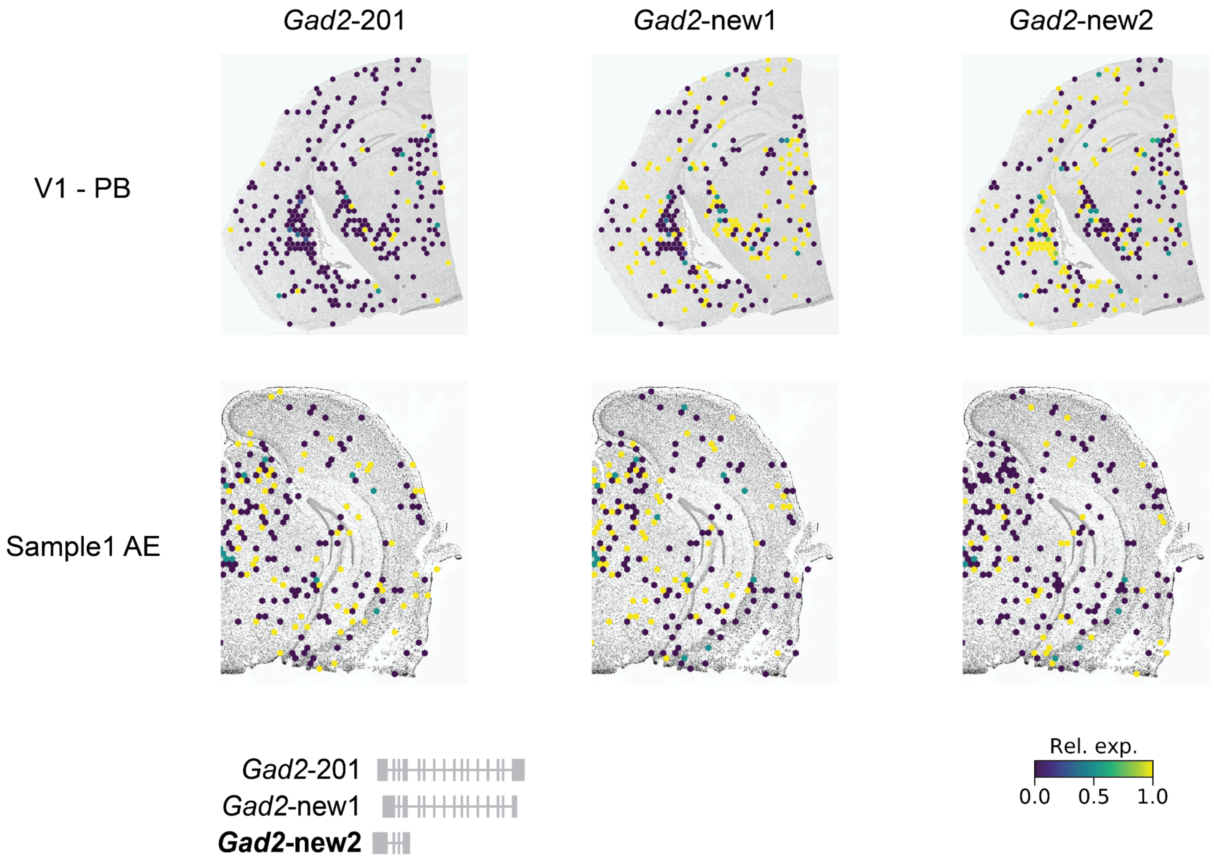

**Figure S37. a)** Spatial distribution of *Gad2* isoforms in V1-PB and Sample1 (AE). Every hexagon shows the mean relative expression of the underlying cells (Methods).

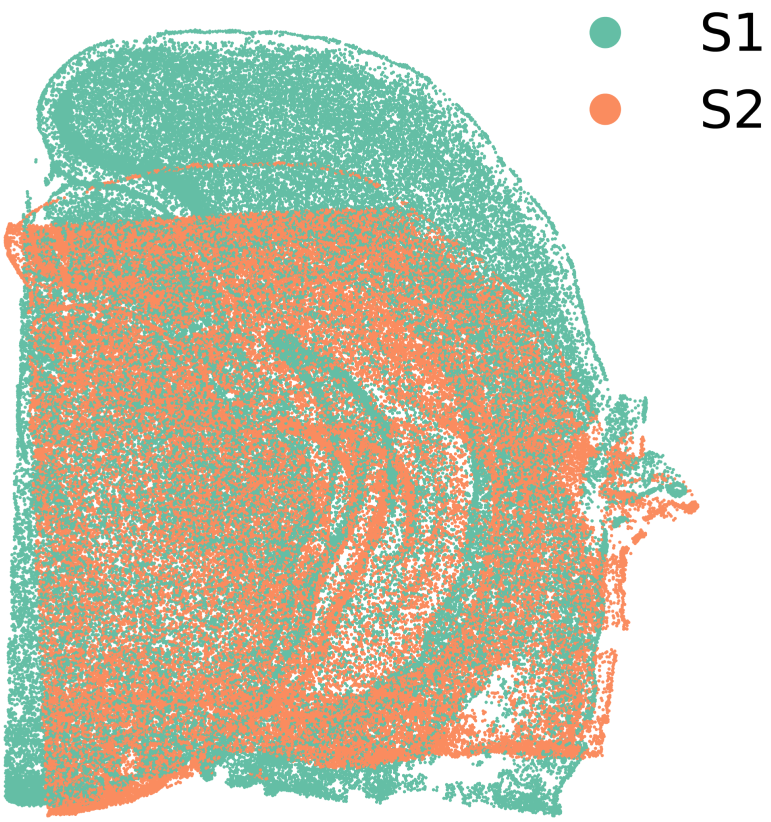

**Figure S38.** Naive alignment of Sample 1 (S1) and Sample 2 (S2)
